# Human gut bacteria produce T_H_17-modulating bile acid metabolites

**DOI:** 10.1101/2021.01.08.425913

**Authors:** Donggi Paik, Lina Yao, Yancong Zhang, Sena Bae, Gabriel D. D’Agostino, Eunha Kim, Eric A. Franzosa, Julian Avila-Pacheco, Jordan E. Bisanz, Christopher K. Rakowski, Hera Vlamakis, Ramnik J. Xavier, Peter J. Turnbaugh, Randy S. Longman, Michael R. Krout, Clary B. Clish, Curtis Huttenhower, Jun R. Huh, A. Sloan Devlin

## Abstract

The microbiota plays a pivotal role in gut immune homeostasis. Bacteria influence the development and function of host immune cells, including T helper cells expressing interleukin-17a (T_H_17 cells). We previously reported that the bile acid metabolite 3-oxolithocholic acid (3-oxoLCA) inhibits T_H_17 cell differentiation^1^. While it was suggested that gut-residing bacteria produce 3-oxoLCA, the identity of such bacteria was unknown. Furthermore, it was not clear whether 3-oxoLCA and other immunomodulatory bile acids are associated with gut inflammatory pathologies in humans. Using a high-throughput screen, we identified human gut bacteria and corresponding enzymes that convert the secondary bile acid lithocholic acid into 3-oxoLCA as well as the abundant gut metabolite isolithocholic acid (isoLCA). Like 3-oxoLCA, isoLCA suppressed T_H_17 differentiation by inhibiting RORγt (retinoic acid receptor-related orphan nuclear receptor γt), a key T_H_17 cell-promoting transcription factor. Levels of both 3-oxoLCA and isoLCA and the 3α-hydroxysteroid dehydrogenase (3α-HSDH) genes required for their biosynthesis were significantly reduced in patients with inflammatory bowel diseases (IBD). Moreover, levels of these bile acids were inversely correlated with expression of T_H_17 cell-associated genes. Overall, our data suggest that bacterially produced T_H_17 cell-inhibitory bile acids may reduce the risk of autoimmune and inflammatory disorders such as IBD.

## Introduction

Bile acids are steroidal natural products that are synthesized from cholesterol in the liver and secreted into the gut postprandially, where they act as detergents that aid in digestion^2,3^. Host-derived primary bile acids that escape enterohepatic recirculation enter the large intestine, forming a concentrated pool of metabolites (200-1000 μM)^4,5^. In the gut, these compounds are metabolized by resident microbes to form a large group of compounds called secondary bile acids^3^. Both primary and secondary bile acids regulate host metabolism, including energy expenditure, glucose and lipid homeostasis^2,6^, and host immune responses, including both innate and adaptive immunity^7–10^.

Bile acids modulate the differentiation and function of T cells, including inflammatory T helper 17 (T_H_17) cells and anti-inflammatory regulatory T (T_reg_) cells, which play critical roles in the protection against extracellular pathogens and the maintenance of host immune tolerance, respectively^11–15^. Specifically, either combinations of primary and secondary bile acids^16^, or the secondary bile acids isoallolithocholic acid (isoalloLCA) and isodeoxycholic acid (isoDCA)^1,17^ modulate differentiation of T_reg_ cells. In addition, 3-oxoLCA, another lithocholic acid derivative, inhibits T_H_17 cell differentiation by blocking the function of the nuclear hormone receptor (NhR) RORγt^1^, which, along with RORα, orchestrates the T_H_17 cell program^18,19^. 3-oxoLCA is absent from the ceca of germ free (GF) B6 mice^1^, indicating that the gut microbiota is necessary for production of this metabolite and suggesting that gut bacteria may directly synthesize 3-oxoLCA. However, it is unknown which commensal bacteria and bacterial enzymes produce 3-oxoLCA (**Fig. 1a**) and whether this compound is relevant to the pathophysiology of major GI tract inflammatory disorders. Moreover, it is likely that there are additional secondary bile acids that modulate T_H_17 cell responses and may be implicated in the pathogenesis of IBD.

**Fig. 1.**
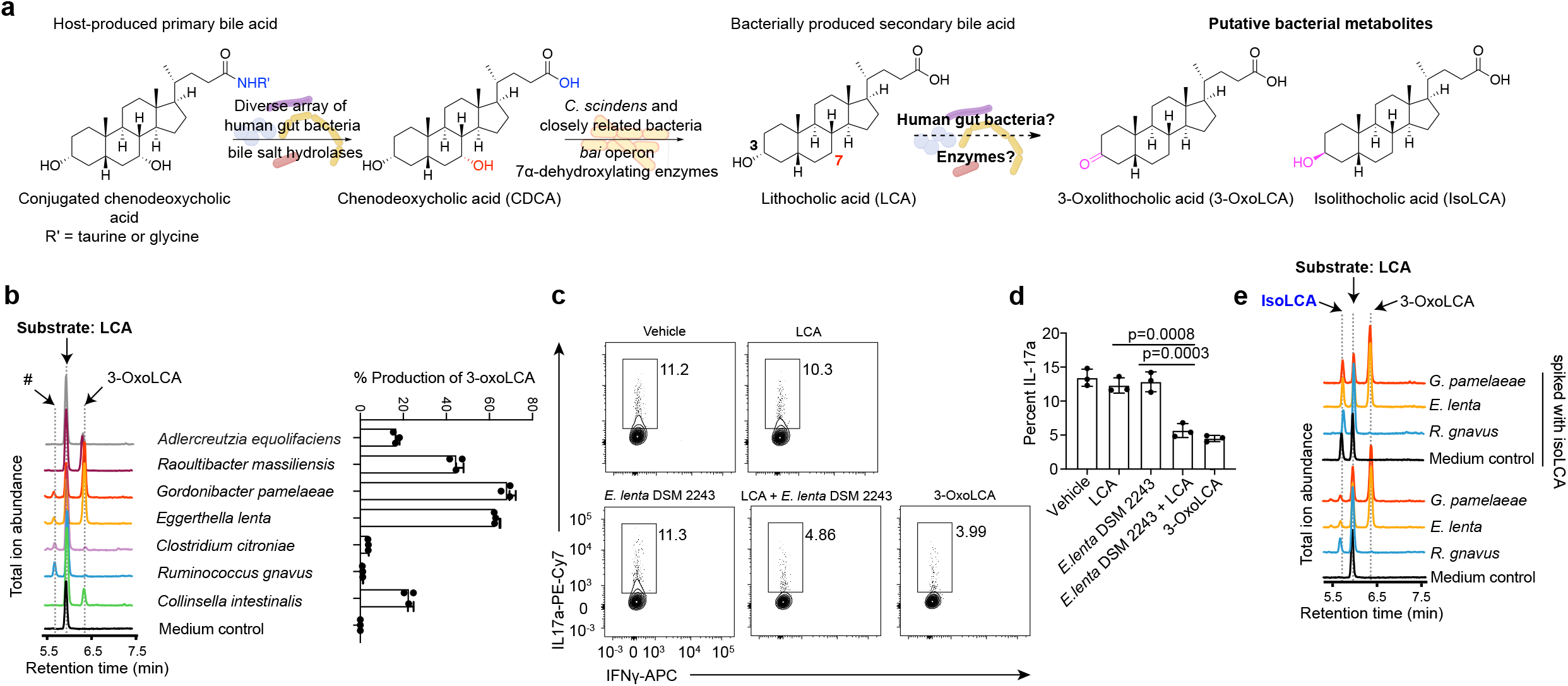
Screening of human stool samples identifies bacterial producers of 3-oxoLCA, a T_H_17-modulating bile acid metabolite. **a**, Human gut bacteria convert host-produced conjugated chenodeoxycholic acid (CDCA) to lithocholic acid (LCA) through the sequential actions of bile salt hydrolase (BSH) and bile acid-inducible (*bai*) operon enzymes. While it has been proposed that gut bacteria convert LCA to the metabolites 3-oxolithocholic acid (3-oxoLCA) and isolithocholic acid (isoLCA), the bacterial strains and enzymes responsible for this conversion were not known. **b**, Representative UPLC-MS traces (left) and percent production of the target molecule 3-oxoLCA (right) by bacteria from the human isolate screen. An unknown peak of m/z 375.2 (#, retention time 5.7 min), was later identified as isolithocholic acid (isoLCA) (see **Fig. 1e**) (n = 3 biological replicates per group). See **Table S2** for full results. **c, d**, Supernatants from *E. lenta* DSM 2243 cultured with LCA inhibited T_H_17 cell differentiation in vitro. Representative FACS plots (**c**) and population frequencies of mouse T_H_17 cells (**d**) activated and expanded in vitro are shown. Mouse naive CD4+ T cells from wild-type B6Jax mice were cultured under T_H_17 cell polarizing conditions for 3 days and bacterial supernatants were added 18 hours after T cell receptor (TCR) activation (n = 3 biologically independent samples per group, data are mean ± SEM, followed by one-way ANOVA with one-way Tukey’s multiple comparison test). **e**, A pure standard of isoLCA was spiked into a subset of bacterial culture extracts containing the new peak (#). Co-elution and an identical m/z match confirmed that the new compound (#) in **Fig. 1b** was isoLCA.

Here, using a high-throughput screen of human isolates, we identify gut bacteria that produce 3-oxoLCA as well as an abundant gut metabolite, isolithocholic acid (isoLCA), which we demonstrate inhibits T_H_17 cell differentiation. Multi-omics analyses of two IBD registries revealed that 3-oxoLCA and isoLCA as well as bacterial genes responsible for the production of these metabolites were negatively associated with IBD and T_H_17/IL-17a related host gene expression. Together, our data suggest that bacterial production of 3-oxoLCA and isoLCA may contribute to gut immune homeostasis in humans.

## Results

### Screen for 3-oxoLCA-producing gut bacteria that inhibit T_H_17 cell differentiation

To identify human gut bacteria that produce 3-oxoLCA, we screened strains isolated from human stool for their ability to convert LCA into this immunomodulatory bile acid. We chose LCA as a substrate because this secondary bile acid is present in high concentrations in human cecal contents (mean ∼160 μM)^5^ and because we reasoned that a variety of gut bacteria might be able to oxidize the C3-hydroxyl group of LCA to produce 3-oxoLCA (**Fig. 1a**). First, we quantified the levels of 3-oxoLCA in stool samples from 15 individuals by ultra-high performance liquid chromatography-mass spectrometry (UPLC-MS) and selected the two stool samples that contained the highest 3-oxoLCA levels to use in the screen (p#3 and p#27, **Extended Data Fig. 1a**). We established a library of 990 culturable isolates using anaerobic culture of serially diluted stool samples (**Extended Data Fig. 1b**). 16S rRNA gene sequencing from pooled cultures isolated from each donor’s stool revealed that the cultivable colonies represented a diverse array of bacteria, with members from all of the major gut phyla represented (**Extended Data Fig. 1c** and **d**, see Methods for details).

A total of 238 bacterial isolates were found with 1% or higher conversion of LCA to 3-oxoLCA (24% positive hit rate) after 48-hour incubation. Full-length 16S rRNA gene sequencing revealed that the identified isolates belonged to 12 bacterial genera (*Adlercreutzia, Bifidobacterium, Enterocloster, Clostridium, Collinsella, Eggerthella, Gordonibacter, Monoglobus, Peptoniphilus, Phocea, Raoultibacter, Mediterraneibacter*) (**Table S2**). Among these, the top producers included *Gordonibacter pamelaeae* P7-E3 (mean 68%), *Eggerthella lenta* P7-G7 (mean 63%), *Raoultibacter massiliensis* P7-A2 (mean 44%), *Collinsella intestinalis* P8-C12 (mean 23%), *Adlercreutzia equolifaciens* P11-C8 (mean 17% conversion), and *Clostridium citroniae* P2-B6 (mean 4%) (**Fig. 1b**). Consistent with our findings, an early study showed that isolates of *Eubacterium lentum* (later reclassified as *Eggerthella lenta*) could produce small amounts of 3-oxoLCA in anaerobic resting cell culture^20^. We verified that the type strains of a subset of these organisms produced a comparable amount of 3-oxoLCA in vitro (**Extended Data Fig. 1e**), giving us access to the associated genome sequences for subsequent biochemical characterization. Taken together, these data indicate that a variety of human gut bacteria from an array of families within the Actinobacteria and Firmicutes phyla produce 3-oxoLCA.

Next, we used in vitro T cell culture assay to investigate whether 3-oxoLCA produced by gut bacteria inhibits T_H_17 cell differentiation. Naive CD4+ T cells were isolated from wild-type C57BL/6J (B6Jax) mice and cultured with *E. lenta* culture supernatant under T_H_17-cell differentiation conditions. Consistent with our UPLC-MS results that the type strain of *E. lenta* converts LCA into 3-oxoLCA, supernatant from cultures of this bacterium incubated with LCA significantly inhibited the differentiation of T_H_17 cells. T_reg_ cell differentiation, used as control, was not affected by the addition of *E. lenta* culture supernatants (**Fig. 1c, d** and **Extended Data Fig. 2a, b**). These data suggest that human gut bacteria producing 3-oxoLCA are capable of suppressing T_H_17 cell differentiation in vitro.

### The abundant gut bacterial metabolite isoLCA inhibits T_H_17 cell differentiation

While analyzing UPLC-MS data from the screen, we noticed that *G. pamelaeae, E. lenta*, and *C. citroniae* produced a new peak (retention time 5.7 min, **Fig. 1b**) in addition to 3-oxoLCA. A *Ruminococcus gnavus* isolate (P4-G2) also generated this peak (**Fig. 1b**). Because this compound had an identical m/z (mass-to-charge ratio) to LCA and it had been previously reported that the type strains of *E. lenta* and *R. gnavus* convert DCA into isoDCA, the 3β-OH isomer of DCA^21^, we reasoned that this unknown metabolite was the LCA isomer isoLCA. A pure standard of isoLCA exhibited the same mass and retention time as the unknown peak, suggesting that the unknown compound was isoLCA. Spike-in of pure isoLCA into bacterial cultures that had produced this new compound confirmed this result (**Fig. 1e**). After LCA (mean ∼160 μM) and DCA (mean ∼200 μM), isoLCA is the most abundant bile acid in healthy human cecal contents (mean ∼50 μM)^5^. Although isoLCA is largely enterohepatically reabsorbed and not excreted^5^, micromolar concentrations are still found in human feces (mean 54 μM, range 0-213 μM, **Extended Data Fig. 1a**). IsoLCA is undetectable in the cecal contents of GF B6 mice^1^. Together, these data indicate that specific members of the microbiome produce the abundant gut metabolite isoLCA.

We next investigated whether isoLCA affects also T helper cell differentiation. Naive CD4^+^ T cells isolated from wild-type B6Jax mice were cultured under T_H_17-cell polarizing conditions with different bile acids, including isoLCA. Intriguingly, isoLCA inhibited the differentiation of T_H_17 as efficiently as 3-oxoLCA, while another abundant iso bile acid, isoDCA, failed to do so (**Fig. 2a, b**). While isoLCA caused a dose-dependent reduction in T_H_17 cell differentiation with little effect on cell viability and their total cell number (**Fig. 2c** and **Extended Data Fig. 3a, b**), it had no effect on T_H_1 and T_reg_ cell differentiation (**Extended Data Fig. 3c-f**). These data suggest that isoLCA, like 3-oxoLCA, may also function as a specific inhibitor of T_H_17 cell differentiation. We next tested whether isoLCA affects T_H_17 cell differentiation in vivo. The mouse commensal species segmented filamentous bacteria (SFB) induces robust T_H_17 cell differentiation in the ileum^22^. We gavaged B6Jax mice, which lack SFB and contain few intestinal T_H_17 cells, with faecal material containing high levels of SFB and maintained these mice on either a chow control diet or chow containing 0.3% (w/w) isoLCA (**Fig. 2d**). Consistent with our in vitro observations, isoLCA treatment resulted in significant reduction in T_H_17 cell differentiation induced by SFB colonization without affecting the T_reg_ cell population (**Fig. 2e-g and Extended Data Fig. 3g**). At steady state, compared to control mice, isoLCA treatment reduced the levels of pre-existing T_H_17 cells in the ileal lamina propria of C57BL/6N (B6Tac) mice, which had been colonized by SFB upon weaning (**Extended Data Fig. 3h-j**). IsoLCA treatment also significantly lowered the T_H_17 cell population frequency without affecting the T_reg_ population in the ileal lamina propria of mice treated with anti-CD3 (**Extended Data Fig. 3k-o**). Intraperitoneal injection of anti-CD3 induces acute, but self-limiting, inflammation in the small intestines, leading to a robust increase in the percentages of T_H_17 cells^23^. These data altogether suggest that isoLCA treatment suppresses T_H_17 cell differentiation in mice at a steady state and under inflammatory conditions.

**Fig. 2.**
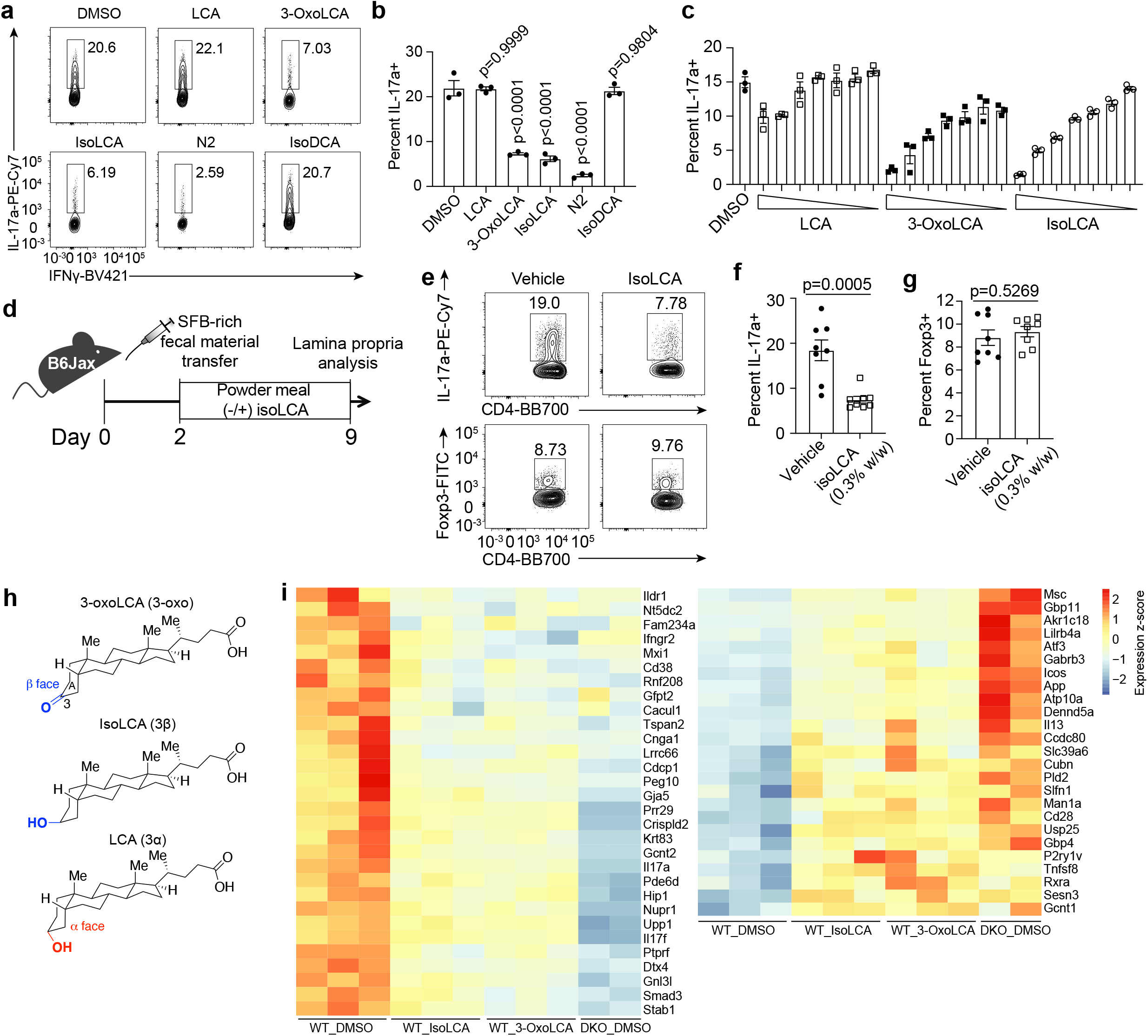
The abundant gut bacterial metabolite isoLCA inhibits T_H_17 cell differentiation. **a**-**c**, IsoLCA inhibited the differentiation of T_H_17 cells in vitro. Representative FACS plots (**a**) and population frequencies of T_H_17 cells cultured in the presence of various bile acids (**b, c**) are shown. Mouse naive CD4+ T cells from wild-type B6Jax mice were cultured under T_H_17 cell polarizing conditions for 3 days and compounds were added 18 hours after TCR activation either at 20 μM (**a, b**) (n = 3 biologically independent samples per condition, data are mean ± SEM, followed by one-way ANOVA with Dunnett’s multiple comparison test, vehicle set as control) or at various concentrations (40, 20, 10, 5, 2.5, 1.25 and 0.625 μM) (**c**) (n = 3 biologically independent samples per condition, data are mean ± SEM). **d-g**, IsoLCA inhibited the differentiation of T_H_17 cells in vivo. Experimental scheme (**d**), representative FACS plots (**e**) and population frequencies of T_H_17 cells (**f**) and T_reg_ cells (**g**) in the ileal lamina propria of SFB-colonized mice are shown. B6 Jax mice were gavaged with SFB-rich faecal pellets and maintained on control or isoLCA-containing powder chow (0.3% w/w) for a week (n=8 mice per group, pooled from two experiments, data are mean ± SEM, followed by unpaired t-test with two-tailed *p*-value). **h**, Three-dimensional structures of 3-oxoLCA, isoLCA, and LCA showing that the C3 oxygenation in 3-oxoLCA and isoLCA is oriented toward the β face while and C3 oxygenation in LCA is oriented toward the α face of the steroidal A ring. **i**, Transcriptional profiling of wild-type (WT) T cells and RORα and RORγ deficient (DKO) T cells, cultured under T_H_17 cell polarization conditions. DMSO or bile acids were added to cells 18 hours after TCR activation. Cells were then harvested and RNA-sequencing was performed. Genes were considered differentially expressed when FDR-adjusted p value <0.05. Heat map represents 55 genes that are commonly regulated by 3-oxoLCA, isoLCA and RORα/RORγ (WT cells, n = 3 mice per condition, DKO cells, n=2 mice per condition).

IsoLCA is structurally similar to 3-oxoLCA. Both compounds possess C3 oxygenation oriented toward the β face of the steroidal A ring. In contrast, LCA, which does not inhibit T_H_17 cell differentiation^1^, possesses an α face-oriented C3 hydroxyl group (**Fig. 2h**). Based on the structural similarities between isoLCA and 3-oxoLCA, we hypothesized that RORγt may also function as a cellular target for isoLCA. To investigate this hypothesis, we used a luciferase reporter plasmid containing the GAL4-DNA-binding domain fused to the RORγt ligand-binding domain upstream of the firefly luciferase gene and assessed transcriptional activity with a luciferase assay. Like 3-oxoLCA and a RORγt synthetic inhibitor ML209^1^, isoLCA treatment reduced RORγt reporter activity in HEK 293 cells, suggesting that isoLCA inhibits transcriptional activity of RORγt (**Extended Data Fig. 3p**). We next performed RNA sequencing analyses with CD4 T cells isolated from wild-type (WT) mice or mice deficient for RORγt and RORα (DKO). Compared to DMSO-treated WT cells, isoLCA- or 3-oxoLCA-treated WT cells exhibited differential mRNA expression of a common set of 125 genes. Among them, 55 genes, including the T_H_17 cell cytokines IL-17a and IL-17f, were also similarly regulated in DMSO-treated DKO cells (**Fig. 2i and Extended Data Fig. 3q**). Together, these data indicate that isoLCA, like 3-oxoLCA, is capable of modulating T_H_17 cell differentiation and function by inhibiting the transcriptional activity of RORγt.

### Bacterial HSDHs convert LCA into 3-oxoLCA and isoLCA

Having determined that isoLCA is a T_H_17 cell modulator, we next sought to uncover gut bacterial producers of this metabolite. We identified *G. pamelaeae* P7-E3, *E. lenta* P7-G7, *R. gnavus* P4-G2, *Clostridium aldenense* P4-B2, and *Clostridium citroniae* P2-B6 as isoLCA producers from our initial screen using LCA as the substrate (**Table S2**). We had previously shown that gut bacteria convert DCA into 3-oxoDCA using a 3α-hydroxysteroid dehydrogenase (3α-HSDH) and 3-oxoDCA into isoDCA using a 3β-hydroxysteroid dehydrogenase (3β--HSDH)^21^. We reasoned that an analogous biosynthetic pathway was responsible for the conversion of LCA into 3-oxoLCA and then isoLCA (**Fig. 3a**). To investigate this hypothesis, we performed an additional round of screening in which we incubated the 990 isolates used in the first screen with 3-oxoLCA (100 μM) as the substrate and monitored production of isoLCA by UPLC-MS.

**Fig. 3.**
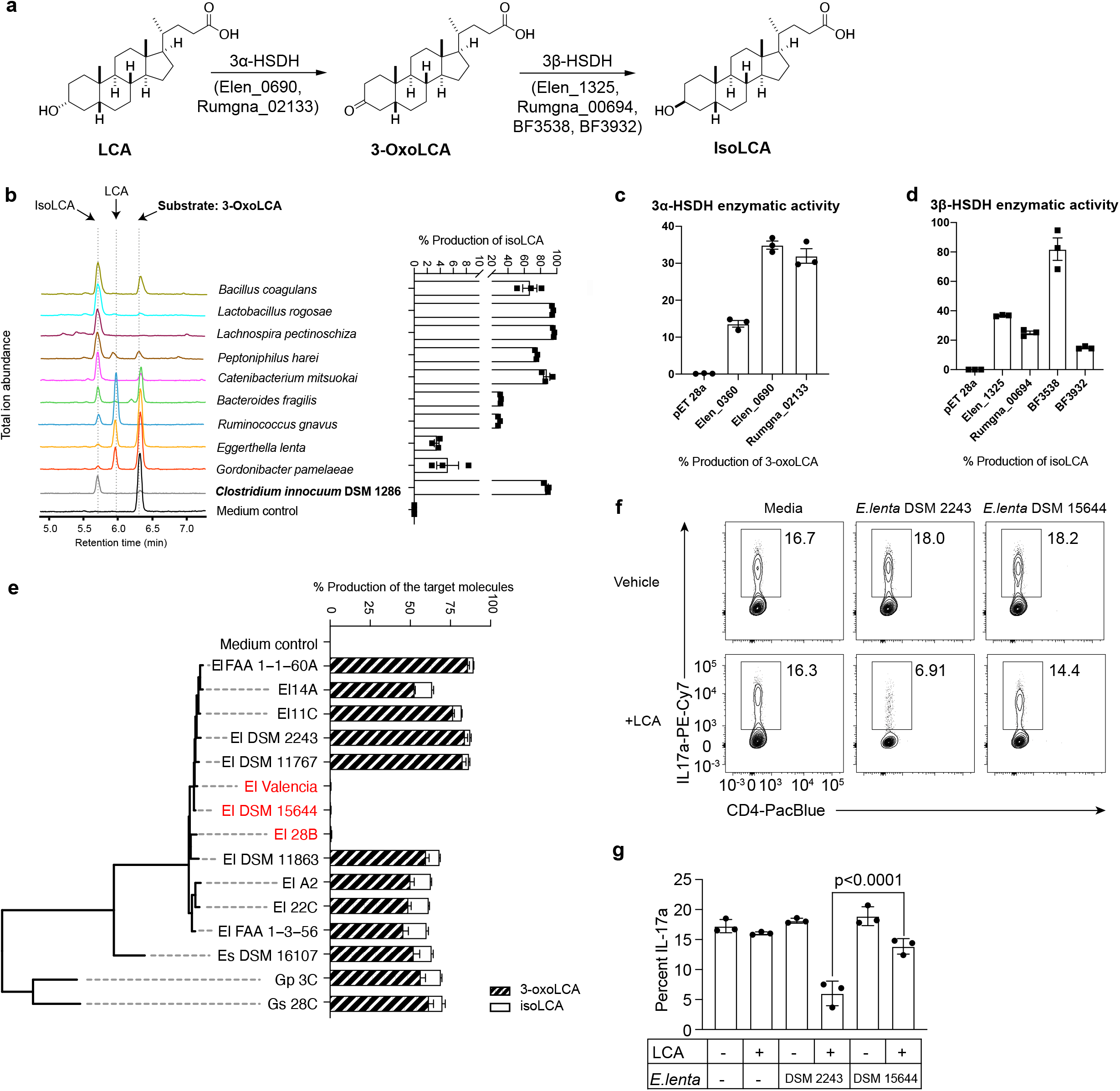
Bacterial hydroxysteroid dehydrogenases (HSDHs) convert LCA to 3-oxoLCA and isoLCA. **a**, Proposed biosynthetic pathway for the conversion of LCA to 3-oxoLCA and isoLCA by *Eggerthella lenta* and *Ruminococcus gnavus* and for the conversion of 3-oxoLCA to isoLCA by *Bacteroides fragilis*. **b**, Representative UPLC-MS traces (left) and percent production of isoLCA (right) by human bacterial isolates incubated with 3-oxoLCA (100 μM) for 48 hours (see **Extended Data Fig. 1b** for overall workflow). *Clostridium innocuum*, which was identified to encode 3β-HSDH homologs in HMP2 metagenomes (see Methods; **Table S9**), was selected for in vitro testing. Production of isoLCA by the *C. innocuum* type strain DSM 1286 verified this prediction (n = 3 biological replicates per group). See **Table S2** for full results. **c, d**, Heterologous expression of candidate HSDHs from *E. lenta* DSM 2243, *B. fragilis* NCTC 9343, and *R. gnavus* ATCC 29149 in *E. coli* revealed that Rumgna_02133, Elen_0360 and Elen_0690 possess 3α-HSDH activity (**c**) while BF3538, BF3932, Rumgna_00694 and Elen_1325 possess 3β-HSDH activity (**d**). *E. coli* lysate was incubated with either 100 μM LCA or 100 μM 3-oxoLCA as a substrate. *E. coli* containing an empty vector was used as a control. All data are reported as percent conversion to product (either 3-oxoLCA or isoLCA) (n = 3 biological replicates per group). See **Extended Data Fig. 4b** for full results. **e**, Cladogram of *E. lenta* and related human isolates and their percent production of 3-oxoLCA and isoLCA (El, *E. lenta*; Es, *Eggerthella sinensis*; Gs, *Gordonibacter sp*., and Gp, *Gordonibacter pamelaeae*). *E. lenta* isolates in red (*E. lenta* 28B, *E. lenta* DSM 15644, *E. lenta* Valencia) that lack a homolog of Elen_0690 did not synthesize 3-oxoLCA from LCA. All strains were incubated with 100 μM LCA as a substrate for 48 hours (n = 3 biological replicates per group). **f, g**, The 3α-HSDH gene of *E. lenta* is required to suppress T_H_17 cell differentiation in vitro. Representative FACS plots (**f**) and population frequencies of T_H_17 cells (**g**) are presented. Naive CD4+ T cells from wild-type B6Jax mice were cultured under T_H_17 cell polarizing conditions for 3 days. Culture supernatants of *E. lenta* DSM 2243 or *E. lenta* DSM15644, an isolate lacking a 3α-HSDH, were added 18 hours after TCR activation (n=3 biologically independent samples per group, data are mean ± SEM, followed by one-way ANOVA with Tukey’s multiple comparison test).

A total of 266 out of the 990 isolates demonstrated more than 1% conversion, and 54 isolates demonstrated more than 50% conversion from 3-oxoLCA to isoLCA. Overall, we identified members of 15 bacterial genera (*Bacillus, Bacteroides, Bifidobacterium, Catenibacterium, Enterocloster, Erysipelatoclostridium, Clostridium, Collinsella, Eggerthella, Gordonibacter, Lachnospira, Lactobacillus, Parabacteroides, Peptoniphilus*, and *Mediterraneibacter*) (**Table S2**) that convert 3-oxoLCA to isoLCA. These data suggest that conversion of 3-oxoLCA to isoLCA requires enzymatic activity found in a diverse array of human gut bacteria. Several strains, including *Lactobacillus rogosae* P2-F2, *Lachnospira pectinoschiza* P2-A2, and *Catenibacterium mitsuokai* P1-A4 exhibited more than 80% conversion of 3-oxoLCA to isoLCA (**Fig. 3b**). We verified that the type strains of a subset of these human isolates could produce a comparable amount of isoLCA in vitro (**Extended Data Fig. 4a**). These data also suggest that collaborative metabolism by several species of gut bacteria could be in part responsible for the production of isoLCA in vivo, with some bacteria converting LCA to 3-oxoLCA and other bacteria converting 3-oxoLCA into isoLCA.

We next sought to identify bacterial enzymes that convert LCA into 3-oxoLCA and isoLCA. We heterologously expressed *E. lenta* and *R. gnavus* genes that had previously been predicted from BLASTP searches for HSDH candidates^21^ in *E. coli*. We then incubated clarified lysate with LCA or 3-oxoLCA to test for 3α- and 3β-HSDH activities, respectively. Conversion of substrate to product was quantified by UPLC-MS (**Extended Data Fig. 4b**). These data indicate that Elen_0690 and Rumgna_02133 convert LCA to 3-oxoLCA (**Fig. 3c**) while Elen_1325 and Rumgna_00694 convert 3-oxoLCA to isoLCA (**Fig. 3d**). Thus, we propose that the former genes are 3α-HSDHs and the latter are 3β-HSDHs.

While the majority of the identified isoLCA-producing bacteria are Gram positive Firmicutes or Actinobacteria, the Gram negative species *Bacteroides fragilis* that we identified in the screen is a robust isoLCA producer. Because *B. fragilis* is a prevalent human gut commensal^24,25^, we reasoned that this 3β-HSDH likely contributes to isoLCA production in humans. To identify this gene, we performed BLASTP searches against the *B. fragilis* NCTC 9343 genome using Elen_1325 and Rumgna_00694 as query sequences (**Table S3**). Seven candidates were selected based on an E value cutoff of 1 E-15. BF3538 was also selected based on predicted secondary structure homologous to known 3β-HSDHs^26^. Heterologous expression of these eight candidate genes followed by incubation of cell lysate with 3-oxoLCA and quantification of isoLCA by UPLC-MS allowed us to identify BF3538 and BF3932 as 3β-HSDHs that produce isoLCA (82% and 15% conversion, respectively, **Fig. 3d** and **Extended Data Fig. 4b**).

The majority of the 3-oxoLCA- and isoLCA-producing strains identified are not genetically manipulable, including the top two 3-oxoLCA producers, *G. pamelaeae* P7-E3 and *E. lenta* P7-G7. These species are closely related members of the family *Eggerthellaceae* (50.8% and 43.9% of protein coding genes, respectively, shared between the type strains). To determine whether the two 3α-HSDHs we identified (Elen_0690 and Elen_0360) are functional in growing bacteria, and to identify non-producer strains for further in vitro and in vivo assays, we utilized a collection of thirteen *Eggerthella* and two *Gordonibacter* human isolates whose genomes are fully or partially sequenced. Through comparative genomics, we identified three *E. lenta* strains (*E. lenta* Valencia, *E. lenta* 28B, *E. lenta* DSM 15644) that lack a homolog for Elen_0690 and two *Gordonibacter* strains (*G. pamelaeae* 3C and *G. species* 28C) that lack a homolog for Elen_0360 (**Table S4**). Neither 3-oxoLCA or its downstream metabolite isoLCA were detected when *E. lenta* Valencia (Elen_0690^-^), *E. lenta* DSM 15644 (Elen_0690^-^), or *E. lenta* 28B (Elen_0690^-^) were cultured with LCA, while *G. pamelaeae* (Elen_0360^-^) and *G*. species 28C (Elen_0360^-^) produced similar amounts of 3-oxoLCA and isoLCA from LCA as control strains containing homologs of both genes (**Fig. 3e**). These data support the hypothesis that Elen_0690 and its homologs, but not Elen_0360 and its homologs, encode 3α-HSDHs responsible for the conversion of LCA to 3-oxoLCA.

To examine whether 3-oxoLCA production via 3α-HSDH modulates T_H_17 differentiation, we again performed an in vitro T cell assay. Isolates of *E. lenta* DSM 2243 (3α-HSDH^+^) or *E. lenta* DSM 15644 (3α-HSDH^-^) were incubated with LCA for 8 hours, and culture supernatants were then added to mouse naive CD4^+^ T cells activated and expanded under T_H_17 polarizing conditions. As expected, T_H_17 differentiation was significantly reduced in cells treated with *E. lenta* DSM 2243 + LCA supernatant compared to those treated with supernatant from either *E. lenta* DSM 15644 + LCA or *E. lenta* DSM 2243 alone (**Fig. 3f, g**). These data suggest that the presence of 3α-HSDH in *E. lenta* affects this organism’s ability to modulate T_H_17 cell differentiation.

To test whether collaborative metabolism by gut bacteria could increase production of isoLCA, we co-cultured a robust 3-oxoLCA producer with organisms possessing 3β-HSDH activity. Co-incubation of *E. lenta* DSM 2243 (3α-HSDH^+^) with either *B. fragilis* NCTC 9343, *R. gnavus* ATCC 29149, or *Clostridium innocuum* DSM 1286 resulted in a higher conversion of LCA to isoLCA compared to *E. lenta* DSM 2243 (3α-HSDH^+^) culture alone (**Extended Data Fig. 5a, Fig. 3e**). In contrast, co-culture of *E. lenta* DSM 15644 (3α-HSDH^-^) with 3β-HSDH-encoding bacteria did not result in conversion of LCA into 3-oxoLCA or isoLCA (**Extended Data Fig. 5a**). These data support a model of synergy for isoLCA production between strains with 3α- and 3β-HSDH activity.

### Gut bacteria produce T_H_17-modulatory LCA metabolites in vivo

We next assessed whether gut bacteria can metabolize LCA in vivo. Age- and sex-matched GF C57BL/6NCrl (B6 GF) mice were colonized with *E. lenta* DSM 2243 (3α-HSDH^+^) or *E. lenta* DSM 15644 (3α-HSDH^-^). Because LCA is absent from GF animals, colonized mice were then fed chow alone or chow supplemented with LCA (0.3% w/w) (**Fig. 4a**)^27^. Significantly higher levels of 3-oxoLCA were detected in the cecal contents of 3α-HSDH^*+*^-colonized mice compared to those of 3α-HSDH^-^-colonized mice (mean 34 picomol/mg wet mass vs mean 6 picomol/mg wet mass, p<0.0001) (**Fig. 4b**).

**Fig. 4.**
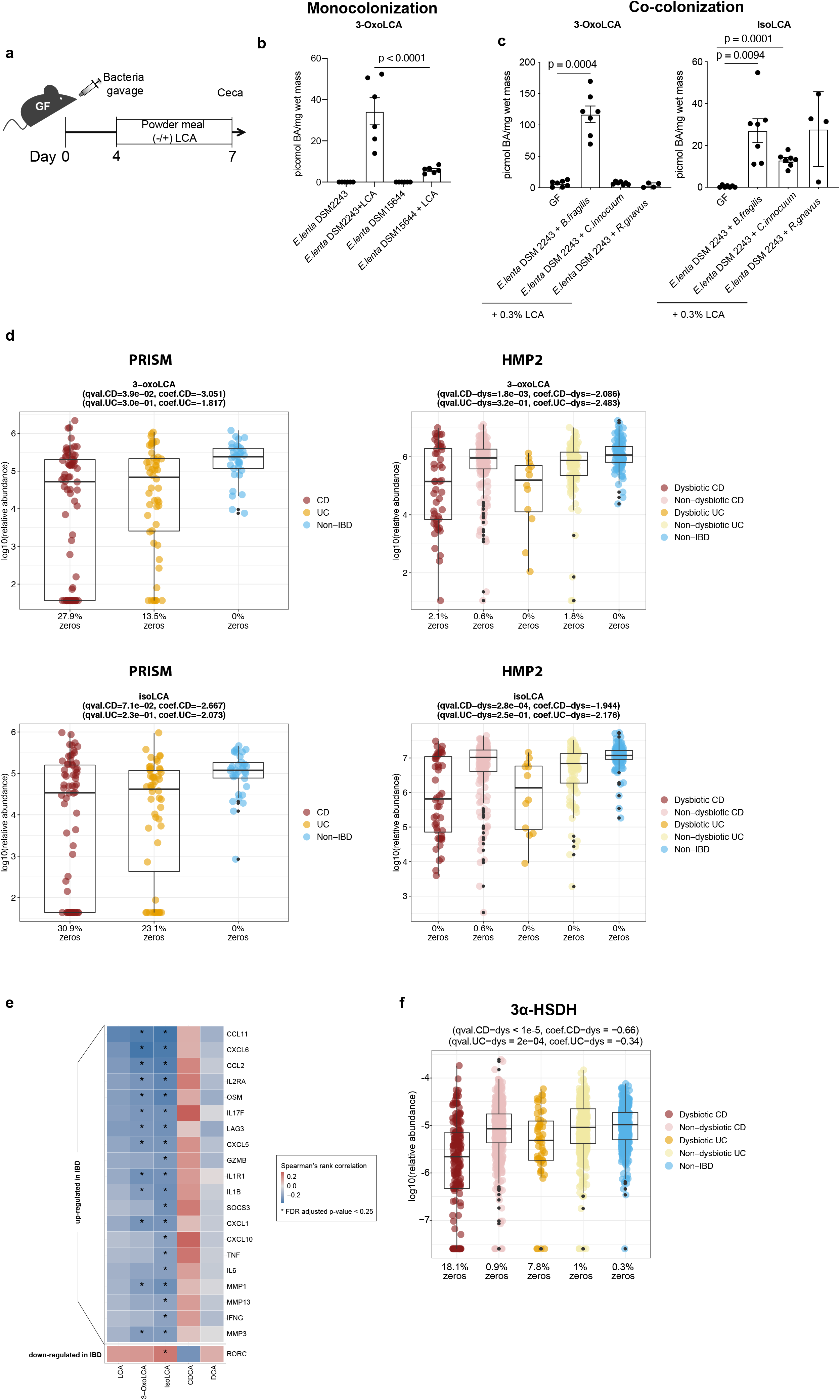
Gut bacteria contribute to differential abundance of immune modulatory LCA metabolites in vivo. **a-c**, Cecal 3-oxoLCA and isoLCA levels in gnotobiotic mice colonized with gut bacteria. Mice were fed control chow or chow containing 0.3% LCA (w/w) and their cecal contents were analyzed by UPLC-MS for LCA metabolites. **a**, Experimental scheme for gnotobiotic experiments. **b**, GF B6 mice were monocolonized with either the *E. lenta* type strain (DSM 2243) or a strain lacking a 3α-HSDH (DSM15644) (n=5 mice in *E. lenta* DSM 2243, n=5 in *E. lenta* DSM 15644 - only, n=6 mice in *E. lenta* DSM 2243+LCA, n=6 in *E. lenta* DSM 15644+LCA groups, pooled from two experiments, data are mean ± SEM, followed by one-way ANOVA with Tukey’s multiple comparisons). **c**, GF B6 mice were co-colonized with *E. lenta* DSM 2243 and individual bacterial strains harboring 3β-HSDH activity (*B. fragilis* NCTC 9343, *C. innocuum* DSM 1286, or *R. gnavus* ATCC 29149) (n=7 mice per group except n=4 for *E. lenta* + *R. gnavus* group, pooled from two experiments, data are mean ± SEM, followed by one-way ANOVA with Dunnett’s multiple comparisons). **d**, Relative abundance distributions of 3-oxoLCA and isoLCA as derived from metabolomic profiles of the PRISM and HMP2 IBD cohorts. Both metabolites were significantly depleted in PRISM CD subjects (n = 68) relative to PRISM controls (n = 34) and in HMP2 dysbiotic CD samples (n = 48) relative to non-dysbiotic controls (n = 169; 50 subjects). The percentage of zeros in each condition are added as x-axis tick labels. Boxplot ‘boxes’ indicate the first, second (median), and third quartiles of the data. The points outside of boxplot whiskers are outliers. Statistical analysis was performed using a linear mixed-model and its coefficient and significance, FDR-adjusted *p*-values, are shown. See **Table S6** for full results. **e**, T_H_17/IL-17-related genes upregulated in IBD were significantly negatively correlated with 3-oxoLCA and isoLCA (FDR-adjusted *p*-value < 0.25) but not the other 3 control bile acids (LCA, DCA, and CDCA). Differentially expressed T_H_17/IL-17-related genes with at least one significant association are shown. This analysis was based on a subset of n = 71 subject-unique samples with matched metagenomic, metabolomic, and host transcriptomic profiling in the HMP2 cohort (33 CD, 21 UC, and 17 non-IBD controls). Correlations were based on residual transcript and metabolite abundance after correcting for diagnosis, consent age, and antibiotic use. See **Table S8** for full results. **f**, Relative abundance distributions of differentially abundant 3α-HSDH homologs profiled from HMP2 metagenomes (*n* = 1,595 samples from 130 subjects; linear mixed-effects model coefficient for dysbiosis within diagnosis, FDR-adjusted *p*-values < 0.05). Boxplots show median and lower/upper quartiles with outliers outside of boxplot ‘whiskers’ (indicating the inner fences of the data). The percentage of zeros in each condition are added as x-axis tick labels. See **Table S9** for full results.

We then asked whether collaborative metabolism by gut bacteria can enhance the production of the T_H_17 cell-inhibitory metabolites of LCA in vivo. B6 GF mice were co-colonized with *E. lenta* DSM 2243 and one of three bacterial species with robust 3β-HSDH activity (*B. fragilis* NCTC 9343, *R. gnavus* ATCC 29149, or *Clostridium innocuum* DSM 1286), followed by feeding chow supplemented with LCA (0.3% w/w). UPLC-MS analysis revealed that mice colonized with each pair of bacteria harbored a higher total combined concentration of 3-oxoLCA and isoLCA in cecal contents than mice colonized with *E. lenta* DSM 2243 alone (**Fig. 4b, c**). The *E. lenta* and *B. fragilis* strain pair produced the highest combined concentration of 3-oxoLCA and isoLCA (mean 144 picomol/mg wet mass). Co-colonization of GF mice with *E. lenta* DSM 15644 (3α-HSDH^-^) and *B. fragilis* resulted in low levels of 3-oxoLCA (mean 7 picomol / mg wet mass) and undetectable levels of isoLCA in the cecal contents, indicating that in vivo production of 3-oxoLCA and isoLCA by gut bacteria requires 3α-HSDH activity (**Extended Data Fig. 5b**).

### Differential abundance of immunomodulatory LCA metabolites in IBD

The finding that 3-oxoLCA and isoLCA exhibit T_H_17-modulatory properties led us to consider the roles of these metabolites in the context of gut inflammation. Specifically, we examined metabolomic profiles of patients with Crohn’s disease (CD) and ulcerative colitis (UC) in the cross-sectional Prospective Registry in IBD Study at MGH (PRISM) IBD cohort^28^ and the longitudinal Integrative Human Microbiome Project (HMP2/ iHMP) IBD cohort^29^. Consistent with the function of 3-oxoLCA and isoLCA in our mouse data, levels of 3-oxoLCA and isoLCA were significantly decreased in CD patients (n=68) from the PRISM cohort relative to non-IBD controls (n = 34; linear model, FDR-corrected *q*-values =0.0392 and 0.0710, respectively). Furthermore, within the HMP2 cohort, levels of both 3-oxoLCA and isoLCA were significantly decreased in CD patients in a dysbiotic state (n=48) compared to their non-dysbiotic baselines (n = 169; 50 subjects, FDR *q*-value =0.0018 and 0.00028, respectively; **Fig. 4d** and **Table S6**). However, levels of other bile acids that do not suppress T_H_17 cell differentiation were not significantly affected in CD patients (**Extended Data Fig. 6**). These data indicate that the anti-inflammatory metabolites 3-oxoLCA and isoLCA are negatively associated with CD in humans.

Next, we investigated whether 3-oxoLCA/isoLCA abundance was associated with the expression of host T_H_17/IL-17 pathway-related genes that are known to be perturbed in IBD. We curated a set of known T_H_17/IL-17 related genes, including genes previously reported as upregulated in IBD and enriched in the IL-17 signaling pathway^29^ as well as a T_H_17 signature^30^ (**Table S7**). We identified a subset of these T_H_17/IL-17-related genes (n=21) that were differentially expressed (DE) in IBD patients from the HMP2 cohort (linear models, FDR-corrected *q*-value < 0.25; see **Methods** and **Table S8**). We then correlated these DE T_H_17/IL-17 signaling-related genes with LCA, 3-oxoLCA, and isoLCA along with two other control bile acids, CDCA and DCA (a human primary bile acid and an abundant secondary bile acid, respectively). This test used Spearman’s *r* on residual abundance values after linear modeling with diagnosis, antibiotics and age as covariates, thus enriching for association between genes and metabolites that cannot be explained by confounding factors such as diagnosis (see **Methods**). Strikingly, 20 out of the 21 genes with FDR-significant correlations (FDR *q*-values < 0.25) were upregulated in IBD and also displayed a significant negative correlation with 3-oxoLCA and/or isoLCA, but not with the three other bile acids (**Fig. 4e**). These data imply that 3-oxoLCA/isoLCA may specifically contribute to IBD by biasing the T_H_17/IL-17 signaling axis.

We further explored the associations between 3α- and 3β-HSDH-related microbial features and 3-oxoLCA/isoLCA during gut inflammation. UniRef^31^-based homologs of 3α-/3β-HSDH-related genes from *E. lenta* DSM 2243, *R. gnavus* ATCC 29149, and *B. fragilis* NCTC 9343 were identified from existing gene-level functional profiles of HMP2 metagenomes (see **Methods**). Interestingly, both 3α-HSDH homologs were significantly depleted in CD and UC patients’ dysbiotic samples relative to non-dysbiotic controls (linear mixed-effects models, FDR *q*-value <1e-5 in dysbiotic CD and 2e−04 in dysbiotic UC; **Methods**; **Table S9**; **Fig. 4f**), and 3β-HSDH homologs were significantly depleted in UC patients’ dysbiotic samples compared to non-dysbiotic controls (**Methods**; **Table S9**; **Extended Data Fig. 7**). Moreover, by correlating the differentially abundant 3α-/3β-HSDH homologs with the five metabolites of interest (LCA, 3-oxoLCA, isoLCA, CDCA and DCA), we found that most 3α-/3β-HSDH homologs were positively correlated with 3-oxoLCA/isoLCA (Spearman’s *r* on residual abundance, FDR *q*-values < 0.25; **Fig. S9**). Additionally, 3-oxoLCA/isoLCA were significantly associated with the species that exhibited 3α-/3β-HSDH activity and differential abundance between dysbiosis states within each IBD phenotype (CD and UC; **Table S10**; **Methods**). Collectively, these results suggest that the decreased abundances of 3-oxoLCA and isoLCA in dysbiotic IBD are linked to changes in the abundance of the 3α-/3β-HSDHs and the species that encode these enzymes.

## Discussion

Here, we identified gut bacteria and enzymes that produce 3-oxoLCA and the abundant gut metabolite isoLCA, bile acids that inhibit T_H_17 cell differentiation and function. The findings that levels of 3-oxoLCA and isoLCA as well as the bacterial genes responsible for the biosynthesis of these metabolites are reduced in IBD patients suggest that bacterial production of these molecules may help to maintain homeostatic immune balance in the gut. Negative correlations between 3-oxoLCA and isoLCA and host genes in the T_H_17/IL-17 signaling axis further suggest that these metabolites modulate the immune response at least in part by regulating T_H_17 function in humans. Consistent with the conclusion these metabolites promote human health, increased levels of 3-oxoLCA- and isoLCA-producing bacteria and their biosynthetic genes were found in centenarian individuals compared to elderly and young control subjects (Kenya Honda, personal communication). Prior work has also demonstrated that bacteria such as *R. gnavus* and *B. fragilis* produce certain secondary bile acids, including isoDCA, and enhance the differentiation of peripheral T_reg_ cells^16,17^. Our findings add to a growing list of gut microbe-metabolite pairs that control host immune responses by directly modulating a distinct subset of immune cells. Given the growing recognition of the importance of bile acid molecules in regulating host physiology and immune responses, gaining a deeper understanding of the role of host-microbiota networks in mediating bile acid biotransformations will offer us opportunities to devise therapeutic interventions for diseases such as IBD, metabolic diseases, and cancers of the enterohepatic system.

## Methods

### Mice

C57BL/6J mice were purchased from Jackson Laboratory. SFB-containing C57BL/6N mice were purchased from Taconic Biosciences. GF C57BL/6NCrl mice were purchased from Charles River Laboratories and maintained in GF isolators at Harvard Medical School. For the bile acid feeding experiments, irradiated powder meal (Teklad Global 19% protein extruded diet, #2019) was evenly mixed with a measured amount of bile acid, provided in glass feeder jars, and replenished when necessary. Colonization of mice with SFB was performed using fresh faecal samples from *il23r-/- Rag2-/-* double-knockout mice that are known to carry higher levels of SFB compared to conventional C57BL/6N mice. Faecal samples were homogenized in sterile 1X DPBS using a 100-μm cell strainer and a 5mL syringe plunger. Approximately a quarter of the mouse faecal pellet in 200 μL 1X DPBS was introduced into each mouse using a 20G gavage needle. Successful colonization was assessed by quantitative PCR using the following primers: SFB-F, 5’-GACGCTGAGGCATGAGAGCAT-3’; SFB-R, 5’-GACGGCACGAATTGTTATTCA-3’. For anti-CD3 experiments, B6Tac mice were fed a control powder chow or a powdered chow containing isoLCA (0.3% w/w) 4 days prior to anti-CD3 injection and given an anti-CD3 injection (10 μg/mouse). 3 days later, mice were euthanized, and ∼10 cm of distal small intestines were harvested for lamina propria immune cell analysis. For gnotobiotic experiments, age- and sex-matched GF mice were orally gavaged with bacterial cultures and maintained in an Isocage system (Tecniplast). Control powder meal (Teklad Global 19% protein extruded diet, #2019) or a chow evenly mixed with 0.3% 3-oxoLCA (w/w) were autoclaved and provided to mice during the experiment. Successful colonization was assessed by quantitative PCR (see **Table S1** for primer sequences and **Table S5** for qPCR data). All animal procedures were approved by the Institutional Animal Care and Use Committee at Harvard Medical School.

### Chemical synthesis of 3-oxoLCA and isoLCA

3-oxoLCA was prepared in large scale by the oxidation of LCA according to a previously reported protocol^**1**^. Detailed synthesis methods and characterization data of isoLCA are included in Supplementary Information (**Fig. S1-6**).

### Isolation of lamina propria lymphocytes

Harvested intestines were cut open and rinsed in ice-cold phosphate-buffered saline (PBS). Associated fats were carefully removed and incubated in pre-warmed 1XHBSS (without calcium and magnesium) supplemented with 1 mM dithiothreitol, 2 mM EDTA and 0.5% fetal bovine serum (FBS) at 37 °C for 20 min in a shaking incubator. Then, the tissues were briefly rinsed in warm RPMI and dissociated in digestion media (RPMI supplemented with 50 μg/mL Liberase TM, 50 μg/mL DNase I and 1% FBS) at 37 °C for 40 min in a shaking incubator. Mononuclear cells were collected at the interface of a 40%/80% Percoll gradient (GE Healthcare). Cells were then analyzed by flow cytometry. The distal 10 cm of the small intestine was considered ileum.

### In vitro T cell culture

Naïve CD4+ (CD25-CD4+ CD25-CD62L+ CD44-) T cells were isolated from the spleens and the lymph nodes of mice by FACS sorting. 96-well T-bottom plates were pre-coated with 50 μL of hamster IgG (MP Biomedicals) at 37 °C for 1 hour. Following multiple washes with 1XDPBS, 40,000 naive CD4+ T cells were seeded in T cell media (RPMI supplemented with 10% fetal bovine serum, 25 mM glutamine, 55 µM 2-mercaptoethanol, 100 U/mL penicillin, 100 mg/mL streptomycin) and their T cell receptor downstream signaling pathways (TCR activation) were activated with soluble anti-CD3 (clone 145-2C11, 0.25 µg/mL) and anti-CD28 (clone 37.51, 1 µg/mL). For T_H_1 cell differentiation, 100 U/mL of IL-2 (Peprotech) and 10 ng/mL of IL-12 (Peprotech) were added. For T_H_17 cell differentiation, IL-6 (eBioscience, 20 ng/mL) and human TGF-β1 (Peprotech, 0.3 ng/mL) were added. For T_reg_ culture, 100 U/mL of IL-2 (Peprotech) and human TGF-β1 (Peprotech, 5 ng/mL) were added. Bacterial culture supernatants or small molecules including bile acids were added 18 hours after TCR activation. Cells were harvested and assayed by flow cytometry on day 3.

### Flow cytometry

Cells harvested from in vitro culture or in vivo mice experiments were stimulated with 50 ng/mL PMA (Phorbol 12-myristate 13-acetate, Sigma) and 1 µM ionomycin (Sigma) in the presence of GolgiPlug (BD) for 2 hours to determine cytokine expression. After stimulation, cells were stained with various surface marker antibodies supplemented with LIVE/DEAD Fixable dye for dead cell exclusion. After washing, cells were then fixed, permeabilized with a FoxP3/Transcription factor staining kit (eBioscience) and intracellularly stained for cytokines and/or transcription factors. Flow cytometry data were acquired on an LSR II flow cytometer or Symphony flow cytometer (both BD) and data were analyzed with FlowJo software (TreeStar) following the gating strategy in **Fig. S6**.

### Luciferase reporter assay

The transcriptional activity of the fusion protein of RORγt ligand-binding domain and GAL4-DNA-binding domain is reported by luciferase expression and reporter assays were conducted as previously described^1^. Briefly, 50,000 human embryonic kidney (HEK) 293 cells per well were plated in 96-well plates in antibiotic-free Dulbecco’s Modified Eagle Media (DMEM) containing 1% fetal calf serum (FCS). 16 hours later, cells were transfected with a DNA mixture containing 50 ng of firefly luciferase reporter plasmid (Promega pGL4.31 [luc2P/Gal4UAS/Hygro]), 5 ng of Renilla luciferase plasmid (Promega pRL-CMV), and 50 ng of Gal4-DNA binding domain-human RORγt-ligand-binding domain fusion protein plasmid per each well. Transfections were performed using GeneJuice (Millipore) according to the manufacturer’s instruction. Bile acids or vehicle control were added 8 hours after transfection and luciferase activity was measured 18 hours later using the luciferase assay kits (Biotium).

### RNA-seq analysis

Total RNA was isolated using Qiagen RNeasy Plus Mini Kit according to the manufacturer’s protocol and quantified using Agilent TapeStation RNA assay on Agilent 4200 TapeStation instrument. Libraries were prepared using KAPA mRNA HyperPre kit following the manufacturer’s instruction. Briefly, 50 ng of total RNA per sample was used to capture total mRNA and cDNA synthesis, adapter ligation, and amplification were conducted subsequently. Following clean-up, the resulting purified libraries were analyzed by Agilent High Sensitivity D1000 ScreenTape assay on Agilent 4200 TapeStation instrument. Then, the libraries were pooled equimolarly and run on an Illumina NextSeq 500 instrument with three runs: a Mid-Output 150-cycle kit and two High-Output 150-cycle kits (to obtain sufficient counts of paired-end 75bp reads). The pool was loaded for these runs at 2.1pM, with 5% PhiX spiked in as a sequencing control. The basecall files were demultiplexed through the Harvard BPF Genomics Core’s pipeline, and the resulting FASTQ files were used in subsequent analysis. Raw sequencing reads were aligned to the Ensembl reference genome GRCm38 and gene counts were quantified using Salmon (v. 1.2.1)^33^. DESeq2 (v.1.28.1) was used for differential expression analysis^34^ using the Wald test with Benjamini-Hochberg correction to determine adjusted p-value < 0.05. Gene Ontology analysis was performed by clusterProfiler (v.3.16.0)^35^. Heatmaps were produced using phetmap (v.1.0.12).

### Bacterial culturing

Culturing of human gut bacteria was performed in an anaerobic chamber (Coy Laboratory Products) with a gas mixture of 5% hydrogen and 20% carbon dioxide (balance nitrogen) unless otherwise stated.

### Human stool microbial isolation and cultivation

Human faecal samples were obtained from patients after faecal microbiota transplant (FMT) under an Institutional-Review-Board-approved protocol at Weill Cornell Medicine. Human isolate screening was performed using a published protocol^36^ with the following modifications. Two frozen faecal samples with the highest levels of 3-oxoLCA in the cohort (p#3 [faecal 3-oxoLCA 44 pmol/mg], and p#27 [faecal 3-oxoLCA at 83 picmol/mg], roughly 0.1 g/sample). The faecal were divided in half. One half was homogenized in reduced phosphate-buffered saline (PBS) (Genesee Scientific) and serially diluted and plated directly onto Cullen-Haiser Gut (CHG) agar^32^, which consists of brain heart infusion media (Bacto BHI, BD) supplemented with 1% BBL vitamin K1-hemin solution (BD), 1% trace minerals solution (ATCC), 1% trace vitamins solution (ATCC), 5% FBS (Genesee), 1 g/L cellobiose (Sigma), 1 g/L maltose (Sigma) and 1 g/L fructose (Sigma), and further supplemented with 0.5% (w/v) arginine (Sigma), and cultured at 37 °C. The other half was treated with an equal volume of 70% (v/v) ethanol (Sigma) for 4 hours at room temperature under ambient aerobic conditions to kill vegetative cells, washed three times with PBS, and plated on CHG agar containing 0.1% sodium taurocholate (TCA, Sigma) anaerobically for spore germination. The picked colonies were restreaked to confirm purity and then cultured in 600 μL CHG media containing 0.5% arginine. Pure human isolates (990 in total) were archived and stored as glycerol stocks at −80 °C in eleven 96-well plates. To assess the diversity of the cultured isolates, we performed Genewiz 16S-EZ sequencing. Individual colonies were incubated in 600 μL CHG media with 0.5% arginine for 48 hours, after which 100 μL aliquots from each fresh culture from the same subject were pooled together. DNA extracts were prepared using Allprep Bacterial DNA/RNA/Protein Kit (Qiagen) following the manufacturer’s instructions and further submitted to Genewiz for bacterial 16S-EZ sequencing (V3 and V4 hypervariable regions) using Illumina® MiSeq with 2×250 bp configuration and data analysis. To screen human isolates for LCA metabolism, isolates were retrieved from the stock plates and cultured in 600 μL CHG media containing 0.5% arginine for 48 hours at 37°C in 96-well plates. Each isolate as well as the negative controls were then diluted 1:10 in new media containing 100 μM LCA or 100 μM 3-oxoLCA for an additional 48 hours. 0.2 mL cultures were harvested and extracted. This experiment was conducted once per substrate for all isolates from the original eleven library plates. Following bile acid analysis (see below), we prioritized the positive metabolizers and performed 16S ribosomal RNA gene sequencing (universal 16S-F, 5’-GAGTTTGATCCTGGCTCAG-3′; universal 16S-R, 5′-GGCTACCTTGTTACGACTT-3′) to enable taxonomic characterization for individual isolates. Positive producer function was verified in biological triplicate using single culture conditions (see below).

### Single culturing

Individual strains were plated from glycerol stocks onto CHG agar supplemented with 0.5% (w/v) arginine and grown for 3 days. Colonies were then inoculated into 3 mL of CHG liquid media in Falcon™ Round-Bottom polystyrene tubes and grown for 48 hours at 37 °C to provide starter cultures, which were diluted 1:100 in triplicate into 5 mL fresh CHG media containing either 100 μM of the corresponding substrate (either LCA [Sigma] or 3-oxoLCA [Steraloids]). Cultures were grown for 48 hours at 37 °C. An aliquot of culture (0.5 mL) was harvested and used for bile acid quantification. The experiments were performed in triplicate and repeated twice unless otherwise stated.

### Co-culturing

Starting from single colonies, individual bacterial strains were grown anaerobically for 48 hours in 3 mL of CHG media at 37 °C. These starter cultures were normalized to an OD_600_=0.1 by dilution into fresh media. 10 μL of each normalized starter culture was diluted in 5 mL of CHG media containing 0.75% (w/v) arginine. LCA (100 μM final concentration) or T-LCA (100 μM final concentration, Steraloids) was then added into the media. Cultures were grown for 48 hours at 37 °C and 0.5 mL aliquots were harvested for bile acid analysis.

### Bacterial supernatant assay for in vitro T cell culture

Seed cultures from brain heart infusion media (Bacto) supplemented with 5 mg/L hemin, 2.5 uL/L Vitamin K, 500 mg/L cysteine HCl (BHI+), and 1% arginine were diluted into OD_600_=0.1 in ISP2 + 1% arginine minimal media, which consists of 4 g/L yeast extract (Bacto), 10 g/L malt extract (Sigma), 4 g/L dextrose (Sigma), 10g/L arginine, pH 7.2, containing 800 μM LCA and grew for 8 hours. Supernatants were harvested by centrifugation (12,000 g for 10 minutes) and subsequently passed through 0.2 μm syringe filters. 10 μL of supernatant was added to 200 μL T cell culture.

### *E. coli* heterologous expression and lysis assays

Candidate genes were placed into pET28 expression vectors under an isopropyl β-d-1-thiogalactopyranoside (IPTG)-inducible operon for heterologous expression. Plasmids were transformed into BL21 expression cell lines containing either the pLysS (Elen and Rumgna) or Rosetta (BF) enhancement cassettes. A negative control of the empty pET28a vector was transformed into both cell lines. All cells were cultured at 37 °C until an OD_600_ between 0.6 and 1.0 was reached. Expression was induced by the addition of IPTG (500 μM final concentration). The cultures were incubated overnight at 18 °C and harvested the following day by centrifugation at 4,100 x g before pellets were stored at −80 °C. The pellets were thawed and resuspended in lysis buffer (50 mM Sodium Phosphate Buffer/ 300 mM NaCl/ 10% glycerol, pH = 8). Cells were lysed by sonication and the lysate clarified by centrifugation at 18,200 x g for 45 minutes. The lysate was then incubated with 100 μM substrate (LCA or 3-oxoLCA) for 6 hours at 37 °C. Mixtures were then frozen to quench the reaction and stored at −80 °C until extraction and analysis (described below). Soluble expression was confirmed by SDS-PAGE or immunoblot (**Fig. S7a-d**). Initial gel analysis was performed using SDS-PAGE immediately following lysate clarification using Coomassie blue staining to visualize protein bands. In the case of the *B. fragilis* candidates and select candidates from *E. lenta* and *R. gnavus*, an anti-His tag (Cell Signaling, 2365S) immunoblot was performed with transfer verified by subsequent Amido Black total protein staining.

### Bile acid analyses

Bile acid analyses were performed using a previously reported method^37^. Stock solutions of all bile acids were prepared by dissolving compounds in molecular biology-grade DMSO (Sigma). These solutions were used to establish standard curves. Glycocholic acid (GCA) (Sigma) was used as the internal standard. HPLC-grade solvents were used for preparing and running UPLC-MS samples. All data is analyzed using Agilent ChemStation and expressed as percent conversions to the predicted product(s) (LCA, 3-oxoLCA, isoLCA) or concentration in μM. Note that the four isomers of LCA that have been reported in human feces, LCA, isoLCA, allolithocholic acid (alloLCA), and isoallolithocholic acid (isoalloLCA), were separable by UPLC-MS^1^.

### Sample preparation for native bacterial culture and *E. coli* cell lysis

Bacterial cultures or cell lysates were acidified to pH=1 using HCl (Sigma) and extracted twice with ethyl acetate (Sigma). The organic phase was collected and dried using a speedvac (Thermo Scientific) for 96-well plate cultures or a turbovap (Biotage) for bacterial tube cultures or eppendorf tube lysates, respectively. Dried extracts were solubilized in 75% HPLC-grade methanol (EMD Millipore) in dH_2_O and analyzed by UPLC-MS (Agilent Technologies 1290 Infinity II UPLC system coupled online to an Agilent Technologies 6120 Quadrupole LC/MS spectrometry in negative electrospray mode) using a published method^37,38^ with modifications outlined as follows. Extracted bile acid solutions were injected onto a Phenomenex 1.7 μm, C18 100 Å, 100 × 21 mm LC column with a flow rate of 0.350 mL/min using 0.05% formic acid in water as mobile phase A and acetone as mobile phase B. The following gradient was applied: 0-1 min: 25-60% B, 1-5 min: 60-70% B, 5-6 min: 70-100% B, 6-7 min: 100% B isocratic, 7-8 min: 100-25% B, 8-10 min: 25% isocratic.

### Sample preparation for mouse and human tissue bile acids

Bile acids were extracted from mouse cecal and human faecal samples and quantified by UPLC-MS as previously reported^37^. GCA or β-muricholic acid (βMCA, Steraloids) was used as the internal standard for mouse and human samples, respectively. The limits of detection of individual bile acids in tissues (in pmol/mg wet mass) are as follows: β-muricholic acid, 0.10; GCA, 0.25; T-LCA, 0.04; LCA, 0.12; 3-oxoLCA, 0.18; and isoLCA, 0.29.

### Genomic and meta-omic sequence analysis

#### *B. fragilis* NCTC 9343 3β-HSDHs

BLASTP searches of the *B. fragilis* NCTC 9343 genome were performed using the JGI Integrated Microbial Genomes and Microbiomes database (version 5.0 in March 2020)^39,40^ with 3β-HSDHs Elen_1325 and Rumgna_00694 as query sequences using an E value cutoff of 1E-2. All candidate genes with E values below 1E-15 were selected for heterologous expression assays. Secondary structure prediction analysis using the JPRED4 server^26^ was then performed on the remaining hits. The predicted structures of the known 3β-HSDHs Elen_1325 and Rumgna_00694 were compared with those of the remaining hits. The best match to the known 3β-HSDHs, BF3538 (CAH09226.1), was also selected for heterologous expression.

### Comparative genomic analysis for *E. lenta* isolates

Genetic variation between the *E. lenta* DSM 2243 type strain and other human isolates was determined using comparative genomic analysis pipelines as previously reported in ElenMatchR v1.0.9003^41^. Elen_0360 (ACV54351.1), Elen_0690 (ACV54671.1), and Elen_1325 (ACV55294.1) for *Eggerthellaceae* isolates (**Table S4**). A phylogenetic tree was created using PhyloPhlAn^42^ and visualized with Ggtree^43^.

### Identification of LCA derivatives from PRISM and HMP2 cohorts

The raw LC-MS data were acquired using the same C18-negative mode LC-MS methods described in the HMP2 and PRISM studies^29,44^. Peaks of unknown ID were confirmed using authentic standards run alongside with the quality control reference stool pool generated in the HMP2 study. The LCA derivatives were confirmed by matching their *m/z* in negative mode and retention time, and subsequently verified using LC-MS/MS (**Fig. S8**). Extracted ion chromatograms (EICs) were generated using QualBrowser (Xcalibur 4.1.31.9; Thermo Fisher Scientific, Waltham, MA). The commercial standards used are: LCA (Sigma, L6250), 3-oxoLCA (Steraloids (C1750-000, isoLCA (Steraloids, C1475-000), isoalloLCA (Steraloids, C0700-000), alloLCA (Steraloids, C0680-000), DCA (Sigma, D2510), 3-oxoDCA (Steraloids, C1725-000), and isoDCA (Steraloids, C1165-000). LCA peak in PRISM: FFA_Cluster_0731, m/z = 375.2898 at 12.42 min, and in HMP2: C18n_QI48, m/z = 375.2905 at 11.98 min; 3-oxoLCA in PRISM: FFA_Cluster_0722, m/z = 373.2744 at 12.63 min and in HMP2: C18n_QI6169, m/z = 373.2749 at 12.2-12.35 min; IsoLCA in PRISM: FFA_Cluster_0733, m/z = 375.2901 at 11.73 min and in HMP2: C18n_QI6230, m/z = 375.2906 at 11.31 min (**Fig. S8**).

### Statistical analysis of PRISM and HMP2 IBD multi-omic datasets

#### Data overview

We used two publicly available IBD metabolomics datasets for determining the differential abundance (DA) of bile acids in disease/dysbiotic conditions, specifically 1) the Prospective Registry in IBD Study at MGH (PRISM)^28^ and 2) the IBDMDB study within the integrative Human Microbiome Project (HMP2)^29^. Additional multi-omic profiles from the HMP2 were further used to associate metabolite abundance with microbial species, gene products, and host gene expression.

The PRISM dataset used is a cross-sectional cohort incorporating subjects diagnosed with Crohn’s disease (CD; n = 68); ulcerative colitis (UC, n = 53); and non-IBD controls (n = 34). As with all metabolomics here, PRISM stool samples were subjected to metabolomic profiling using a combination of four LC-MS methods. Paired metagenomic profiles from PRISM samples were not used in this study. Differential abundance in the PRISM cohort was determined as described below based on diagnosis (i.e. comparing the CD and UC subpopulations with controls). Metabolomics profiles from the PRISM cohort were taken from the associated publication’s supporting information^28^.

The IBDMDB HMP2 comprises a longitudinal cohort containing 132 participants with CD (n = 67), UC (n = 38), and non-IBD controls (n = 27) who were followed for up to one year each. Taxonomic and functional profiles for HMP2 metagenomes (MGX), metabolomes (MBX), and host transcriptomes (HTX) were downloaded from http://ibdmdb.org in July 2020. These were based on 1,595 MGX samples, 546 MBX samples from 106 subjects (CD, n = 50; UC, n = 30; non-IBD, n = 26), and 254 HTX samples from 90 subjects (CD, n=43; UC, n=25; non-IBD, n = 22). MGX samples had been previously profiled for microbial taxonomic composition using MetaPhlAn v2.6.0^45^ and for UniRef90^46^-level gene functional content using HUMAnN v2.11.0^47^. MGX and MBX samples were strictly matched for multi-omic association if they were derived from the same subject and sampling time point (yielding 461 samples from 106 participants: 50 with CD, 30 with UC, and 26 non-IBD controls). MGX and HTX samples were matched more leniently to compensate for the smaller total number of HTX samples. Specifically, we considered the first pair of MGX:HTX samples from each subject that were separated by no more than 2 weeks (yielding 71 samples from 71 individuals: 33 with CD, 21 with UC, and 17 non-IBD controls).

### Identifying differentially abundant metabolites

We used separate statistical models and definitions of disease activity when determining metabolites’ DA status in the PRISM and HMP2 cohorts owing to their cross-sectional vs. longitudinal designs. Specifically, PRISM subjects classified as having CD or UC were compared with non-IBD controls, whereas HMP2 subjects were compared between “active” (dysbiotic) and “inactive” (non-dysbiotic) states within individual time courses as described previously^29^. Prior to statistical model fitting, gut metabolome profiles of PRISM and HMP2 subjects were 1) median-normalized to reduce technical sample-to-sample variation; 2) prevalence-filtered to remove low-confidence features (requiring > 30% non-zero values); and 3) log-transformed for variance-stabilization (replacing zero values with half the smallest non-zero measurement on a per-feature basis).

Differential abundance over disease phenotype (diagnosis) was determined within the PRISM cohort by evaluating the following linear model for each metabolite in base R version 4.0.2:

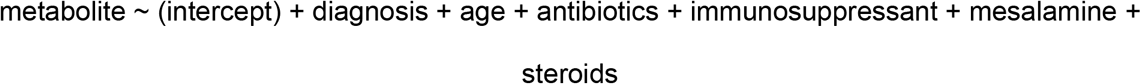

Diagnosis was coded as a categorical variable (CD, UC, non-IBD control) with non-IBD control as the reference state. Age was coded as a continuous covariate and four medication exposures (use of antibiotics, immunosuppressants, mesalamine, and steroids) were coded as binary covariates. Individual medications within these broad classes (e.g. specific antibiotic treatments) were insufficiently numerous to merit separate coding.

DA status over disease activity (dysbiosis) was determined within the longitudinally sampled HMP2 cohort by evaluating the following linear mixed-effects model for each metabolite using R’s nlme package:

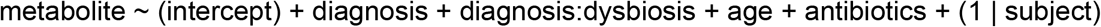

Diagnosis was coded as described above in the context of the PRISM cohort. Dysbiosis within diagnosis (diagnosis:dysbiosis) was used to determine DA status, with per-diagnosis non-dysbiotic samples serving as the reference state. Age at study consent was included as a continuous covariate and per-sample antibiotics exposure as a binary covariate. A per-subject random effect was included to compensate for repeated sampling and to “absorb” potential confounders that were invariant over subjects (e.g. recruitment site).

Model coefficients of the diagnosis (PRISM) and diagnosis:dysbiosis (HMP2) terms were interpreted as DA effect sizes, and their associated two-tailed *p*-values were used to determine statistical significance (Wald’s test). Where applicable, simultaneously derived *p*-values were adjusted for multiple hypothesis testing using the Benjamini-Hochberg FDR method (**Table S6**; **Fig. 4d**).

### Identifying differentially abundant microbial features and metabolite associations

3α- and 3β-HSDH homologs were identified based on mapping query sequences to known protein family clusters as defined in UniRef^31^ (release 2014_07). We first identified the UniRef90 annotations (i.e. protein sequences with >90% amino acid identity and >80% coverage) of the genes identified as 3α- and/or 3β-HSDHs (Elen_0690 and Elen_1325 from *E. lenta*; Rumgna_02133 and Rumgna_00694 from *R. gnavus* ATCC 29149; BF3538 from *B. fragilis* NCTC 9343). We refer to these as “query UniRef90s.” To identify homologs of a given query UniRef90s, we collected all UniRef90 families belonging to the same UniRef50 family as the query (i.e. a set of proteins expected to have >50% identity and high coverage of the query) (**Table S9**). We then estimated the per-sample abundance of 3α-HSDH and 3β-HSDH homologs in HMP2 metagenomes by summing over the abundances of homologous UniRef90 sequences (which had been pre-computed using HUMAnN) (**Table S9; Fig. 4f; Fig. S9**).

We tested 3α-/3β-HSDH homologs for differential abundance over dysbiosis states following a very similar approach to the one introduced above in the context of HMP2 metabolomics. Gene abundance values were similarly zero-smoothed and log-transformed prior to linear model fitting within the MaAsLin 2 package^46^. The same random effects model formulation applied to HMP2 metabolomics was applied here within MaAsLin. We additionally applied this modeling approach to the abundances of microbial species for which corresponding strains had been found to express 3α-/3β-HSDH activity in vitro.

In order to associate microbial features (genes and species) with metabolites of interest within the HMP2 dataset, we computed Spearman correlations between the gene and metabolites’ residual abundances from the previously described linear models (**Table S10; Fig. S10**). This procedure helps to identify correlation between features that cannot be explained by the confounding effects of covariates included in the models. Conversely, a metabolite and gene that both correlate strongly with dysbiosis would be expected to also correlate with one another in the raw data, but this would not be suggestive of a direct link between their changing abundances. Here, module residuals reflect variation in gene and metabolite abundance after subtracting differences due to dysbiosis (in addition to diagnosis, age, antibiotics use, and per-subject variation), and therefore any remaining correlation cannot be attributed to those variables. Two-tailed *p*-values associated with these Spearman correlation coefficients were subjected to FDR correction following the procedures introduced above for model coefficients.

### Identifying differentially abundant host transcripts and metabolite associations

Using paired MBX and HTX samples from the HMP2 dataset, we identified human genes that were differentially expressed (DE’ed) with respect to diagnosis (note: because this analysis considers only one HTX sample per HMP2 subject, we focus on per-subject diagnosis as a phenotype rather than per-sample dysbiotic state) (**Table S7-8; Fig. 4e**). We performed initial normalization on raw sample-by-gene HTX count data using the voom method implemented in R’s limma package^48,49^. We then used the normalized counts as a basis for linear modeling within MaAsLin 2 to detect differential gene expression:

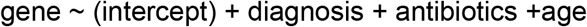

That is, the transformed abundance of each gene was modeled as a function of diagnosis, consent age, and antibiotics use as defined above for the HMP2 cohort. The coefficient of the diagnosis term and its FDR-adjusted *p*-value were used to determine the effect size and statistical significance of potential differential expression. Note that DE analysis was carried out in MaAsLin 2 rather than limma voom itself 1) for consistency with other linear modeling analyses in this work and 2) to enable export of model residuals for multi-omic correlation analysis. From the subset of human DE’ed genes with FDR-adjusted *p* < 0.25, we selected a subset that were previously identified as T_H_17-related^29,30^. We then followed the approach outlined above in the context of microbial features to associate residual expression of these human genes with residual metabolite expression using Spearman correlation.

## Data availability

The 16S amplicon and RNA-Seq datasets are available through NCBI under accession number PRJNA675599 and GSE161977, respectively.

### Code availability

The software packages used in this study are free and open source. Source code for ElenMatchR is available at github.com/turnbaughlab/ElenMatchR. MaAsLin2 is available via http://huttenhower.sph.harvard.edu/maaslin as source code and installable packages. The R package limma is available from https://www.bioconductor.org/packages/release/bioc/html/limma.html. Analysis scripts employing these packages are available from the authors upon request.

## Acknowledgments

We thank members of the Devlin, Huh, and Clardy labs (Harvard Medical School-HMS) for helpful discussions. We thank the HMS ICCB-Longwood Screening Facility, BPF Genomics Core Facility at Harvard Medical School for their expertise and instrument support, N. Lee, J. Vasquez, C. Powell, and M. Henke for technical support and advice, and M. Trombly and S. Blacklow for critical reading of the manuscript. We are grateful to the human patients who participated in the human stool screen, PRISM and HMP2 studies. This work was supported by National Institutes of Health grants R01 DK110559 (J.R.H.), R01AR074500 (P.J.T.), U54DE023798 (C.H.), R24DK110499 (C.H.), and MIRA R35 GM128618 (A.S.D.), a Harvard Medical School Dean’s Innovation Grant in the Basic and Social Sciences (A.S.D. and J.R.H.), a John and Virginia Kaneb Fellowship (A.S.D.), a Harvard Medical School Christopher Walsh Fellowship (L.Y.) and a Wellington Postdoctoral Fellowship (L.Y.). J.E.B was the recipient of a Natural Sciences and Engineering Research Council of Canada Postdoctoral Fellowship and is supported by the National Institute of Allergy and Infectious Diseases (K99AI147165). P.J.T. is a Chan Zuckerberg Biohub investigator. The computations in this paper were run in part on the FASRC Cannon cluster supported by the FAS Division of Science Research Computing Group at Harvard University. Figure panels in Extended Data Fig 1b, c were created using BioRender.

## Author contributions

J.R.H. and A.S.D. conceptualized the study. D.P., L.Y., J.R.H., and A.S.D. conceived the project and designed the experiments. D.P. performed mouse experiments, in vitro T cell and reporter assays. L.Y. performed human isolate screen, bacterial in vitro culture experiments, and bile acid profiling. G.D.D. performed HSDH enzyme characterization. Y.Z. and S.B. performed the bioinformatics analyses. E.A.F. and C.H. supervised the computational analyses. J.A.P. and C.C. performed LCA derivative identification in PRISM and HMP2 metabolomics. E.K. performed T cell RNA-Seq analysis. J.E.B. performed comparative genomics on *E. lenta*. C.K.R. and M.R.K. synthesized some of the bile acid derivatives. J.E.B. and P.J.T., supervised the *E. lenta* human isolate studies. H.V. and R.J.X. provided bacterial strains and technical support. R.L. provided the patient stool samples. D.P., L.Y., Y.Z., S.B., G.D.D., E.A.F., J.R.H., and A.S.D. wrote the manuscript, with contributions from all authors.

## Extended Data Figures

**Extended Data Fig. 1.**
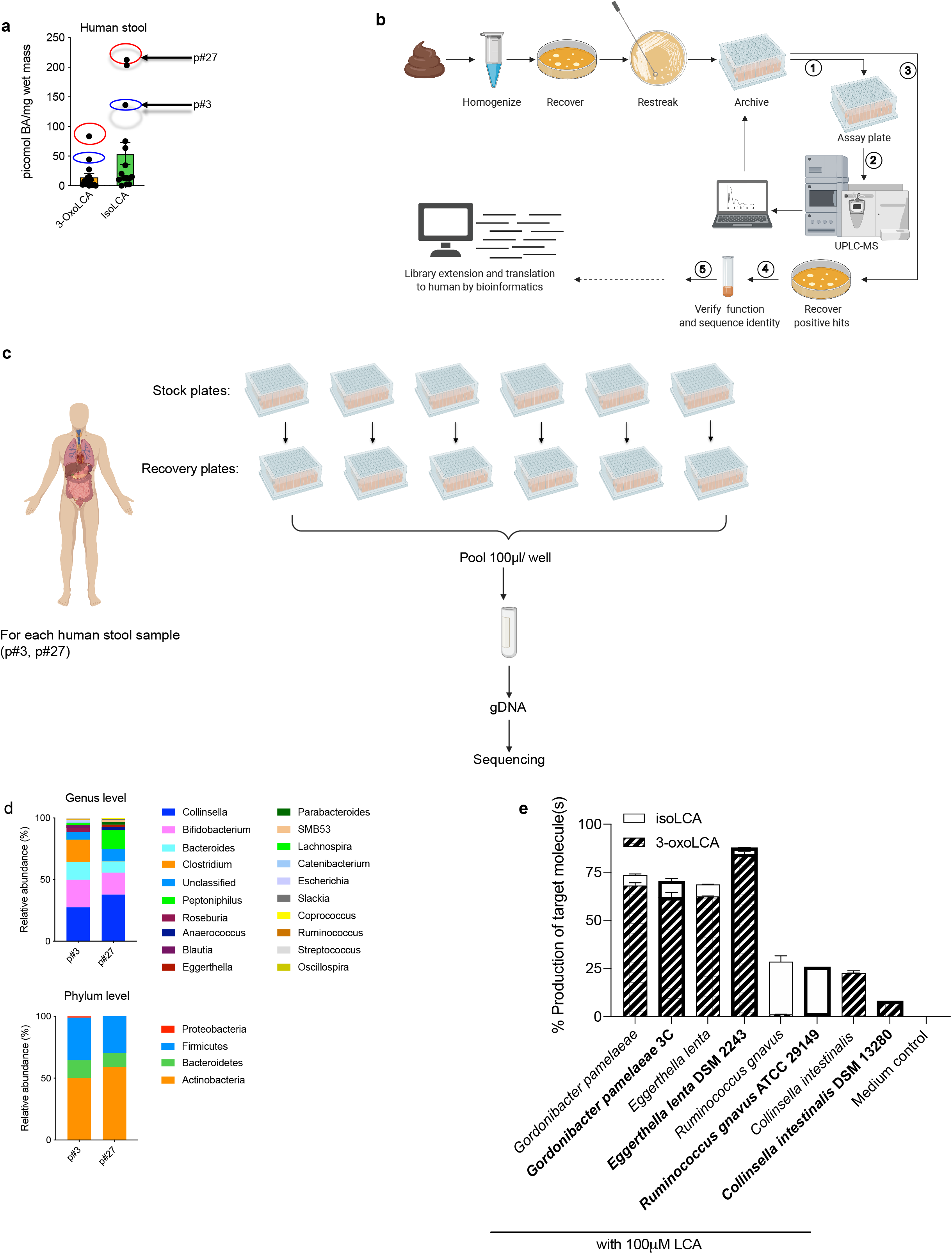
3-oxoLCA biosynthetic pathway and microbial diversity from the human screen. **a**, Quantification of 3-oxoLCA and isoLCA in stool samples from patients after faecal microbiota transplant (FMT) (n = 15). Stool samples from patient p#3 (3-oxoLCA: 44 picmol/mg, isoLCA: 136 picmol/mg) and patient p#27 (3-oxoLCA: 83 picmol/mg, isoLCA: 213 picmol/mg) were used to screen for 3-oxoLCA producers. **b**, Schematic of the screen for bacterial producers of the LCA metabolite 3-oxoLCA from human stool samples. In total, 990 bacterial colonies were isolated, restreaked, and archived from two human stool samples. ① Replicate plates (assay plates) were then used for the screen. ② Individual isolates were incubated anaerobically with LCA (100 μM) (see **Fig. 1b**) or 3-oxoLCA (100 μM) (see **Fig. 2b**) for 48 hours. Cultures were harvested, acidified, extracted, and bile acid metabolites were quantified by UPLC-MS. ③ Positive hits containing 3-oxoLCA were re-selected from the archived stock plates, and recovered on new plates. ④ Activity was verified and each producer species was identified by full-length 16S rRNA sequencing. Finally, bacterial enzymes responsible for the LCA metabolite production were identified (see **Fig. 3**), and ⑤ corresponding genes were utilized as query sequences in BLASTP searches for novel putative bacterial producers and enzymes. **c**, Sample preparation workflow for the determination of cultured bacteria from the human stool sample screen. For each patient, individual isolates were recovered and cultured for 48 hours. These isolates were then pooled together, and genomic DNA was extracted from the pooled pellet. Illumina® MiSeq sequencing on the V3 and V4 hypervariable regions of 16S rRNA was then performed. **d**, Genus and phylum-level microbial community composition for each human stool sample. **e**, 3-oxoLCA and/or isoLCA production was verified in the type strains of a subset of 3-oxoLCA-producing human isolates (n = 3 biological replicates per group).

**Extended Data Fig. 2.**
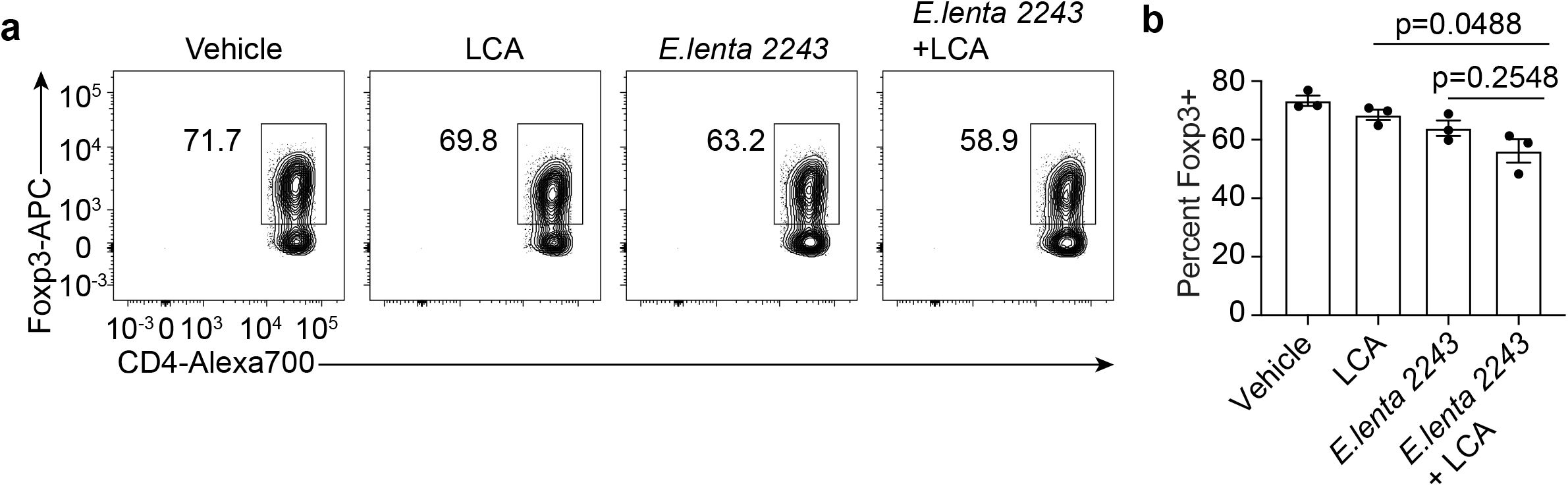
Supernatants from LCA metabolite-producing bacteria do not affect T_reg_ cell differentiation in vitro. **a, b**, Representative FACS plots (**a**) and population frequencies (**b**) of CD4+ T cells, cultured under T_reg_ polarization conditions in vitro are presented. Bacterial culture supernatants were added 18 hours after TCR activation (n = 3 biologically independent samples per group. Data are mean ± SEM, followed by one-way ANOVA with Tukey’s multiple comparison test).

**Extended Data Fig. 3.**
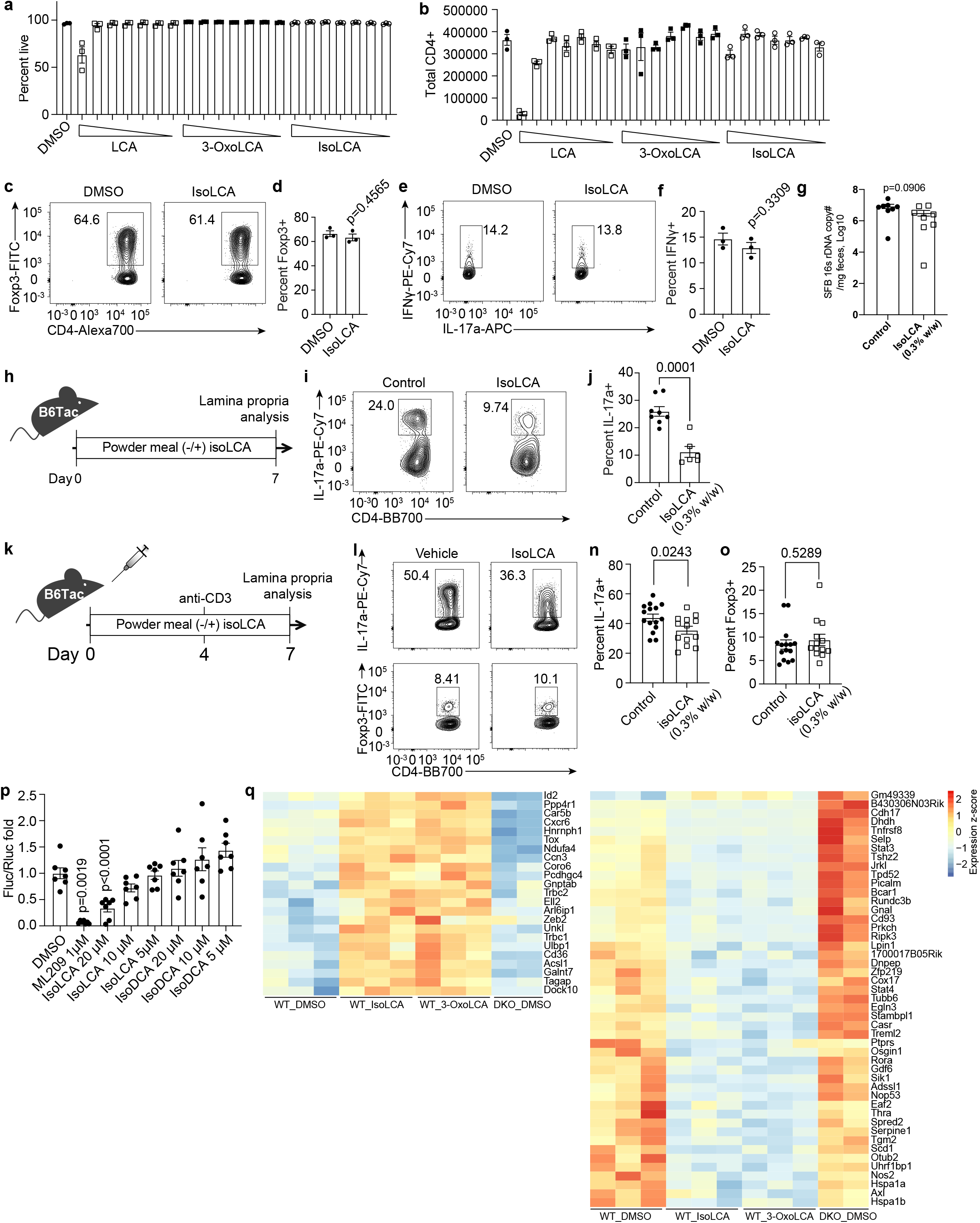
IsoLCA neither affects T cell viability nor inhibits T_reg_ and T_H_1 cell differentiation in vitro. **a, b**, IsoLCA does not reduce T cell viability or proliferation. Percentages of viable cells (**a**) and total cell numbers (**b**) at the end of T cell culture under T_H_17 polarization conditions in the presence of LCA, 3-oxoLCA, or isoLCA. The same concentrations were used as listed in **Fig. 2b** (40, 20, 10, 5, 2.5, 1.25 and 0.625 μM) (n = 3 biologically independent samples, data are mean ± SEM, one-way ANOVA with Dunnett’s multiple comparisons). **c-f**, IsoLCA does not affect T_reg_ or T_H_1 cell differentiation in vitro. Flow cytometry and quantification of intracellular staining for Foxp3 (**c, d**) or IFN-γ (**e, f**). Mouse naive CD4 T cells from wild-type B6Jax mice were cultured under T_H_1- or T_reg_-polarizing conditions and DMSO or isoLCA was added 18 hours after TCR activation (n = 3 biologically independent samples per condition, data are mean ± SEM, followed by unpaired t-test with two-tailed *p*-value). **g**, SFB colonization measured by qPCR analysis in **Fig.2 d-g**, calculated as SFB 16s rRNA copy number (n = 8 mice per group, pooled from two experiments, data are mean ± SEM, followed by unpaired t-test with two-tailed *p*-value). **h–j**, Experimental scheme of Th17 induction by SFB (**h**), representative FACS plots (**i**) and population frequencies of T_H_17 cells (**j**), isolated from the ileal lamina propria of control or isoLCA-treated mice (n = 8 mice for control, n=6 mice for isoLCA-treated groups, pooled from two experiments). B6 Tac mice were fed a control or a isoLCA (0.3% w/w)-containing diet for 7 days (data are mean ± SEM, by unpaired t-test with two-tailed *p*-value). **k–o**, Experimental scheme of anti-CD3 experiment (**k**), representative FACS plots (**l, m**) and population frequencies of T_H_17 (**n**) and T_reg_ cells (**o**) of the ileal lamina propria of control or isoLCA-treated mice (n = 15 mice for control, 13 mice for isoLCA-treated groups, pooled from three experiments.). B6 Tac mice were intraperitoneally injected with anti-CD3 and fed a control diet or isoLCA-containing (0.3% w/w) diet during the experiments (data are mean ± SEM, by unpaired t-test with two-tailed *p*-value). **p**, RORγt luciferase reporter assay in HEK293 cells, treated with a synthetic RORγ inhibitor ML209 (1 μM), isoLCA (20 μM, 10 μM, 5 μM), isoDCA (20 μM, 10 μM, 5 μM) or DMSO. The fold ratios of firefly luciferase (FLuc) to Renilla luciferase (RLuc) activity is presented on the y-axis. DMSO-treated group set to 1 (n = 7 independent transfections per group, pooled from two experiments. Data are mean ± SEM, one-way ANOVA with Dunnett’s multiple comparisons, vehicle set as control). **q**, Heat map representing 70 genes that are commonly regulated by isoLCA and 3-oxoLCA, but not by RORα and RORγt (WT: wild-type, n = 3 mice per condition, DKO: RORα/RORγ double deficient cells, n=2 mice per condition).

**Extended Data Fig. 4.**
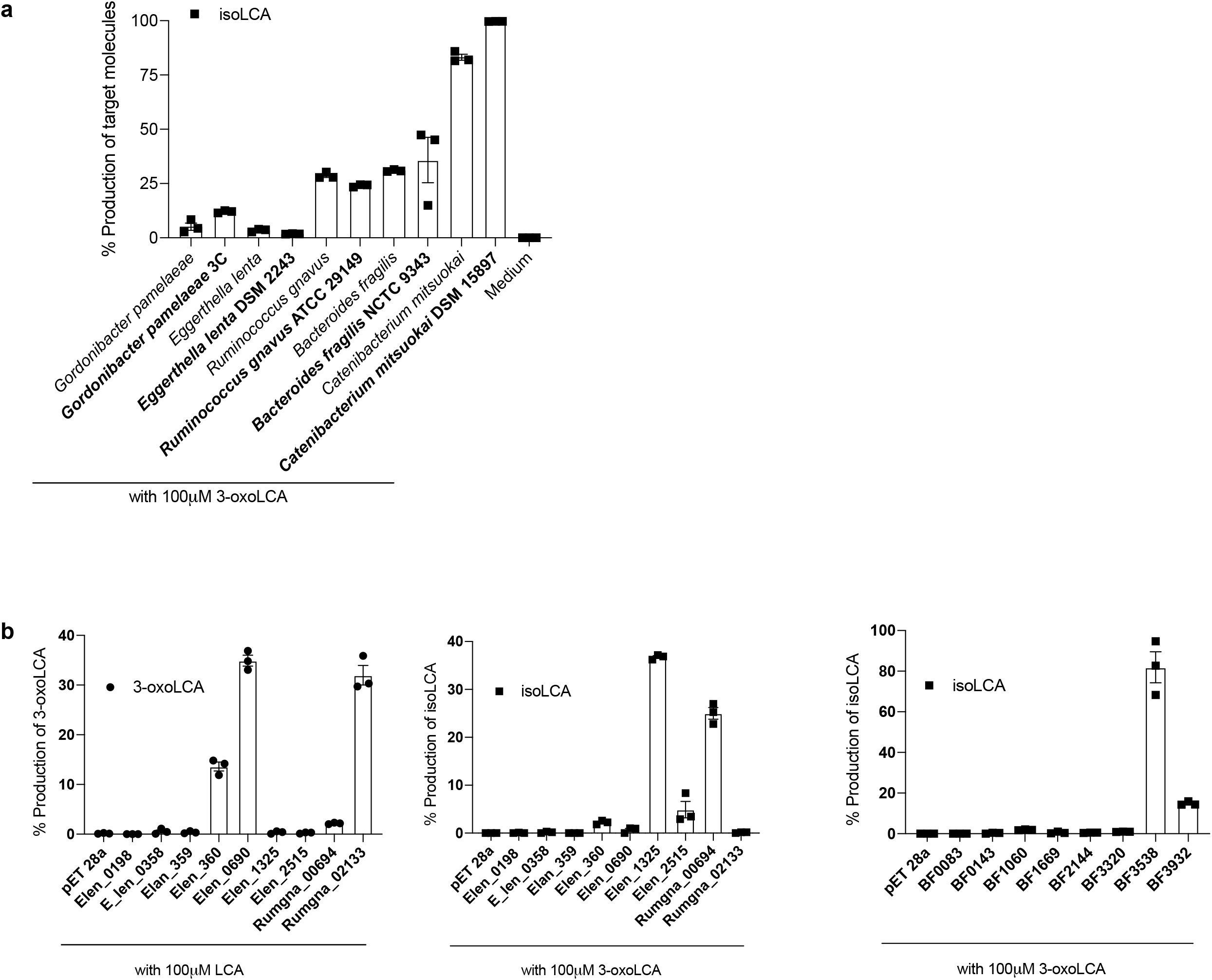
Screen of the candidate HSDH enzymes from gut bacteria. **a**, isoLCA production from 3-oxoLCA (100 μM) was verified in the type strains of a subset of isoLCA-producing human isolates (n = 3 biological replicates per group). **b**, Results of lysis assay in which the *E. lenta* DSM 2243 (Elen), *R. gnavus* ATCC 29149 (Rumgna), and *B. fragilis* NCTC 9343 (BF) candidate HSDH enzymes were expressed in *E. coli* BL21 pLysS and their ability to convert LCA to 3-oxoLCA (left, 3α-HSDH activity) and 3-oxoLCA to isoLCA (middle and right, 3β-HSDH activity) was analyzed by UPLC-MS. All results are reported as percent conversion of the target product (n = 3 biological replicates per group).

**Extended Data Fig. 5.**
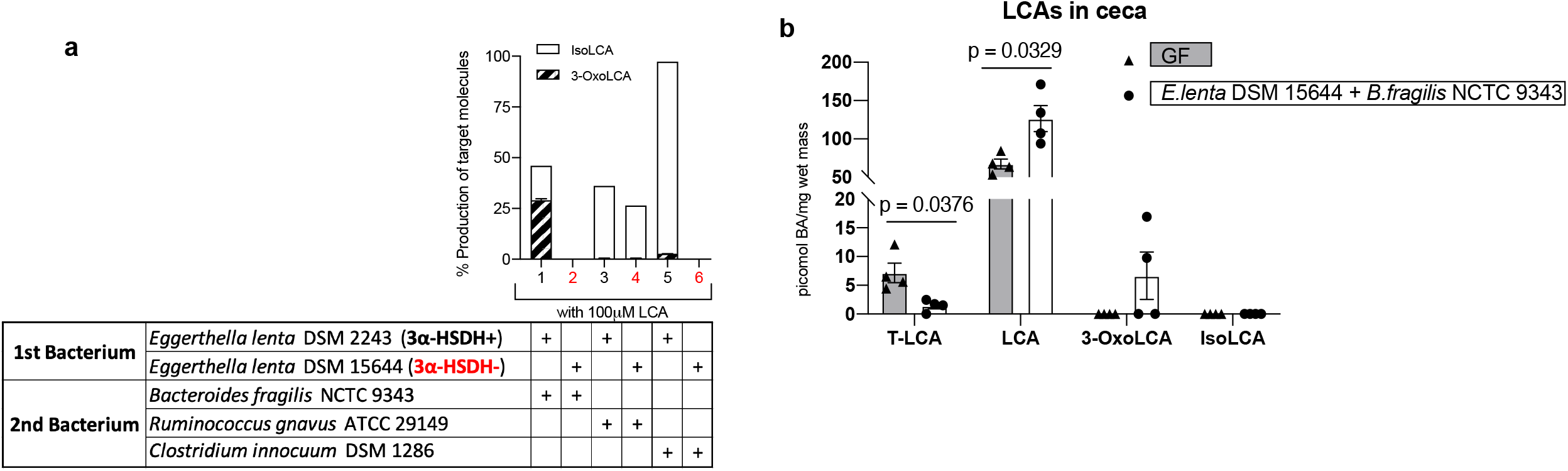
Production of 3-oxoLCA and isoLCA by in vitro co-culture and in vivo colonization. **a**, Production of 3-oxoLCA and isoLCA from LCA (100 μM) by co-cultures of human gut bacteria types strains in vitro. *E. lenta* DSM 2243 was co-cultured with a second bacterium (*B. fragilis* NCTC 9343, *R. gnavus* ATCC 29149, or *C. innocuum* DSM 1286. As a negative control, the *E. lenta* human isolate (*E. lenta* DSM 15644) which lacks a 3α-HSDH homolog was paired with the same bacteria (in red numbers). **b**, Levels of LCA derivatives in cecal contents of gnotobiotic mice co-colonized with *E. lenta* DSM 15644 and *B. fragilis* NCTC 9343. LCA (0.3% w/w) in mouse chow was provided and cecal contents were analyzed by UPLC-MS (n=4 mice per group from a single experiment, data are mean ± SEM, followed by Welch’s t tests).

**Extended Data Fig. 6.**
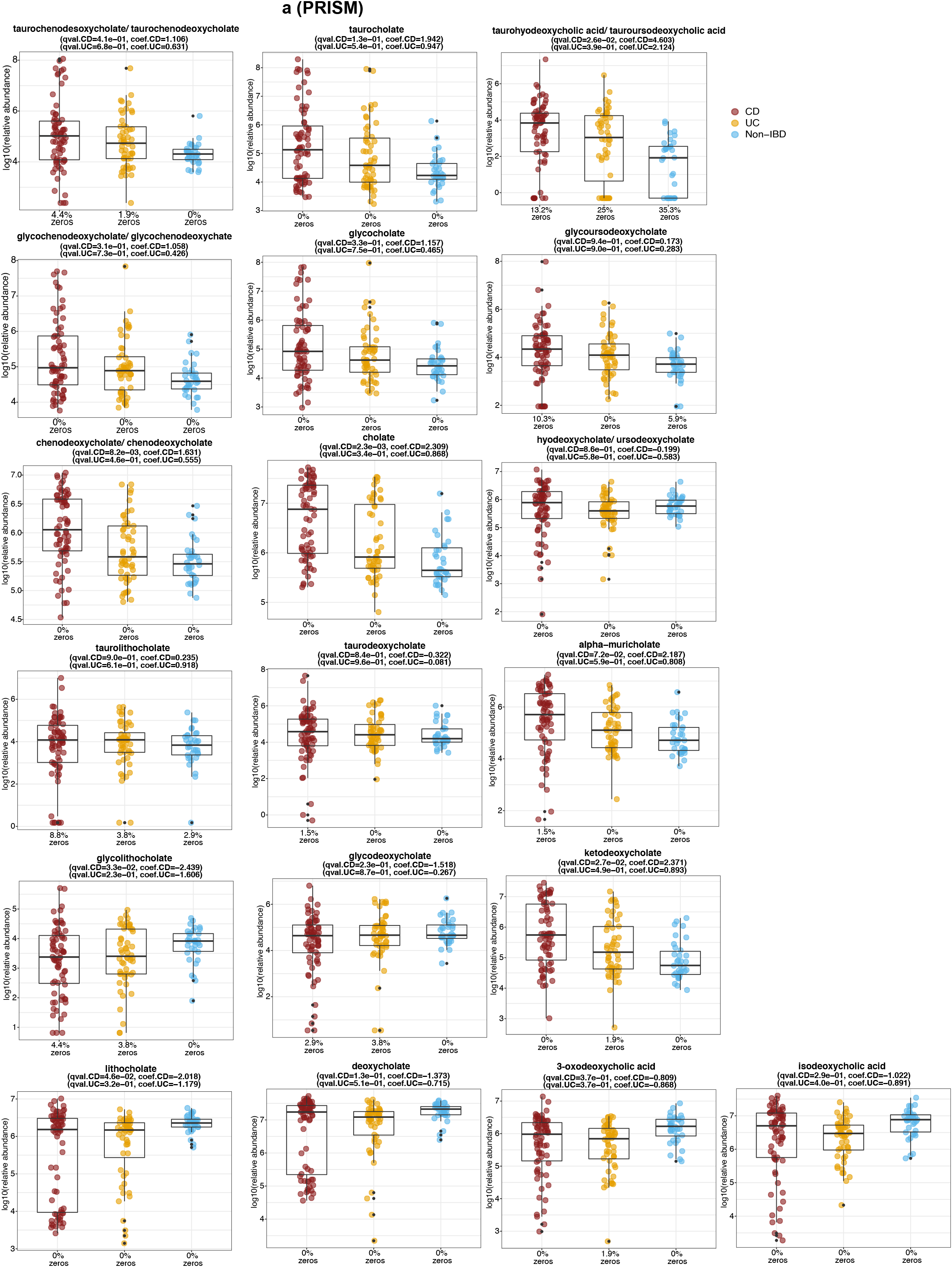
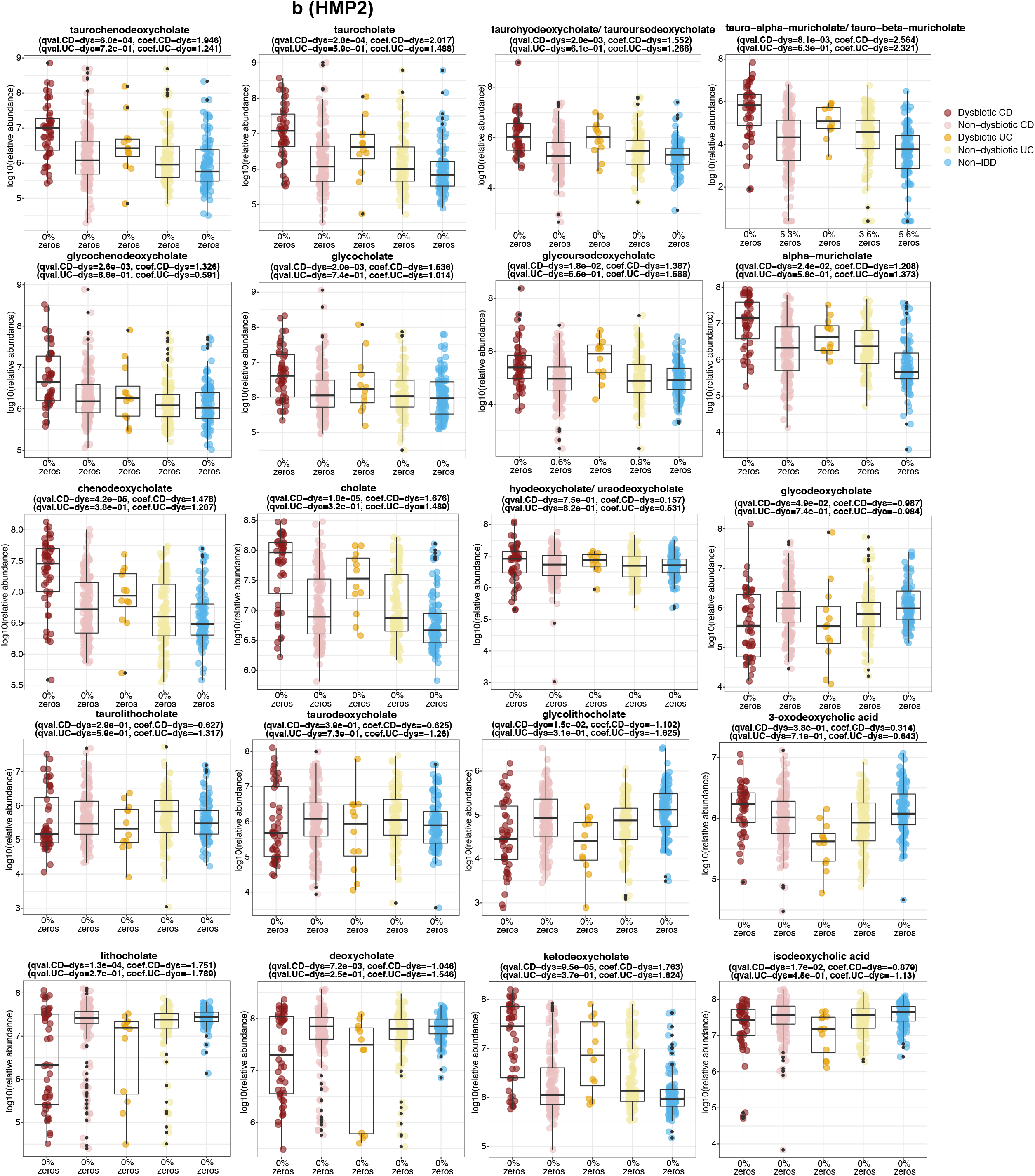
Levels of bile acid metabolites detected in human cohorts. **a, b** Abundances of identifiable bile acids in healthy subjects and Crohn’s disease (CD) patients from PRISM (**a**) and HMP2 cohorts (**b**). Bile acid levels were not universally decreased in CD patients, indicating that decreased levels of LCA, 3-oxoLCA, and isoLCA were not due to lower levels of all bile acids in these cohorts. Boxplots show median and lower/upper quartiles with outliers outside of boxplot ‘whiskers’ (indicating the inner fences of the data). The percentage of zeros in each condition are added as x-axis tick labels. See **Table S6** for full results.

**Extended Data Fig. 7.**
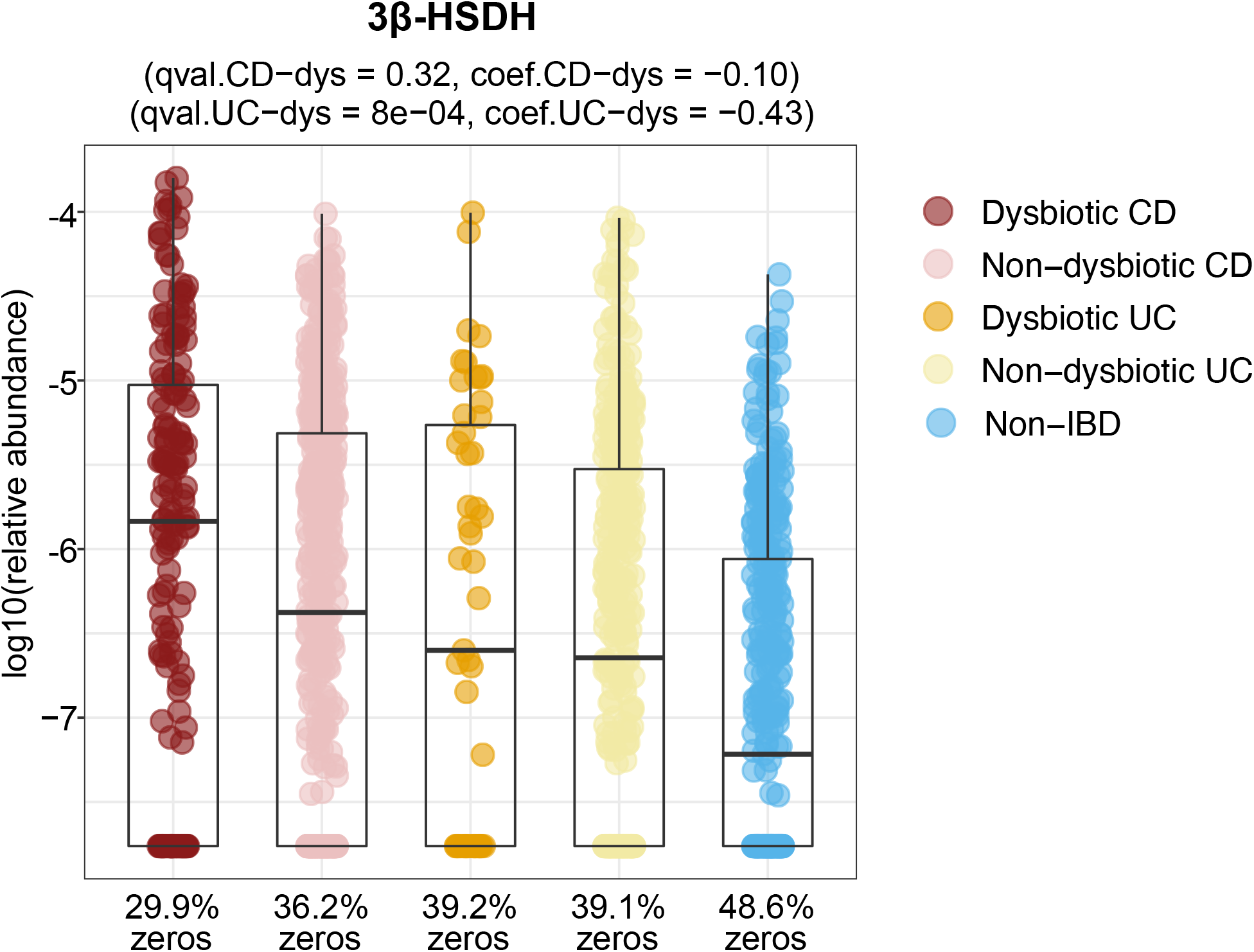
Relative abundance distributions of differentially abundant 3β-HSDH homologs profiled from HMP2 metagenomes. The statistical results came from the linear-mixed effects model with coefficient for dysbiosis state within IBD phenotype (CD and UC) and FDR-adjusted *p*-values < 0.05 (**Methods;** n = 1,595 samples from 130 subjects). Boxplots show median and lower/upper quartiles with outliers outside of boxplot ‘whiskers’ (indicating the inner fences of the data). The percentage of zeros in each condition are added as x-axis tick labels. See **Table S9** for full results.

