## Supplementary Information for "Human gut bacteria produce T_H_17-modulating bile acid metabolites"

##### IsoLCA Synthesis Procedure.

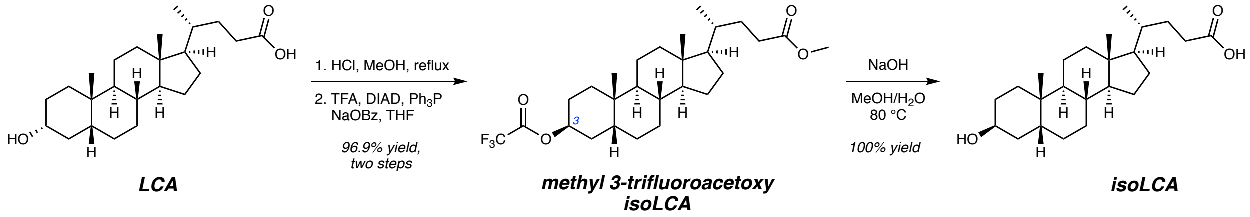


**IsoLCA.** Isolithocholic acid (isoLCA) was prepared from lithocholic acid (LCA) by carboxylic acid protection, then C3 alcohol inversion[1], followed by ester cleavage. Methanol (125 mL, 0.25 M) and HCl (622 µL of 1 M solution in methanol, 0.622 mmol, 0.02 equivalents) were added to a 500 mL round-bottom flask containing lithocholic acid (11.71 g, 31.10 mmol, 1 equivalent) and a stir bar. A reflux condenser was affixed to the flask and the contents were brought to reflux by immersion in a 90 °C oil bath with mixing. Upon completion by TLC analysis (1:1 hexanes/ethyl acetate; *p*-Anisaldehyde; overnight), the flask was cooled to room temperature and concentrated in vacuo. The resulting solid was further dried under high vacuum to afford methyl lithocholate as a free-flowing white solid (12.143 g, 31.087 mmol, 100% yield). $R_f$ = 0.35 (2:1 hexanes/ethyl acetate; *p*-Anisaldehyde); mp = 130–132 °C (lit.[3] mp 130–132 °C); $^1$H NMR (600 MHz, $CDCl_3$): δ 3.66 (s, 3H), 3.62 (td, *J* = 10.9, 5.3 Hz, 1H), 2.35 (ddd, *J* = 15.4, 10.2, 5.1 Hz, 1H), 2.22 (ddd, *J* = 15.7, 9.9, 6.5 Hz, 1H), 1.95 (dt, *J* = 12.4, 3.2 Hz, 1H), 1.88–1.73 (m, 5H), 1.68–1.64 (m, 1H), 1.59–1.55 (m, 1H), 1.53–1.49 (m, 1H), 1.43–1.19 (m, 12H), 1.16–1.03 (m, 5H), 0.97 (td, *J* = 14.2, 3.4 Hz, 1H), 0.92 (s, 3H), 0.91 (d, *J* = 6.5 Hz, 3H), 0.64 (s, 3H); HRMS (DART+) *m/z*: $[M + NH_4]^+$ calculated for $C_{25}H_{46}O_3N$ 408.3478, found 408.3468; $[\alpha]_D^{21.3}$ +30.62 (*c* 1.07, $CH_2Cl_2$). All other compound data are consistent with reported values[2,3].

Tetrahydrofuran (105 mL, 0.3 M) was added to a flame-dried 500 mL round-bottom flask charged with methyl lithocholate (12.332 g, 31.571 mmol, 1 equivalent) and a stir bar under argon. To this colorless solution was added diisopropyl azodicarboxylate (DIAD: 6.84 mL, 34.73 mmol, 1.1 equivalents) and trifluoroacetic acid (TFA: 2.66 mL, 34.73 mmol, 1.1 equivalents), and the flask was placed in a room temperature water bath (T = 21 °C). Addition of triphenylphosphine ($Ph_3P$: 9.108 g, 34.73 mmol, 1.1 equivalents) to this bright orange solution resulted in a rapid color fading. Sodium benzoate (NaOBz: 5.005 g, 34.73 mmol, 1.1 equivalents) was then added and the white suspension was thoroughly mixed at room temperature. Upon completion by TLC analysis (2:1 hexanes/ethyl acetate; *p*-Anisaldehyde; ca. 4 h), hexanes (65 mL) were added. After mixing for at least 15 min, the reaction was filtered through a medium porosity Buchner funnel and the cloudy filtrate was concentrated in vacuo to yield an off-white solid. This crude solid (comprised of desired product, $Ph_3PO$, and reduced DIAD) was dissolved in $CH_2Cl_2$ (ca. 150 mL), dry-loaded onto a mixture of Celite (15 g) and $SiO_2$ (90 g), and purified by flash chromatography (6 x 21 cm $SiO_2$, 10% ethyl acetate in hexanes; collect fractions in 125–250 mL Erlenmeyer flasks) to afford methyl 3-trifluoroacetoxy isolithocholate as a white crystalline solid (14.885 g, 30.589 mmol, 96.9% yield). $R_f$ = 0.37 (9:1 hexanes/ethyl acetate; *p*-Anisaldehyde); mp 142–145 °C; $^1$H NMR (600 MHz, $CDCl_3$): δ 5.30 (br s, 1H), 3.66 (s, 3H), 2.35 (ddd, *J* = 15.4, 10.2, 5.2 Hz, 1H), 2.22 (ddd, *J* = 15.8, 9.9, 6.5 Hz, 1H), 2.05 (ddd, *J* = 15.7, 13.5, 2.9 Hz, 1H), 1.99–1.96 (m, 1H), 1.93–1.77 (m, 3H), 1.73–1.53 (m, 6H), 1.46–1.26 (m, 9H), 1.21–1.00 (m, 6H), 0.98 (s, 3H), 0.91 (d, *J* = 6.5 Hz, 3H), 0.65 (s, 3H).; $^{13}$C{$^1$H} NMR (151 MHz, $CDCl_3$): δ 174.9, 157.1 (q, *J* = 41.6 Hz), 114.8 (q, *J* = 286.2 Hz), 76.7, 56.7, 56.1, 51.6, 42.9, 40.3, 40.1, 37.3, 35.7, 35.5, 35.0, 31.2, 31.1, 30.5, 30.4, 28.3, 26.4, 26.2, 24.8, 24.3, 23.8, 21.2, 18.4, 12.2; $^{19}$F NMR (376 MHz, $CDCl_3$): δ –75.4 (s); IR (ATR): 2973, 2929, 1777, 1737, 1435, 1213, 1185, 1152, 859 cm$^{-1}$; HRMS (DART+) *m/z*: $[M + NH_4]^+$ calculated for $C_{27}H_{45}O_4F_3N$ 504.3301, found 504.3289; $[\alpha]_D^{22.0}$ +15.95 (*c* 1.08, $CH_2Cl_2$). NMR spectra are shown in **Supplementary Fig. 1, 2** and **3**.

Methanol (146 mL), $H_2O$ (29 mL; 5:1 methanol:$H_2O$, 0.1 M), and NaOH (4.24 g, 106 mmol, 3 equivalents) were added to a 1 L round-bottom flask charged with methyl 3-trifluoroacetoxy isoLCA (17.254 g, 35.312 mmol, 1 equivalent) and a stir bar. A reflux condenser was affixed to the flask, and the contents were placed under argon and warmed in an 80 °C oil bath with mixing. The suspension turned homogeneous over an hour. Upon reaction completion by TLC analysis (2:1 hexanes/EtOAc; *p*-Anisaldehyde; ca. 3 h), the flask was cooled to room temperature and methanol was removed in vacuo to yield a viscous slurry. $H_2O$ (400 mL) was added to this slurry and the contents were sonicated, followed by acidification to pH < 3 with 1 M HCl (ca. 125 mL). The precipitate was sonicated for at least 10 min to ensure a free-flowing solid, then cooled in an ice bath and isolated via vacuum filtration. Washing and further drying of the filter cake (air, then 110 °C oven over night, then high vacuum) yielded isolithocholic acid as a powdery white solid (13.419 g, 35.634 mmol, 100% yield).

***Characterization data[4]:*** $R_f$ = 0.12 (1:1 hexanes/ethyl acetate; *p*-Anisaldehyde); mp 154–158 °C; $^{1}$H NMR (600 MHz, $CD_3OD$): δ 4.03 (p, *J* = 2.7 Hz, 1H), 2.32 (ddd, *J* = 15.2, 9.8, 5.3 Hz, 1H), 2.19 (ddd, *J* = 15.7, 9.5, 6.9 Hz, 1H), 2.01 (dq, *J* = 13.8, 3.0 Hz, 2H), 1.97–1.85 (m, 2H), 1.79 (dddd, *J* = 13.1, 9.8, 6.9, 2.8 Hz, 1H), 1.77–1.72 (m, 1H), 1.60 (dtd, *J* = 10.9, 6.1, 5.5, 3.3 Hz, 1H), 1.56 (ddd, *J* = 14.2, 4.7, 2.5 Hz, 1H), 1.49–1.38 (m, 8H), 1.35–1.26 (m, 4H), 1.22–1.06 (m, 6H), 0.97 (s, 3H), 0.95 (d, J = 6.6 Hz, 3H), 0.69 (s, 3H); $^{13}$C{$^{1}$H} NMR (151 MHz, $CD_3OD$): δ 178.2, 67.8, 58.0, 57.5, 43.9, 41.6, 41.1, 37.8, 37.1, 36.7, 36.2, 34.4, 32.3, 32.01, 31.0, 29.2, 28.5, 27.9, 27.4, 25.3, 24.4, 22.2, 18.8, 12.5; IR (ATR): 3232, 2922, 2864, 1710, 1442, 1217, 1038, 957 cm$^{-1}$; HRMS (DART–) *m/z*: $[M - H]^-$ calculated for $C_{24}H_{39}O_3$ 375.2899, found 375.2908; $[\alpha]_D^{22.4}$ +26.3 (*c* 0.805, $CH_3OH$). NMR spectra are shown in **Supplementary Fig. 4** and **5**.

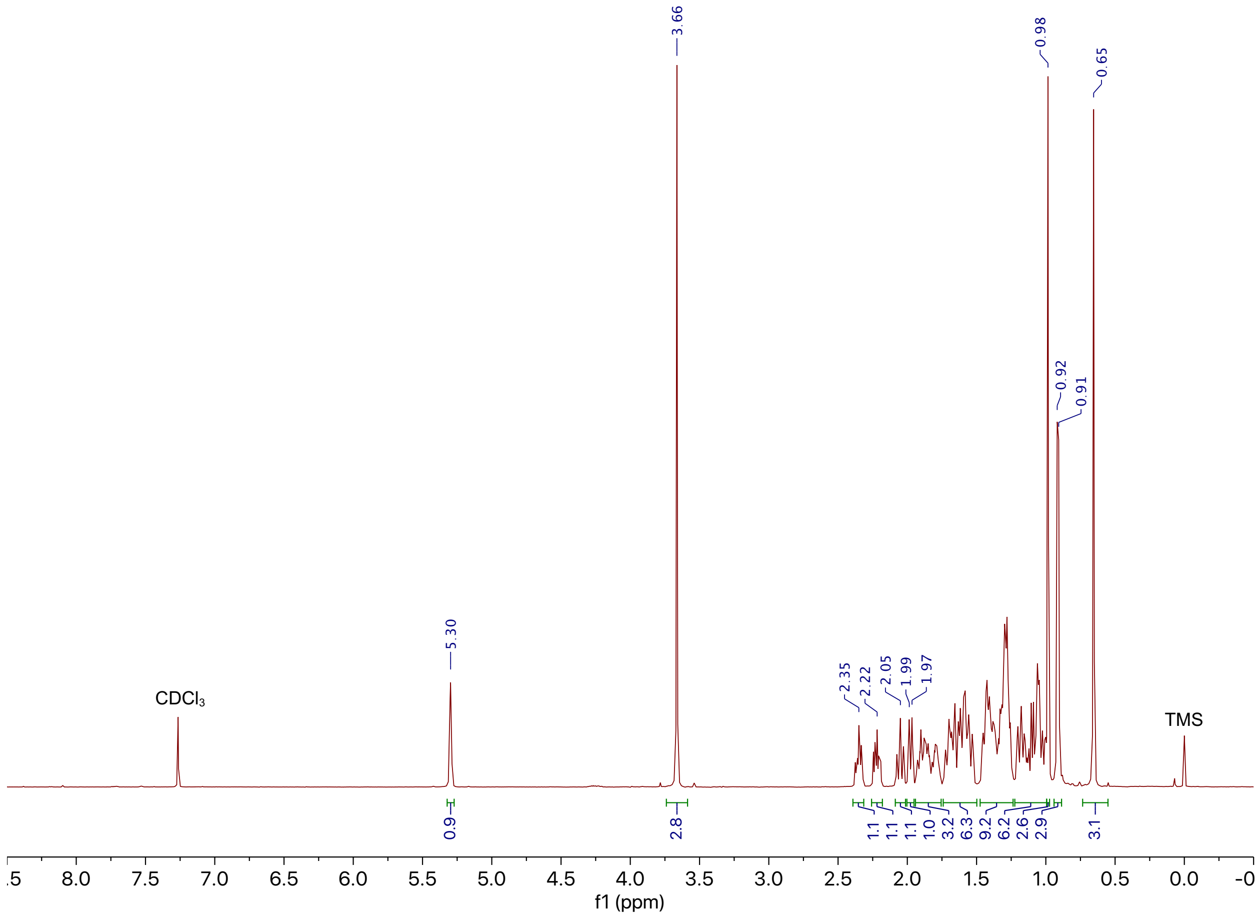


**Supplementary Figure 1. $^1H$ NMR spectrum (600 MHz, $CDCl_3$) of methyl 3-trifluoroacetoxy isolithocholate.** The compound data and spectrum of methyl 3-trifluoroacetoxy isolithocholate are representative of at least six synthesis experiments.

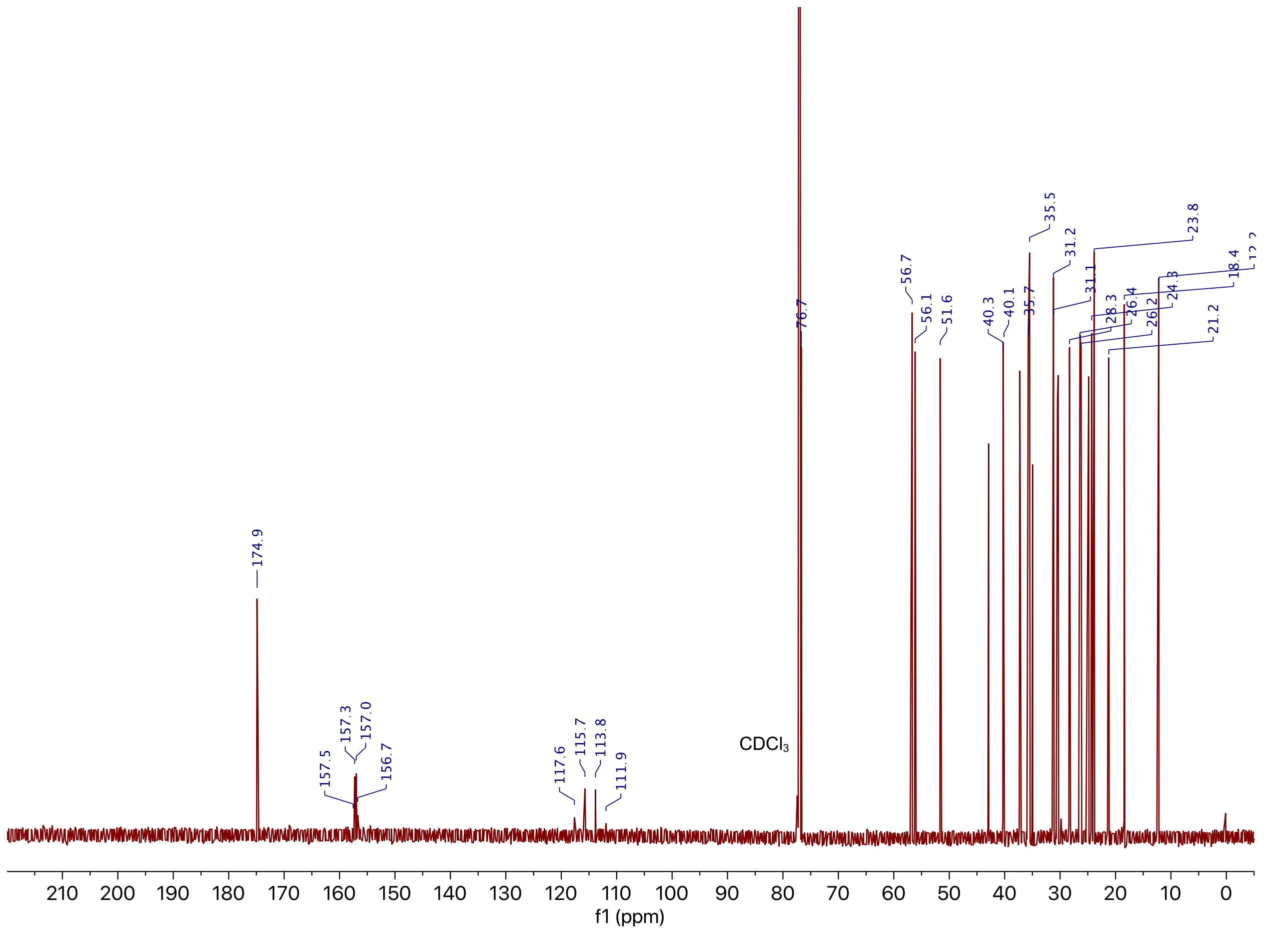


**Supplementary Figure 2. $^{13}C\{^{1}H\}$ NMR spectrum (151 MHz, $CDCl_3$) of methyl 3-trifluoroacetoxy isolithocholate.** The compound data and spectrum of methyl 3-trifluoroacetoxy isolithocholate are representative of at least six synthesis experiments.

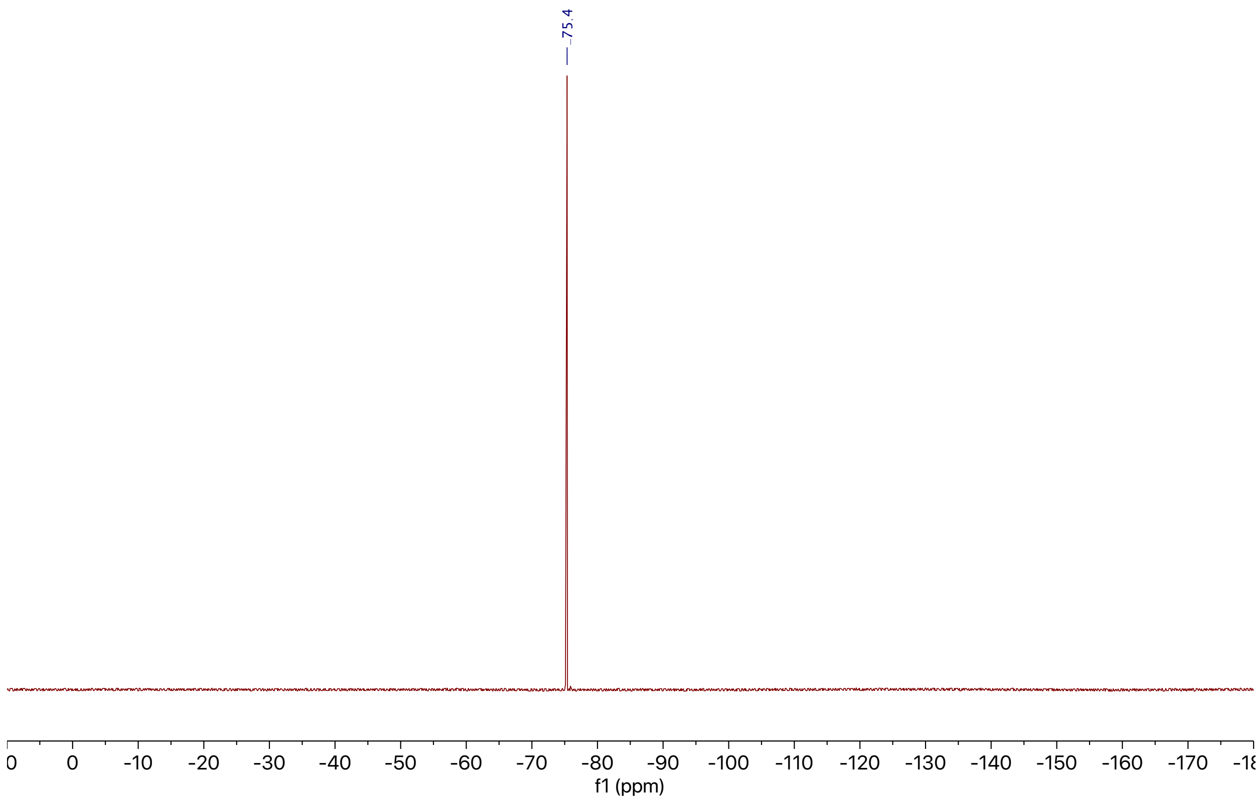


**Supplementary Figure 3. $^{19}F$ NMR spectrum (376 MHz, $CDCl_3$) of methyl 3-trifluoroacetoxy isolithocholate.** The compound data and spectrum of methyl 3-trifluoroacetoxy isolithocholate are representative of at least six synthesis experiments.

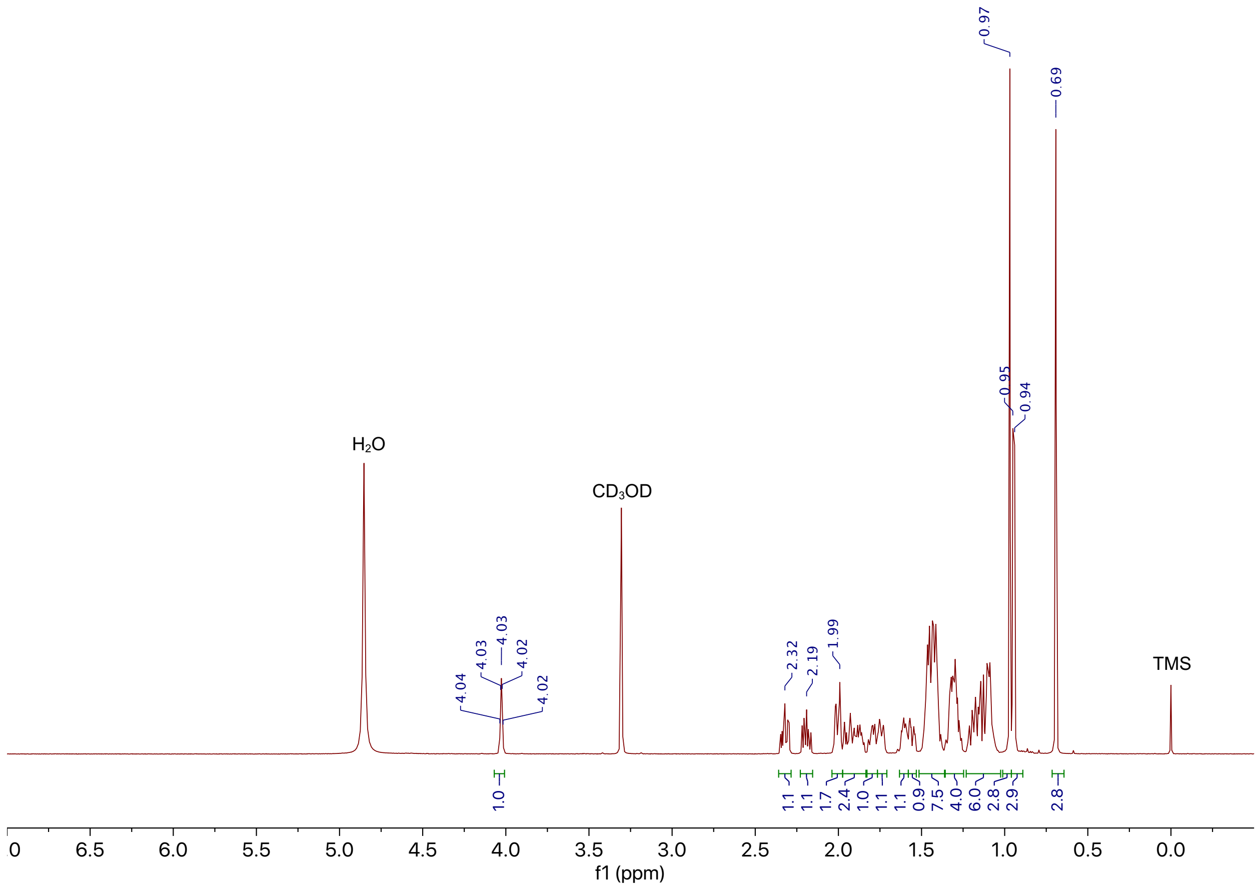


**Supplementary Figure 4. $^1$H NMR spectrum (600 MHz, $CD_3OD$) of isolithocholic acid.** The compound data and spectrum of isoLCA are representative of four synthesis experiments.

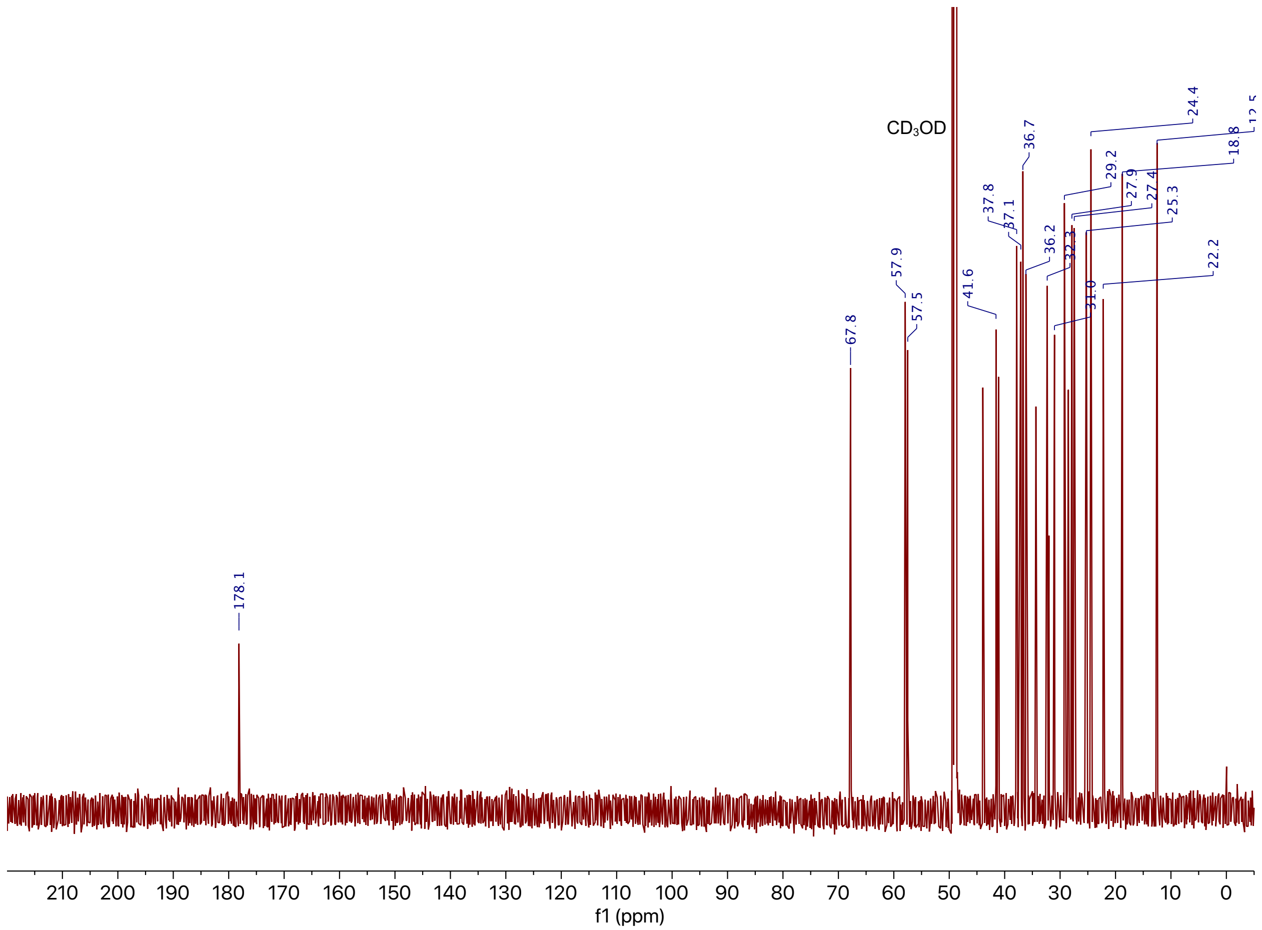


**Supplementary Figure 5. $^{13}C\{^1H\}$ NMR spectrum (151 MHz, $CD_3OD$) of isolithocholic acid.** The compound data and spectrum of isoLCA are representative of four synthesis experiments.

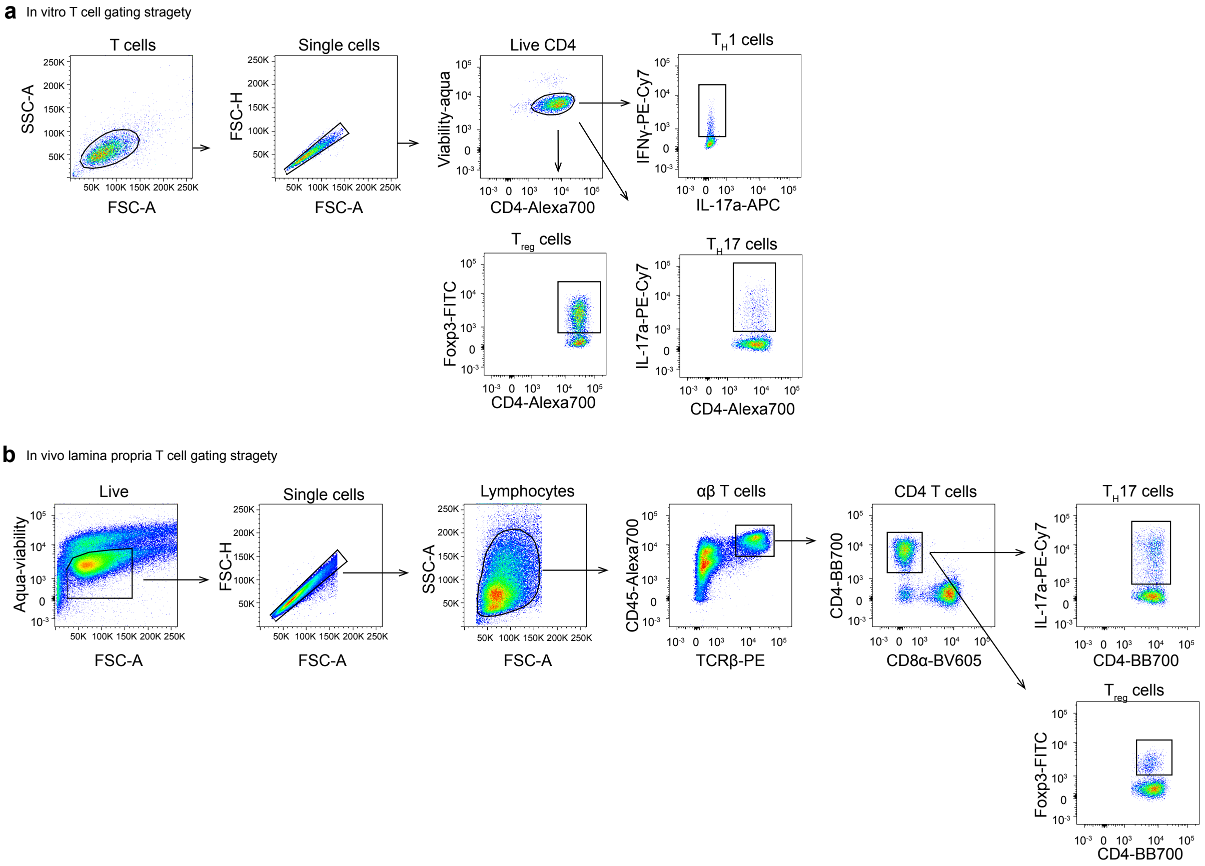


**Supplementary Figure 6.** Gating strategy for the flow cytometric analyses of in vitro cultured T cells (**a**) and in vivo derived cells from the lamina propria (**b**).

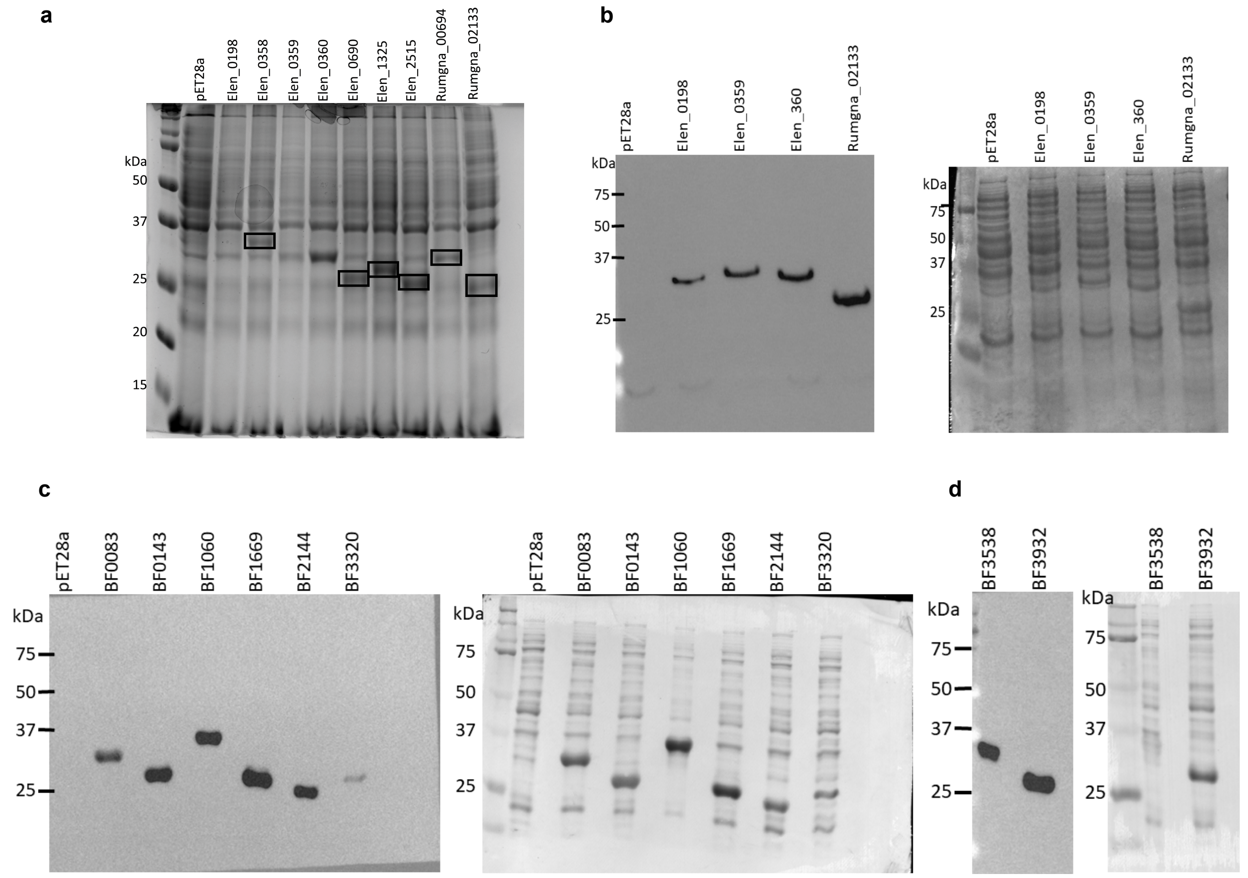


**Supplementary Figure 7. Gel analysis confirms heterologous expression of candidate genes. a**, SDS-PAGE analysis of candidate gene expression from *E. lenta* DSM 2243 and *R. gnavus* ATCC 29149 (Elen_0358, Elen_690, Elen_1325, Elen_2515, Rumgna_00694, and Rumgna_02133). **b**, Western blot of the expression of Elen_0198, Elen_0359, Elen_0360, and Rumgna_02133. Anti-His tag labeling (left). Amido black total protein stain of membrane (right). **c**, Western blot of the expression of BF0083, BF0143, BF1060, BF1669, BF2144, and BF3320. Anti-His tag labeling (left). Amido black total protein stain of membrane (right). **d**, Western blot of the expression of Bf3538 and Bf3932. Anti-His tag labeling (left). Amido black total protein stain of membrane (right).

a

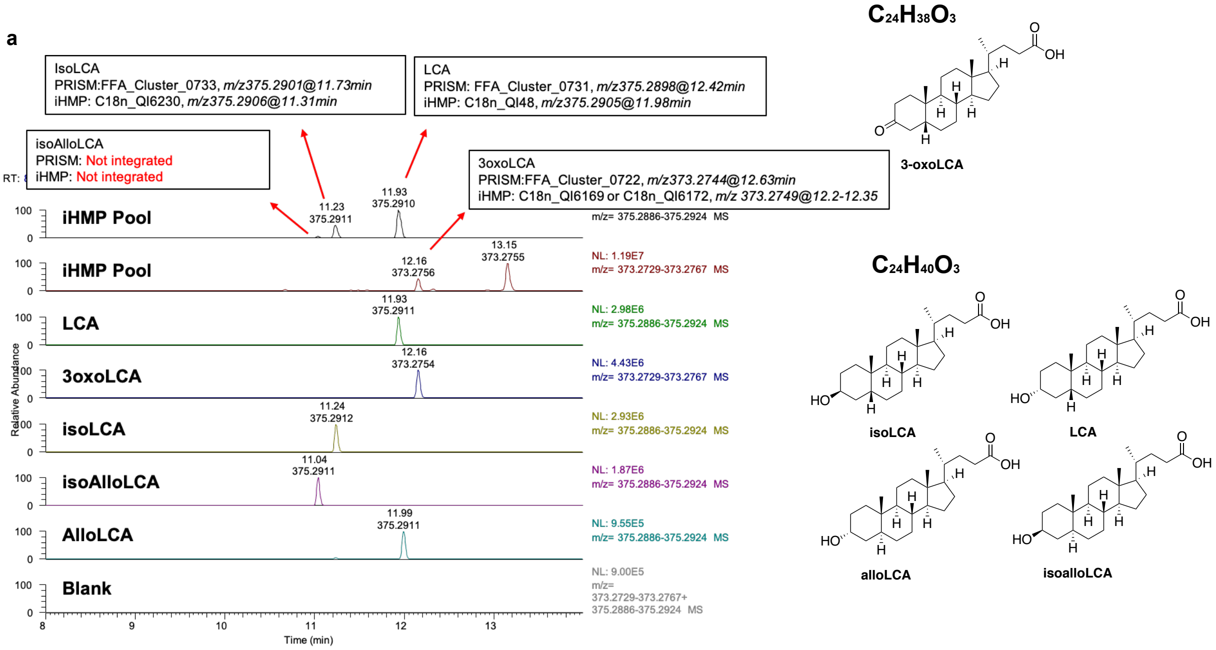


b

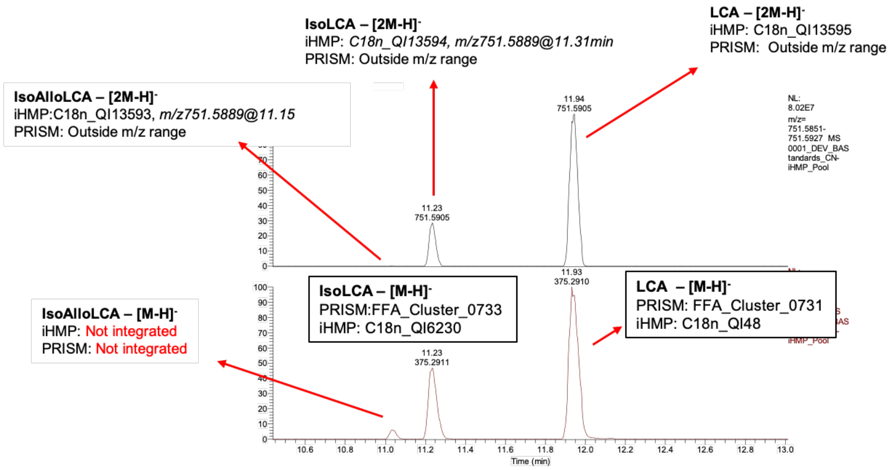


c

| Compound | m/z | RT | iHMP Feature | iHMP m/z | iHMP RT (min) | Adduct | PRISM Feature | PRISM m/z | PRISM RT (min) | Adduct |
|---|---|---|---|---|---|---|---|---|---|---|
| 3-oxoLCA | 373.2748 | 12.16 | C18n_QI6169 or C18n_QI6172 | 373.2749 | 12.2-12.35 | [M-H]- | FFA_Cluster_0722 | 373.274 | 12.63 | [M-H]- |
| LCA | 375.2905 | 11.93 | C18n_QI48 | 375.2905 | 11.98 | [M-H]- | FFA_Cluster_0731 | 375.29 | 12.42 | [M-H]- |
| | | | C18n_QI13595 | 751.5891 | 11.98 | [2M-H]- | | | | |
| isoLCA | 375.2905 | 11.24 | C18n_QI6230 | 375.2906 | 11.31 | [M-H]- | FFA_Cluster_0733 | 375.29 | 11.73 | [M-H]- |
| | | | C18n_QI13594 | 751.5889 | 11.31 | [2M-H]- | | | | |
| isoalloLCA | 375.2905 | 11.04 | C18n_QI13593 | 751.5889 | 11.15 | [2M-H]- | | | | |
| alloLCA | 375.2905 | 11.99 | Can't tell from LCA | | | | Can't tell from LCA | | | |

**d**

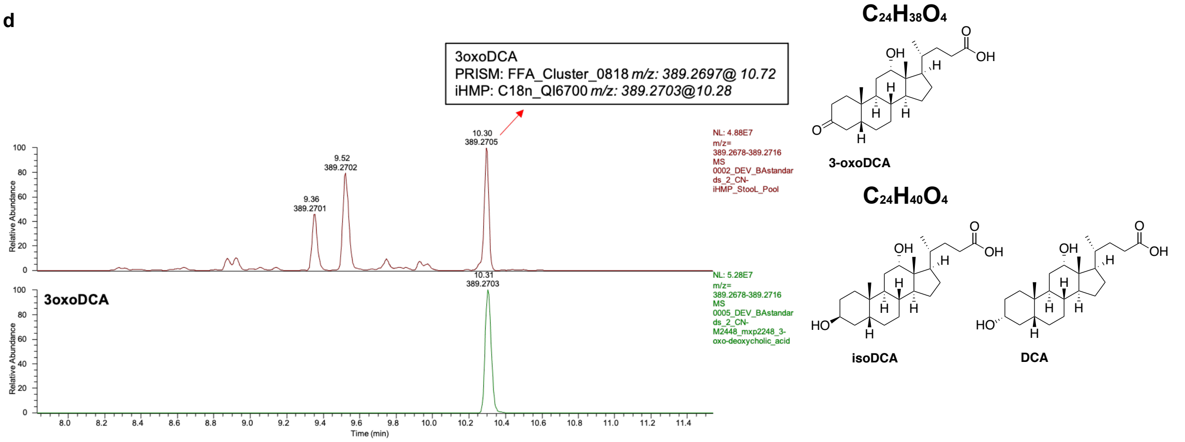
3oxoDCA
PRISM: FFA_Cluster_0818 m/z: 389.2697@ 10.72
iHMP: C18n_QI6700 m/z: 389.2703@10.28
NL: 4.88E7
m/z= 389.2678-389.2716
MS 0002_DEV_BAstandards_2_CN-iHMP_StooL_Pool
NL: 5.28E7
m/z= 389.2678-389.2716
MS 0005_DEV_BAstandards_2_CN-M2448_mxp2248_3-oxo-deoxycholic_acid
3oxoDCA
Relative Abundance
Time (min)
C24H38O4
3-oxoDCA
C24H40O4
isoDCA
DCA

**e**

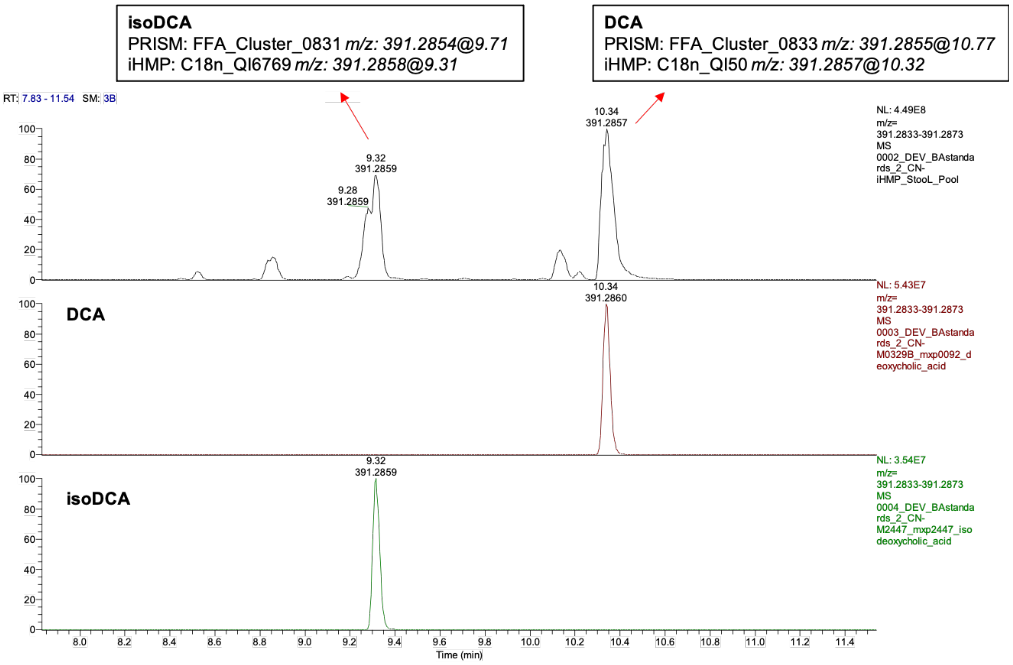
isoDCA
PRISM: FFA_Cluster_0831 m/z: 391.2854@9.71
iHMP: C18n_QI6769 m/z: 391.2858@9.31
DCA
PRISM: FFA_Cluster_0833 m/z: 391.2855@10.77
iHMP: C18n_QI50 m/z: 391.2857@10.32
RT: 7.83 - 11.54 SM: 3B
NL: 4.49E8
m/z= 391.2833-391.2873
MS 0002_DEV_BAstandards_2_CN-iHMP_StooL_Pool
NL: 5.43E7
m/z= 391.2833-391.2873
MS 0003_DEV_BAstandards_2_CN-M0329B_mxp0092_deoxycholic_acid
NL: 3.54E7
m/z= 391.2833-391.2873
MS 0004_DEV_BAstandards_2_CN-M2447_mxp2447_isodeoxycholic_acid
DCA
isoDCA
Time (min)

**Supplementary Figure 8. Identification of modified LCAs and DCAs mapped to PRISM and HMP2 metabolome database. a**, EIC of molecules of interest run alongside the HMP2 stool pool. **b**, Additional features (adducts and fragments) were detected for the LCA isomers. Among the strongest signals was one corresponding to the formation of a dimer (2M-H). Unlike the M-H, this dimer m/z was integrated for all compounds in the non-targeted data in HMP2. This feature is not reported in PRISM because the method used had a narrower m/z range. **c**, Summary table of modified LCAs in PRISM and HMP2. **d**, EIC of 3-oxoDCA run alongside the HMP2 stool pool. **e**, EIC of isoDCA and DCA run alongside the HMP2 stool pool.

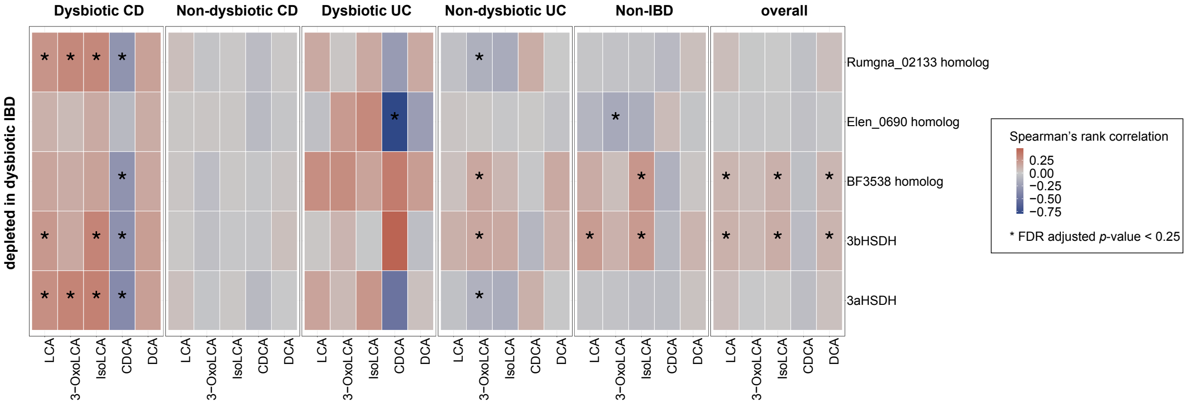


**Supplementary Figure 9. 3α- and 3β-HSDH homologs are likely to be positively correlated with 3-oxoLCA/isoLCA in HMP2.** Differentially abundant 3α-/3β-HSDH homologs (FDR adjusted p-value < 0.05) with at least one significant metabolite association (Spearman correlation with FDR adjusted p-value < 0.25). Correlations were computed over a subset of paired metabolomes and metagenomes from the HMP2 cohort derived from 106 participants (CD, n=50; UC, n=30; Non-IBD, n=26).

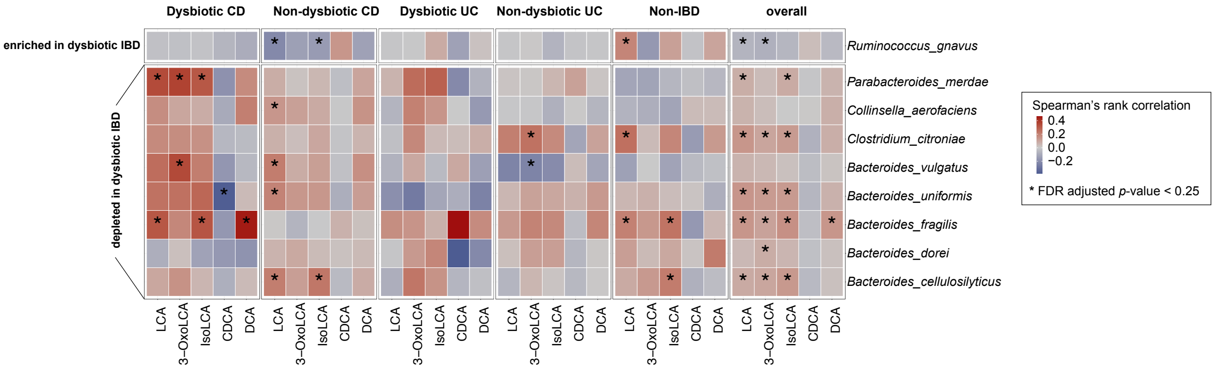


**Supplementary Figure 10. Species with 3α-/3β-HSDH activity are likely to be positively correlated with 3-oxoLCA/isoLCA in HMP2.** Differentially abundant species with validated 3α- /3β-HSDH activity (FDR adjusted p-value < 0.05) with at least one significant association (Spearman correlation FDR adjusted p-value < 0.25) with the five metabolites are shown for the paired metabolome and metagenome samples from 106 participants (CD, n=50; UC, n=30; Non-IBD, n=26) in HMP2.

**Supplementary Table 1 | Key Reagent Table**

| | | |
|---|---|---|
| **Chemicals** | | |
| DMSO | Sigma | D8418 |
| Ethyl acetate | Sigma | 319902 |
| HCl | Sigma | 258148 |
| Methanol | EMD Millipore | MX0475 |
| Ethanol | Sigma | E7023 |
| PBS | Genesee Scientific | 25-507 |
| Vitamin K1-hemin | BD Biosciences | 212354 |
| Trace minerals | ATCC | MD-TMS |
| Trace vitamins | ATCC | MD-VS |
| FBS | Genesee Scientific | 25-514 |
| Cellobiose | Sigma | C7252 |
| Maltose | Sigma | M5895 |
| Fructose | Sigma | F0127 |
| Arginine | Sigma | A5006 |
| Yeast extract | Bacto | 212750 |
| Malt extract | Sigma | 70146 |
| Dextrose | Sigma | G5767 |
| Phorbol 12-myristate 13-acetate (PMA) | Sigma | P1585 |
| Ionomycin | Sigma | I3909 |
| GolgiPlug | BD Biosciences | 555029 |
| Liberase TM | Sigma | 5401127001 |
| Dnase I (grade II, from bovine pancreas) | Sigma | 10104159001 |
| **Bile acids** | | |
| LCA | Sigma | L6250 |
| 3-oxoLCA | Steraloids | C1750-000 |
| IsoLCA | Steraloids | C1475-000 |
| IsoalloLCA | Steraloids | C0700-000 |
| AlloLCA | Steraloids | C0680-000 |
| T-LCA | Steraloids | C1472-000 |
| DCA | Sigma | D2510 |
| 3-oxoDCA | Steraloids | C1725-000 |
| IsoDCA | Steraloids | C1165-000 |
| CDCA | AstaTech | 76487 |
| TCA | Sigma | T4009 |
| βMCA | Steraloids | C1895-000 |
| GCA | Sigma | G2878 |
| **Antibodies** | | |
| *For flow cytometry; specific folocs are indicated in the figures.* | | |
| Anti-IL-17a (eBio17B7) | eBioscience | 25-7177-82 |
| Anti-FoxP3 (FJK-16s) | eBioscience | 11-5773-82 |
| Anti-RORγt (B2D) | eBioscience | 17-6981-82 |
| Anti-IFNγ (XMG1.2) | eBioscience | 48-7311-82 |
| Anti-CD3ε (145-2C11) | eBioscience | 48-0031-82 |

**Supplementary Table 1 | Key Reagent Table (continued)**

| | | |
|---|---|---|
| Anti-CD25 (PC61.5) | eBioscience | 25-0251-82 |
| Anti-CD62L (MEL-14) | eBioscience | 11-0621-85 |
| Anti-CD4 (RM4-5) | eBioscience | 56-0042-82 |
| Anti-CD45 (30-F11) | Biolegend | 103128 |
| Anti-CD8α (53-6.7) | Biolegend | 100744 |
| Anti-CD19 (6D5) | Biolegend | 115540 |
| Anti-CD44 (IM7) | Biolegend | 103032 |
| Anti-CD4 (RM4-5) | BD | 566407 |
| *For in vitro T cell culture* | | |
| Anti-CD3ε (145-2C11) | eBioscience | 16-0031-82 |
| Anti-CD28 (37.51) | eBioscience | 16-0281-82 |
| Anti-hamster IgG (whole molecule) goat affinity-purified | MPBio | 856984 |
| *For western blotting* | | |
| Anti-His-tag Antibody (rabbit polyclonal) | Cell Signaling | 2365 |
| **Cytokines** | | |
| Human IL-2 | PEPROTECH | 200-02 |
| Mouse IL-6 | eBioscience | 14-8061-62 |
| Mouse IL-12 | PEPROTECH | 210-12 |
| Human TGFβ1 | PEPROTECH | 100-21 |
| **Primers** | | |
| 16s_ElentaF | CAGCAGGGAAGAAATTCGAC | |
| 16s_ElentaR | TTGAGCCCTCGGATTAGAGA | |
| 16s_BragilisF | TGTAACACGTGTGTAGCCCC | |
| 16s_BragilisR | GGTTATGCTGAGGACTCTAG | |
| 16s_RgnavusF | GGTAGTTGGTGGGGTAACGG | |
| 16s_RgnavusR | TGTCTCAGTCCCAATGTGGC | |
| 16s_CinnocuumF | CAGCTCGTGTCGTGAGATGT | |
| 16s_CinnocuumR | CTCGCATGAGTCCCCAACTT | |

**Supplementary Table 2 | Human isolate screen metatables.** Table fields in the 1-10 sheets named PLATE1 through PLATE11 indicate eleven 96-well plates containing 990 colonies from two patients' stool samples were collected (Methods). Assays were performed in replicate plates using either 100 μM LCA or 100 μM 3-oxoLCA as the substrate. Conversion rates in percentage for each well are shown in yellow gradient highlights for 3α-HSDH activity and green gradient highlights for 3β-HSDH activity (named as PLATE X). Table fields in the 12-13 sheets follow the similar format as the previous eleven sheets for the positive bacterial metabolizers verified in bacterial culture tubes in triplicates and their conversion rates were shown in yellow gradient highlights for 3α-HSDH activity and green gradient highlights for 3β-HSDH activity.

**- PLATE1**

#### Sequence Summary Report

| | | LCA as substrate | | 3-OxoLCA as substrate | |
|---|---|---|---|---|---|
| Plate1 | 16S rDNA sequencing | 3-OxoLCA production (%) | isoLCA production (%) | LCA production (%) | isoLCA production (%) |
| P1-A1 | | 0.44604 | 0.00000 | 97.52560 | 0.04763 |
| P1-A10 | Bacteroides fragilis | 0.02749 | 0.12314 | 23.64419 | 31.58931 |
| P1-A11 | | | 0.06440 | 0.00000 | 0.00000 |
| P1-A12 | | 0.09501 | 0.00000 | 0.08520 | 0.00000 |
| P1-A2 | | 0.02380 | 0.09519 | 34.13654 | 5.18806 |
| P1-A3 | | 0.33216 | 0.00000 | 97.35310 | 0.03396 |
| P1-A4 | Catenibacterium mitsuokai | 0.03153 | 0.03378 | 0.02209 | 91.11984 |
| P1-A5 | | 0.14841 | 0.03544 | 98.09128 | 0.00000 |
| P1-A6 | | 0.25392 | 0.03681 | 97.84818 | 0.05627 |
| P1-A7 | | 0.00000 | 0.00000 | 0.12799 | 0.83481 |
| P1-A8 | | 0.21991 | 0.00000 | 83.22174 | 0.00000 |
| P1-A9 | | 0.12935 | 0.05605 | 35.99106 | 5.64579 |
| P1-B1 | | 0.02744 | 0.00000 | 97.30672 | 0.06369 |
| P1-B10 | | 0.43195 | 0.00000 | 5.09541 | 1.14501 |
| P1-B11 | | 0.02002 | 0.12977 | 0.00000 | 1.06125 |
| P1-B12 | Bacteroides dorei | 0.06070 | 0.00000 | 1.43275 | 9.54963 |
| P1-B2 | | 0.03586 | 0.05186 | 0.04441 | 21.90368 |
| P1-B3 | | 0.04314 | 0.02988 | 97.75235 | 0.07245 |
| P1-B4 | | 0.03052 | 0.00000 | 0.04674 | 0.02771 |
| P1-B5 | | 0.19071 | 0.00000 | 98.24994 | 0.08661 |
| P1-B6 | | 0.18698 | 0.04394 | 98.22308 | 0.13030 |
| P1-B7 | | 0.05210 | 0.10616 | 33.89225 | 4.82254 |
| P1-B8 | Bacteroides cellulosilyticus/ intestinalis | 0.02399 | 0.10406 | 0.00000 | 19.78247 |
| P1-B9 | | 0.00000 | 0.05178 | 33.02611 | 4.59280 |
| P1-C1 | | 0.11860 | 0.50134 | 94.94904 | 2.27888 |
| P1-C10 | | 0.09886 | 0.04715 | 33.83207 | 5.07079 |
| P1-C11 | Bacteroides vulgatus | 0.00000 | 0.08015 | 2.67418 | 7.53473 |
| P1-C12 | | 0.12093 | 0.00000 | 0.01887 | 0.11314 |
| P1-C2 | | 0.13438 | 0.07398 | 97.55771 | 0.00000 |
| P1-C3 | Bifidobacterium longum | 1.05545 | 0.00000 | 11.21356 | 16.90184 |
| P1-C4 | | 0.03670 | 0.04099 | 1.06936 | 0.00000 |

Table S2 - PLATE 1 (continued)

#### Sequence Summary Report

| | | | | | |
|---|---|---|---|---|---|
| P1-C5 | | 0.11097 | 0.05688 | 33.57453 | 5.04727 |
| P1-C6 | | 0.13993 | 0.05684 | 33.98396 | 5.02675 |
| P1-C7 | | 0.02790 | 0.00000 | 33.84925 | 4.72375 |
| P1-C8 | | 0.08036 | 0.04408 | 97.56328 | 0.26278 |
| P1-C9 | | 0.15869 | 0.02868 | 82.28676 | 0.00000 |
| P1-D1 | | 0.11188 | 0.08829 | 0.22668 | 2.20661 |
| P1-D10 | | 0.07922 | 0.02980 | 32.73685 | 4.91519 |
| P1-D11 | | 0.00000 | 0.00000 | 0.00000 | 31.10003 |
| P1-D12 | | 0.08476 | 0.20323 | 0.00858 | 0.00000 |
| P1-D2 | Bacteroides uniformis | 0.05365 | 0.03028 | 0.00000 | 28.13904 |
| P1-D3 | | 0.20849 | 0.00000 | 98.11412 | 0.00000 |
| P1-D4 | Bacteroides cellulosilyticus | 0.02121 | 0.07653 | 0.00000 | 27.95430 |
| P1-D5 | | 0.01667 | 0.03098 | 0.08488 | 0.57821 |
| P1-D6 | | 0.02126 | 0.06600 | 97.97734 | 0.00000 |
| P1-D7 | Bacteroides cellulosilyticus | 0.02476 | 0.07838 | 0.00000 | 28.68572 |
| P1-D8 | Parabacteroides merdae | 0.02301 | 0.05161 | 0.00000 | 17.86859 |
| P1-D9 | | 0.03792 | 0.03596 | 6.92291 | 1.38880 |
| P1-E1 | | 0.04253 | 0.03403 | 36.19458 | 5.75837 |
| P1-E10 | | 0.02118 | 0.00000 | 0.01217 | 0.03165 |
| P1-E11 | | 0.34342 | 0.00000 | 99.39099 | 0.10886 |
| P1-E12 | | 0.27301 | 0.13269 | 99.88469 | 0.08919 |
| P1-E2 | | 0.00000 | 0.11259 | 0.00000 | 0.24271 |
| P1-E3 | | 0.01720 | 0.02635 | 99.77632 | 0.00000 |
| P1-E4 | | 0.02336 | 0.00000 | 0.01630 | 0.07094 |
| P1-E5 | | 0.17141 | 0.03486 | 91.61052 | 1.53582 |
| P1-E6 | | 0.00000 | 0.06590 | 0.48258 | 8.59895 |
| P1-E7 | | 0.15425 | 0.10111 | 99.57845 | 0.08853 |
| P1-E8 | | 0.04396 | 0.03345 | 0.00000 | 14.44001 |
| P1-E9 | | 0.05514 | 0.05776 | 31.01376 | 1.19468 |
| P1-F1 | | 0.03585 | 0.04717 | 0.31920 | 0.00000 |
| P1-F10 | Bacteroides uniformis | 0.02609 | 0.09498 | 0.00000 | 21.57717 |
| P1-F11 | | 0.13779 | 0.00000 | 98.65641 | 0.09182 |
| P1-F12 | | 0.00000 | 0.16891 | 33.63692 | 3.40649 |
| P1-F2 | Bifidobacterium pseudocatenulatum | 0.08834 | 0.00000 | 1.73747 | 24.01944 |
| P1-F3 | Bacteroides cellulosilyticus/ intestinalis | 0.03397 | 0.03940 | 0.00000 | 27.15558 |
| P1-F4 | | 0.09185 | 0.02476 | 0.01002 | 0.09501 |
| P1-F5 | | 0.10864 | 0.02827 | 0.03360 | 0.33838 |

Table S2 - PLATE 1 (continued)

#### Sequence Summary Report

| | | | | | |
|---|---|---|---|---|---|
| P1-F6 | | 0.02059 | 0.00000 | 98.14397 | 0.05338 |
| P1-F7 | Bacteroides cellulosilyticus/ intestinalis | 0.06908 | 0.03878 | 0.00000 | 26.56359 |
| P1-F8 | | 0.02955 | 0.03666 | 0.00000 | 26.05974 |
| P1-F9 | | 0.05560 | 0.07295 | 99.06028 | 0.11376 |
| P1-G1 | | 0.11790 | 0.02846 | 98.06354 | 0.26742 |
| P1-G10 | | 0.21314 | 0.07202 | 99.33661 | 0.06390 |
| P1-G11 | | 0.19963 | 0.00000 | 99.73534 | 0.08072 |
| P1-G12 | | 0.00000 | 0.00000 | 1.69723 | 1.08668 |
| P1-G2 | Bacteroides rodentium | 0.00000 | 0.04228 | 0.00000 | 26.91233 |
| P1-G3 | | 0.06964 | 0.00000 | 0.04478 | 0.05463 |
| P1-G4 | | 0.09200 | 0.03686 | 99.37866 | 0.07556 |
| P1-G5 | | 0.04229 | 0.02481 | 0.00881 | 0.00000 |
| P1-G6 | | 0.04120 | 0.04161 | 0.02156 | 0.07405 |
| P1-G7 | | 0.02579 | 0.00000 | 0.55323 | 0.00000 |
| P1-G8 | | 0.13460 | 0.13430 | 0.13097 | 0.28562 |
| P1-G9 | | 0.13972 | 0.00000 | 0.01361 | 0.20632 |
| P1-H1 | Bacteroides uniformis | 0.04826 | 0.00000 | 0.04207 | 24.57614 |
| P1-H10 | | 0.18343 | 0.05449 | 98.50101 | 0.07989 |
| P1-H11 | | 0.04622 | 0.00000 | 0.03578 | 0.04149 |
| P1-H12 | Bacteroides sp. / Bacteroides cellulosilyticus | 0.09583 | 0.16651 | 0.00000 | 16.34538 |
| P1-H2 | Bifidobacterium longum | 1.18248 | 0.00000 | 7.56200 | 12.50201 |
| P1-H3 | | 0.27860 | 0.00000 | 97.97894 | 0.00000 |
| P1-H4 | | 0.02079 | 0.07200 | 0.02956 | 0.02814 |
| P1-H5 | Catenibacterium mitsuokai/ Catenibacterium sp. | 0.00000 | 0.12400 | 0.03176 | 85.95947 |
| P1-H6 | | 0.03371 | 0.04313 | 0.00920 | 0.02804 |
| P1-H7 | | 0.03182 | 0.00000 | 0.55225 | 1.71917 |
| P1-H8 | Bifidobacterium longum | 0.87125 | 0.00000 | 10.65127 | 16.71063 |
| P1-H9 | | 0.14837 | 0.08053 | 99.50336 | 0.00000 |

Table S2 - PLATE 2

### Sequence Summary Report

| | | LCA as substrate | | 3-OxoLCA as substrate | |
|---|---|---|---|---|---|
| Plate2 | 16S rDNA sequencing | 3-OxoLCA production (%) | isoLCA production (%) | LCA production (%) | isoLCA production (%) |
| P2-A1 | | 0.12362 | 0.25781 | 2.35201 | 0.85601 |
| P2-A10 | | 0.04265 | 0.15479 | 0.00000 | 0.00000 |
| P2-A11 | | 0.10892 | 0.55658 | 0.00000 | 0.09692 |
| P2-A12 | | 0.09747 | 0.15616 | 0.26136 | 39.61100 |
| P2-A2 | Lachnospira pectinoschiza | 0.04369 | 0.00000 | 0.03024 | 72.25347 |
| P2-A3 | | 0.00000 | 0.00000 | 0.78339 | 0.62250 |
| P2-A4 | | 0.10093 | 0.56119 | 3.95509 | 0.26514 |
| P2-A5 | | 0.11654 | 0.00000 | 0.00000 | 0.00000 |
| P2-A6 | | 0.24784 | 0.38302 | 0.00000 | 0.10160 |
| P2-A7 | | 0.11079 | 0.06529 | 0.10613 | 0.00000 |
| P2-A8 | | 0.22618 | 0.00000 | 0.00000 | 0.00000 |
| P2-A9 | | 0.07968 | 0.09809 | 0.04790 | 0.00000 |
| P2-B1 | | 0.17984 | 0.00000 | 1.98632 | 1.22241 |
| P2-B10 | | 0.09287 | 0.12224 | 0.00000 | 0.00000 |
| P2-B11 | | 0.08035 | 0.16093 | 0.00000 | 0.00000 |
| P2-B12 | | 0.00000 | 0.00000 | 0.46000 | 0.82530 |
| P2-B2 | | 0.07889 | 0.08219 | 0.45197 | 1.67710 |
| P2-B3 | | 0.23817 | 0.05960 | 0.00000 | 0.00000 |
| P2-B4 | | 0.15764 | 0.10205 | 0.00000 | 0.00000 |
| P2-B5 | | 0.15129 | 0.09621 | 0.00000 | 0.00000 |
| P2-B6 | Clostridium citroniae | 3.36551 | 5.59647 | 13.51704 | 1.48025 |
| P2-B7 | | 0.09889 | 0.07473 | 0.00000 | 6.70554 |
| P2-B8 | | 0.08234 | 0.07432 | 0.00000 | 0.00000 |
| P2-B9 | | 0.13331 | 0.11532 | 0.00000 | 0.00000 |
| P2-C1 | | 0.16696 | 0.09594 | 0.00000 | 0.00000 |
| P2-C10 | | 0.06820 | 0.08913 | 0.00000 | 0.41442 |
| P2-C11 | | 0.21407 | 0.07980 | 0.38246 | 0.00000 |
| P2-C12 | | 0.07224 | 0.13861 | 0.00000 | 0.00000 |
| P2-C2 | | 0.07709 | 0.00000 | 0.00000 | 0.00000 |
| P2-C3 | | 0.20313 | 0.10968 | 0.00000 | 0.00000 |
| P2-C4 | | 0.18868 | 0.00000 | 0.00000 | 0.00000 |

Table S2 - PLATE 2 (continued)

#### Sequence Summary Report

| | | | | | |
|---|---|---|---|---|---|
| P2-C5 | Clostridium citroniae | 0.99419 | 2.06441 | 2.86249 | 9.24056 |
| P2-C6 | | 0.21486 | 0.00000 | 0.00000 | 0.00000 |
| P2-C7 | | 0.34449 | 0.21059 | 0.64004 | 0.00000 |
| P2-C8 | | 0.11697 | 0.00000 | 0.00000 | 0.00000 |
| P2-C9 | | 0.11503 | 0.18023 | 0.00000 | 0.00000 |
| P2-D1 | | 0.05209 | 0.00000 | 0.28362 | 0.00000 |
| P2-D10 | | 0.19091 | 0.21042 | 0.61587 | 0.00000 |
| P2-D11 | | 0.09181 | 0.10310 | 0.00000 | 0.00000 |
| P2-D12 | Phocea massiliensis | 4.93739 | 0.08653 | 0.19928 | 0.00000 |
| P2-D2 | | 0.06144 | 0.10463 | 0.00000 | 0.00000 |
| P2-D3 | | 0.25152 | 0.00000 | 0.00000 | 0.00000 |
| P2-D4 | | 0.08885 | 0.07625 | 0.00000 | 0.00000 |
| P2-D5 | | 0.11977 | 0.10956 | 0.00000 | 0.00000 |
| P2-D6 | | 0.09073 | 0.06691 | 0.00000 | 0.00000 |
| P2-D7 | | 0.00000 | 0.05867 | 0.00000 | 19.10382 |
| P2-D8 | | 0.03079 | 0.00000 | 0.00000 | 0.00000 |
| P2-D9 | | 0.07003 | 0.06638 | 0.00000 | 0.00000 |
| P2-E1 | | 0.05233 | 0.00000 | 0.24556 | 0.23075 |
| P2-E10 | | 0.52698 | 0.00000 | 0.00000 | 0.00000 |
| P2-E11 | | 0.17586 | 0.07684 | 0.00000 | 0.00000 |
| P2-E12 | | 0.05184 | 0.20163 | 0.00000 | 0.00000 |
| P2-E2 | | 0.00000 | 0.09547 | 0.00000 | 0.00000 |
| P2-E3 | | 0.04098 | 0.00000 | 0.00000 | 0.00000 |
| P2-E4 | | 0.06008 | 0.06777 | 0.00000 | 0.00000 |
| P2-E5 | | 0.00000 | 0.11699 | 0.00000 | 0.00000 |
| P2-E6 | | 0.00000 | 0.13195 | 0.00000 | 0.00000 |
| P2-E7 | | 0.00000 | 0.11782 | 0.00000 | 0.00000 |
| P2-E8 | | 0.00000 | 0.25850 | 38.06156 | 16.32686 |
| P2-E9 | | 0.11684 | 0.00000 | 1.02610 | 0.00000 |
| P2-F1 | | 0.06205 | 0.07664 | 0.20147 | 0.00000 |
| P2-F10 | | 0.07216 | 0.00000 | 0.00000 | 3.60249 |
| P2-F11 | | 0.08346 | 0.25721 | 0.00000 | 0.00000 |
| P2-F12 | | 0.07238 | 0.15120 | 0.00000 | 0.00000 |
| P2-F2 | Lactobacillus rogosae | 0.00000 | 0.00000 | 0.00000 | 61.27788 |
| P2-F3 | | 0.05579 | 0.11597 | 0.00000 | 0.55976 |
| P2-F4 | | 0.17660 | 0.87449 | 0.55185 | 8.44035 |
| P2-F5 | | 0.23013 | 0.13340 | 0.50879 | 15.24047 |

Table S2 - PLATE 2 (continued)

#### Sequence Summary Report

| | | | | | |
|---|---|---|---|---|---|
| P2-F6 | | 0.09196 | 0.27258 | 0.56447 | 0.00000 |
| P2-F7 | | 0.18298 | 0.07514 | 0.00000 | 0.00000 |
| P2-F8 | | 0.06161 | 2.18083 | 0.00000 | 0.00000 |
| P2-F9 | | 0.08817 | 0.08383 | 0.00000 | 0.00000 |
| P2-G1 | Lactobacillus rogosae | 0.20420 | 0.14005 | 0.10165 | 54.33423 |
| P2-G10 | | 0.20916 | 0.32300 | 0.00000 | 0.00000 |
| P2-G11 | | 0.17788 | 0.00000 | 0.40326 | 0.00000 |
| P2-G12 | | 0.07224 | 0.12949 | 0.00000 | 0.00000 |
| P2-G2 | | 0.14303 | 0.00000 | 0.52918 | 0.26432 |
| P2-G3 | | 0.22225 | 0.06524 | 0.20311 | 11.54807 |
| P2-G4 | | 0.08906 | 0.08379 | 0.34502 | 6.42764 |
| P2-G5 | | 0.11325 | 0.23689 | 0.00000 | 0.00000 |
| P2-G6 | | 0.26286 | 0.10425 | 0.00000 | 0.00000 |
| P2-G7 | | 0.78107 | 0.27826 | 0.00000 | 0.87031 |
| P2-G8 | | 0.49365 | 0.00000 | 0.00000 | 0.00000 |
| P2-G9 | | 0.12059 | 0.00000 | 0.00000 | 0.00000 |
| P2-H1 | | 0.00000 | 0.09745 | 0.07264 | 9.19698 |
| P2-H10 | | 0.13659 | 0.26695 | 0.00000 | 0.00000 |
| P2-H11 | | 0.07915 | 0.00000 | 0.00000 | 0.00000 |
| P2-H12 | | 0.30870 | 0.00000 | 0.09035 | 0.00000 |
| P2-H2 | | 0.00000 | 0.24335 | 1.12360 | 0.00000 |
| P2-H3 | | 0.00000 | 0.00000 | 0.20900 | 0.31638 |
| P2-H4 | | 0.25369 | 0.08703 | 0.00000 | 0.00000 |
| P2-H5 | | 0.08617 | 0.10684 | 0.00000 | 0.00000 |
| P2-H6 | | 0.00000 | 0.18526 | 0.00000 | 0.00000 |
| P2-H7 | | 0.11217 | 0.00000 | 0.00000 | 0.00000 |
| P2-H8 | | 0.00000 | 0.07500 | 0.00000 | 0.00000 |
| P2-H9 | | 0.07229 | 0.12166 | 0.11989 | 0.00000 |

Table S2 - PLATE 3

### Sequence Summary Report

| | | LCA as substrate | | 3-OxoLCA as substrate | |
|---|---|---|---|---|---|
| Plate3 | 16S rDNA sequencing | 3-OxoLCA production (%) | isoLCA production (%) | LCA production (%) | isoLCA production (%) |
| P3-A1 | | 0.04669 | 0.04038 | 0.00000 | 0.00000 |
| P3-A10 | | 0.19176 | 0.28934 | 0.59043 | 0.00000 |
| P3-A11 | | 0.09648 | 0.10672 | 0.00000 | 0.00000 |
| P3-A12 | | 0.03841 | 0.06944 | 0.00000 | 0.00000 |
| P3-A2 | | 0.08071 | 0.06476 | 0.00000 | 0.00000 |
| P3-A3 | | 0.00000 | 0.05054 | 0.00000 | 0.00000 |
| P3-A4 | | 0.00000 | 0.04563 | 0.02744 | 0.00000 |
| P3-A5 | | 0.00000 | 0.00000 | 0.00000 | 0.00000 |
| P3-A6 | | 0.11399 | 0.05170 | 0.00000 | 0.00000 |
| P3-A7 | | 0.02592 | 0.00000 | 0.00000 | 0.00000 |
| P3-A8 | | 0.01965 | 0.02871 | 0.00000 | 0.00000 |
| P3-A9 | | 0.09683 | 0.00000 | 0.00000 | 0.00000 |
| P3-B1 | | 0.09850 | 0.07424 | 0.00000 | 0.00000 |
| P3-B10 | | 0.09136 | 0.00000 | 0.00000 | 0.00000 |
| P3-B11 | | 0.02729 | 0.03438 | 0.00000 | 0.00000 |
| P3-B12 | | 0.09084 | 0.06265 | 0.00000 | 0.00000 |
| P3-B2 | | 0.02591 | 0.13213 | 0.00000 | 0.00000 |
| P3-B3 | | 0.05501 | 0.00000 | 0.00000 | 0.00000 |
| P3-B4 | | 0.03076 | 0.03734 | 0.00000 | 0.00000 |
| P3-B5 | | 0.02277 | 0.00000 | 0.00000 | 0.00000 |
| P3-B6 | | 0.05139 | 0.00000 | 0.02944 | 0.00000 |
| P3-B7 | | 0.04378 | 0.05759 | 0.00000 | 0.00000 |
| P3-B8 | | 0.00000 | 0.10633 | 0.00000 | 0.00000 |
| P3-B9 | | 0.02763 | 0.00000 | 0.00000 | 0.00000 |
| P3-C1 | Clostridium perfringens | 5.57412 | 0.00000 | 24.02252 | 0.00000 |
| P3-C10 | | 0.71723 | 0.11029 | 14.57273 | 0.00000 |
| P3-C11 | Clostridium perfringens | 1.63981 | 0.07242 | 49.21909 | 0.00000 |
| P3-C12 | | 0.11244 | 0.16934 | 0.02252 | 0.00000 |
| P3-C2 | | 0.73451 | 0.00000 | 89.24200 | 0.00000 |
| P3-C3 | Clostridium perfringens | 1.07343 | 0.04509 | 99.95836 | 0.00000 |
| P3-C4 | | 0.53319 | 0.03920 | 75.70073 | 0.00000 |

Table S2 - PLATE 3 (continued)

#### Sequence Summary Report

| | | | | | |
|---|---|---|---|---|---|
| P3-C5 | | 0.41412 | 0.06213 | 100.00000 | 0.00000 |
| P3-C6 | | 0.77330 | 0.06612 | 95.72283 | 0.00000 |
| P3-C7 | | 0.24067 | 0.05611 | 97.00607 | 0.00000 |
| P3-C8 | | 0.13672 | 0.00000 | 0.03182 | 0.00000 |
| P3-C9 | | 0.05478 | 0.00000 | 73.88949 | 0.00000 |
| P3-D1 | | 0.96008 | 0.03758 | 0.78782 | 0.00000 |
| P3-D10 | | 0.32439 | 0.00000 | 72.36535 | 0.00000 |
| P3-D11 | | 0.48075 | 0.07814 | 98.89026 | 0.00000 |
| P3-D12 | Clostridium perfringens | 2.17388 | 0.14828 | 38.81224 | 0.00000 |
| P3-D2 | | 0.07424 | 0.07785 | 0.16122 | 0.00000 |
| P3-D3 | | 0.76857 | 0.10536 | 100.00000 | 0.00000 |
| P3-D4 | Clostridium perfringens | 1.15756 | 0.07215 | 99.25482 | 0.00000 |
| P3-D5 | Clostridium perfringens | 1.62943 | 0.08049 | 45.94084 | 0.00000 |
| P3-D6 | Clostridium perfringens | 1.04895 | 0.05123 | 87.27977 | 0.00000 |
| P3-D7 | | 0.07881 | 0.03989 | 29.77756 | 0.00000 |
| P3-D8 | | 0.95645 | 0.05630 | 49.13060 | 0.00000 |
| P3-D9 | | 0.69771 | 0.09575 | 45.30661 | 0.00000 |
| P3-E1 | Clostridium perfringens | 6.35653 | 0.06872 | 95.91740 | 0.00000 |
| P3-E10 | | 0.11336 | 0.15894 | 46.89499 | 0.00000 |
| P3-E11 | | 0.85345 | 0.07484 | 94.27890 | 0.00000 |
| P3-E12 | | 0.12876 | 0.11144 | 77.43442 | 0.00000 |
| P3-E2 | Clostridium perfringens | 5.96444 | 0.10204 | 1.50844 | 0.00000 |
| P3-E3 | | 0.23619 | 0.04408 | 90.97546 | 0.00000 |
| P3-E4 | | 0.66690 | 0.08194 | 1.20138 | 0.00000 |
| P3-E5 | | 0.68799 | 0.05956 | 20.97578 | 0.00000 |
| P3-E6 | | 0.79123 | 0.00000 | 47.18241 | 0.00000 |
| P3-E7 | | 0.61406 | 0.04279 | 16.54851 | 0.00000 |
| P3-E8 | | 0.20598 | 0.04937 | 99.63419 | 0.00000 |
| P3-E9 | Clostridium perfringens | 1.50024 | 0.04180 | 15.39342 | 0.00000 |
| P3-F1 | | 0.79554 | 0.04375 | 41.33744 | 0.00000 |
| P3-F10 | | 0.39814 | 0.00000 | 0.37659 | 0.00000 |
| P3-F11 | Clostridium perfringens | 1.94334 | 0.03782 | 74.48873 | 0.00000 |
| P3-F12 | | 0.12627 | 0.12733 | 51.94301 | 0.00000 |
| P3-F2 | | 0.94855 | 0.13978 | 0.03661 | 0.00000 |
| P3-F3 | | 0.29007 | 0.00000 | 82.98616 | 0.00000 |
| P3-F4 | | 0.10861 | 0.15433 | 4.76170 | 0.00000 |
| P3-F5 | | 0.23673 | 0.03057 | 98.11292 | 0.00000 |

Table S2 - PLATE 3 (continued)

#### Sequence Summary Report

| | | | | | |
|---|---|---|---|---|---|
| P3-F6 | Clostridium perfringens | 1.08508 | 0.71104 | 4.88340 | 0.00000 |
| P3-F7 | Clostridium perfringens | 0.01983 | 25.57211 | 2.60407 | 97.39593 |
| P3-F8 | Ruminococcus gnavus | 0.18143 | 27.10880 | 52.49258 | 46.14957 |
| P3-F9 | | 0.38468 | 0.00000 | 98.81418 | 0.00000 |
| P3-G1 | | 0.23529 | 0.03248 | 22.90683 | 0.00000 |
| P3-G10 | Clostridium perfringens | 1.14055 | 0.05183 | 97.08495 | 0.00000 |
| P3-G11 | Clostridium perfringens | 0.33132 | 27.73381 | 0.78034 | 98.35997 |
| P3-G12 | | 0.04911 | 0.06980 | 0.18124 | 0.29218 |
| P3-G2 | Ruminococcus gnavus | 0.24153 | 22.18868 | 48.42299 | 43.77997 |
| P3-G3 | | 0.58240 | 0.00000 | 32.75538 | 0.00000 |
| P3-G4 | | 0.61205 | 0.06516 | 36.90437 | 0.00000 |
| P3-G5 | | 0.53059 | 0.05577 | 43.83541 | 0.00000 |
| P3-G6 | | 0.04822 | 0.11764 | 98.49617 | 0.00000 |
| P3-G7 | | 0.03987 | 0.08650 | 100.00000 | 0.00000 |
| P3-G8 | | 0.74578 | 0.05977 | 67.48157 | 0.00000 |
| P3-G9 | Clostridium perfringens | 1.08321 | 0.07836 | 99.88428 | 0.00000 |
| P3-H1 | | 0.30252 | 0.08074 | 20.71473 | 0.00000 |
| P3-H10 | | 0.06888 | 0.06570 | 0.02478 | 0.00000 |
| P3-H11 | | 0.10529 | 0.00000 | 0.00000 | 0.00000 |
| P3-H12 | | 0.04466 | 0.06503 | 0.00000 | 0.00000 |
| P3-H2 | | 0.83888 | 0.04650 | 99.51622 | 0.00000 |
| P3-H3 | | 0.09169 | 0.00000 | 10.12035 | 0.00000 |
| P3-H4 | | 0.65165 | 0.03291 | 100.00000 | 0.00000 |
| P3-H5 | | 0.81587 | 0.46591 | 98.96462 | 0.00000 |
| P3-H6 | Clostridium perfringens | 1.81324 | 0.04175 | 25.68884 | 0.00000 |
| P3-H7 | | 0.08223 | 0.00000 | 13.78581 | 0.00000 |
| P3-H8 | | 0.87914 | 0.20223 | 99.77441 | 0.00000 |
| P3-H9 | | 0.05896 | 0.00000 | 0.00000 | 0.00000 |

Table S2 - PLATE 4

#### Sequence Summary Report

| | | LCA as substrate | | 3-OxoLCA as substrate | |
|---|---|---|---|---|---|
| Plate4 | 16S rDNA sequencing | 3-OxoLCA production (%) | isoLCA production (%) | LCA production (%) | isoLCA production (%) |
| P4-A1 | | 0.04131 | 0.06218 | 10.64598 | 2.68225 |
| P4-A10 | | 0.00000 | 0.00000 | 0.22471 | 0.11792 |
| P4-A11 | | 0.00000 | 0.06980 | 0.52897 | 0.00000 |
| P4-A12 | | 0.13686 | 0.00000 | 0.00000 | 0.00000 |
| P4-A2 | | 0.02931 | 0.00000 | 0.00000 | 0.00000 |
| P4-A3 | | 0.00000 | 0.08458 | 0.00000 | 0.00000 |
| P4-A4 | | 0.00000 | 0.09108 | 0.00000 | 0.07221 |
| P4-A5 | | 0.00000 | 0.05421 | 0.00000 | 0.00000 |
| P4-A6 | | 0.03343 | 0.00000 | 0.00000 | 0.00000 |
| P4-A7 | Bacillus coagulans | 0.00000 | 0.00000 | 0.17659 | 53.55268 |
| P4-A8 | Bacillus coagulans | 0.09968 | 0.00000 | 0.21065 | 64.15554 |
| P4-A9 | | 0.02208 | 0.00000 | 88.76916 | 5.42824 |
| P4-B1 | | 0.24958 | 0.00000 | 3.37946 | 16.66502 |
| P4-B10 | Phocea massiliensis | 1.32086 | 0.07467 | 86.42320 | 3.39077 |
| P4-B11 | | 0.00000 | 0.00000 | 0.00000 | 0.00000 |
| P4-B12 | | 0.09366 | 0.00000 | 0.00000 | 0.00000 |
| P4-B2 | Clostridium aldenense | 2.17636 | 0.76229 | 34.88247 | 61.22177 |
| P4-B3 | | 0.02809 | 0.05109 | 0.00000 | 0.08333 |
| P4-B4 | | 0.00000 | 0.10525 | 0.00000 | 0.00000 |
| P4-B5 | | 0.00000 | 0.00000 | 0.00000 | 0.00000 |
| P4-B6 | | 0.00000 | 0.00000 | 0.00000 | 0.00000 |
| P4-B7 | | 0.00000 | 0.09794 | 0.00000 | 0.00000 |
| P4-B8 | | 0.10434 | 0.08155 | 0.32707 | 0.00000 |
| P4-B9 | | 0.11091 | 0.00000 | 0.20501 | 0.00000 |
| P4-C1 | | 0.63381 | 0.00000 | 0.00000 | 0.00000 |
| P4-C10 | | 0.07761 | 0.00000 | 88.42194 | 4.94213 |
| P4-C11 | | 0.08862 | 0.00000 | 0.54059 | 0.53067 |
| P4-C12 | | 0.00000 | 0.00000 | 86.85391 | 0.21107 |
| P4-C2 | | 0.05000 | 0.00000 | 0.18937 | 5.52353 |
| P4-C3 | | 0.00000 | 0.07861 | 0.00000 | 0.00000 |
| P4-C4 | | 0.04253 | 0.00000 | 0.00000 | 0.00000 |

Table S2 - PLATE 4 (continued)

#### Sequence Summary Report

| | | | | | |
|---|---|---|---|---|---|
| P4-C5 | | 0.00000 | 0.08674 | 0.00000 | 0.00000 |
| P4-C6 | | 0.00000 | 0.07933 | 0.00000 | 0.00000 |
| P4-C7 | | 0.13754 | 0.00000 | 0.00000 | 0.00000 |
| P4-C8 | | 0.03823 | 0.00000 | 0.00000 | 0.00000 |
| P4-C9 | | 0.34134 | 0.08930 | 0.61111 | 0.56821 |
| P4-D1 | | 0.03713 | 0.05281 | 96.15253 | 3.51286 |
| P4-D10 | | 0.05479 | 0.00000 | 0.00000 | 0.00000 |
| P4-D11 | | 0.00000 | 0.00000 | 0.00000 | 0.00000 |
| P4-D12 | | 0.05575 | 0.03772 | 0.00000 | 0.00000 |
| P4-D2 | | 0.00000 | 0.09721 | 0.00000 | 0.00000 |
| P4-D3 | | 0.00000 | 0.10769 | 0.00000 | 0.00000 |
| P4-D4 | | 0.00000 | 0.17380 | 0.00000 | 0.00000 |
| P4-D5 | | 0.00000 | 0.00000 | 0.00000 | 0.00000 |
| P4-D6 | | 0.66174 | 0.10490 | 18.28490 | 4.51408 |
| P4-D7 | | 0.21615 | 0.00000 | 0.00000 | 0.00000 |
| P4-D8 | | 0.00000 | 0.06668 | 0.00000 | 0.00000 |
| P4-D9 | | 0.00000 | 0.18756 | 0.00000 | 0.00000 |
| P4-E1 | | 0.03683 | 0.00000 | 0.00000 | 0.00000 |
| P4-E10 | | 0.36658 | 0.00000 | 0.00000 | 0.00000 |
| P4-E11 | | 0.10058 | 0.08605 | 0.00000 | 0.00000 |
| P4-E12 | | 0.08287 | 0.25604 | 0.00000 | 0.00000 |
| P4-E2 | | 0.00000 | 0.06973 | 11.68166 | 13.29531 |
| P4-E3 | | 0.11122 | 0.00000 | 0.00000 | 0.00000 |
| P4-E4 | | 0.09589 | 0.11567 | 10.78270 | 1.10043 |
| P4-E5 | | 0.00000 | 0.07495 | 7.24833 | 1.54549 |
| P4-E6 | | 0.08421 | 0.08216 | 0.00000 | 0.00000 |
| P4-E7 | | 0.00000 | 0.00000 | 0.00000 | 0.00000 |
| P4-E8 | | 0.00000 | 0.07612 | 0.00000 | 0.00000 |
| P4-E9 | | 0.12054 | 0.00000 | 0.00000 | 0.00000 |
| P4-F1 | | 0.44095 | 0.57193 | 0.00000 | 0.00000 |
| P4-F10 | Clostridium perfringens | 21.62110 | 0.03675 | 93.58619 | 0.49505 |
| P4-F11 | Clostridium perfringens | 15.86673 | 0.07946 | 81.61440 | 0.32529 |
| P4-F12 | Clostridium perfringens | 6.26080 | 0.24302 | 82.76126 | 0.47362 |
| P4-F2 | | 0.00000 | 0.00000 | 0.00000 | 0.00000 |
| P4-F3 | | 0.00000 | 0.00000 | 0.03490 | 0.00000 |
| P4-F4 | | 0.00000 | 0.00000 | 0.00000 | 0.00000 |
| P4-F5 | | 0.00000 | 0.06903 | 0.00000 | 0.00000 |

Table S2 - PLATE 4 (continued)

#### Sequence Summary Report

| | | | | | |
|---|---|---|---|---|---|
| P4-F6 | Clostridium perfringens | 2.43789 | 0.07638 | 96.35296 | 0.00000 |
| P4-F7 | Clostridium perfringens | 1.31019 | 0.00000 | 92.41206 | 0.24208 |
| P4-F8 | Clostridium aldenense | 2.70961 | 0.44668 | 32.66861 | 6.71354 |
| P4-F9 | | 0.10195 | 0.04457 | 9.85060 | 0.18221 |
| P4-G1 | Clostridium perfringens | 3.06696 | 0.00000 | 83.72991 | 0.24167 |
| P4-G10 | Clostridium perfringens | 2.76120 | 0.00000 | 90.75036 | 0.15843 |
| P4-G11 | Clostridium perfringens | 16.70926 | 0.05442 | 88.09501 | 0.45870 |
| P4-G12 | Clostridium perfringens | 38.33151 | 0.19432 | 88.81575 | 0.46315 |
| P4-G2 | Ruminococcus gnavus | 0.68704 | 19.13871 | 73.00227 | 23.67676 |
| P4-G3 | Clostridium perfringens | 34.02939 | 0.04036 | 86.32963 | 1.08307 |
| P4-G4 | Clostridium perfringens | 29.73023 | 0.00000 | 90.96978 | 0.56736 |
| P4-G5 | Clostridium perfringens | 33.17728 | 0.00000 | 91.14764 | 0.40673 |
| P4-G6 | Clostridium perfringens | 26.80538 | 0.00000 | 89.99235 | 0.15519 |
| P4-G7 | Clostridium perfringens | 20.99210 | 0.00000 | 90.88960 | 0.37955 |
| P4-G8 | Clostridium perfringens | 12.21771 | 0.00000 | 97.80056 | 0.00000 |
| P4-G9 | Clostridium perfringens | 21.50584 | 0.02950 | 96.73967 | 0.34898 |
| P4-H1 | Clostridium perfringens | 34.20743 | 0.00000 | 84.64784 | 0.41899 |
| P4-H10 | Clostridium perfringens | 2.29594 | 0.00000 | 93.13971 | 0.60197 |
| P4-H11 | Clostridium perfringens | 3.34450 | 0.10198 | 87.11719 | 0.15687 |
| P4-H12 | Clostridium perfringens | 22.41180 | 0.00000 | 93.25305 | 0.34212 |
| P4-H2 | Clostridium perfringens | 1.68172 | 0.16722 | 89.76903 | 0.15412 |
| P4-H3 | Clostridium perfringens | 2.44717 | 0.00000 | 87.56843 | 0.16944 |
| P4-H4 | Clostridium perfringens | 1.72430 | 0.10102 | 88.99064 | 0.12301 |
| P4-H5 | Clostridium perfringens | 2.00302 | 0.04339 | 96.00979 | 0.26234 |
| P4-H6 | Clostridium perfringens | 24.30641 | 0.02946 | 88.94044 | 0.47597 |
| P4-H7 | Clostridium perfringens | 28.92956 | 0.04700 | 92.33367 | 6.61562 |
| P4-H8 | | 0.25859 | 0.00000 | 99.13505 | 0.00000 |
| P4-H9 | Clostridium perfringens | 1.64309 | 0.24424 | 88.99835 | 0.17231 |

**Table S2 - PLATE 5**

### Sequence Summary Report

| | | LCA as substrate | | 3-OxoLCA as substrate | |
|---|---|---|---|---|---|
| Plate5 | 16S rDNA sequencing | 3-OxoLCA production (%) | isoLCA production (%) | LCA production (%) | isoLCA production (%) |
| P5-A1 | Collinsella aerofaciens | 6.35832 | 0.17785 | 100.00000 | 0.00000 |
| P5-A10 | | 0.00000 | 0.00000 | 7.83984 | 0.48520 |
| P5-A11 | | 0.00000 | 0.13360 | 0.05129 | 0.00000 |
| P5-A12 | | 0.00000 | 0.24641 | 12.08809 | 40.28507 |
| P5-A2 | Bifidobacterium longum | 0.16383 | 1.34041 | 0.02586 | 1.99457 |
| P5-A3 | Collinsella aerofaciens | 4.48771 | 0.00000 | 99.43387 | 0.00000 |
| P5-A4 | | 0.00000 | 0.27769 | 0.00000 | 47.46135 |
| P5-A5 | Peptoniphilus harei | 0.00000 | 0.00000 | 1.91468 | 82.43851 |
| P5-A6 | | 0.00000 | 0.00000 | 0.00000 | 24.12213 |
| P5-A7 | | 0.10067 | 0.33789 | 0.31795 | 2.08422 |
| P5-A8 | | 0.50229 | 0.26525 | 0.60464 | 1.85514 |
| P5-A9 | Collinsella aerofaciens | 3.29954 | 0.46866 | 93.94722 | 0.00000 |
| P5-B1 | | 0.00000 | 0.00000 | 0.00000 | 0.00000 |
| P5-B10 | | 0.51554 | 0.61793 | 99.75056 | 0.13289 |
| P5-B11 | | 0.00000 | 0.51917 | 0.00000 | 0.00000 |
| P5-B12 | | 0.00000 | 0.00000 | 11.27135 | 0.71933 |
| P5-B2 | | 0.00000 | 0.73827 | 0.00000 | 0.00000 |
| P5-B3 | | 0.00000 | 0.93371 | 7.44576 | 0.55707 |
| P5-B4 | | 0.12778 | 0.86557 | 0.00000 | 0.00000 |
| P5-B5 | | 0.00000 | 0.00000 | 0.00000 | 0.00000 |
| P5-B6 | | 0.00000 | 0.37197 | 1.76918 | 93.87159 |
| P5-B7 | | 0.21324 | 0.00000 | 99.94257 | 0.00000 |
| P5-B8 | | 0.00000 | 0.00000 | 0.00000 | 0.06868 |
| P5-B9 | Bifidobacterium longum | 1.47636 | 2.29794 | 2.36788 | 7.56313 |
| P5-C1 | Collinsella aerofaciens | 4.16783 | 0.00000 | 47.03655 | 0.00000 |
| P5-C10 | | 0.00000 | 0.41384 | 0.00000 | 0.10701 |
| P5-C11 | Bifidobacterium longum | 0.75415 | 2.16399 | 0.09530 | 3.46336 |
| P5-C12 | | 0.00000 | 0.00000 | 0.00000 | 0.00000 |
| P5-C2 | Collinsella aerofaciens | 0.29273 | 6.13747 | 89.01105 | 8.91820 |
| P5-C3 | | 0.00000 | 0.00000 | 0.03648 | 10.35777 |
| P5-C4 | Bifidobacterium longum | 1.45408 | 2.12153 | 0.04816 | 4.52518 |

Table S2 - PLATE 5 (continued)

#### Sequence Summary Report

| | | | | | |
|---|---|---|---|---|---|
| P5-C5 | | 0.00000 | 0.00000 | 10.42374 | 0.62101 |
| P5-C6 | | 0.00000 | 0.15480 | 0.00000 | 0.00000 |
| P5-C7 | | 0.00000 | 0.00000 | 9.41029 | 0.72405 |
| P5-C8 | | 0.00000 | 0.00000 | 0.01172 | 0.00000 |
| P5-C9 | | 0.00000 | 0.67108 | 0.00000 | 0.00000 |
| P5-D1 | | 0.00000 | 0.28006 | 14.56738 | 1.56613 |
| P5-D10 | | 1.59360 | 12.22718 | 0.04800 | 0.00000 |
| P5-D11 | | 0.70233 | 0.00000 | 99.32288 | 0.00000 |
| P5-D12 | | 0.52481 | 0.18082 | 0.24355 | 4.30147 |
| P5-D2 | | 0.00000 | 0.00000 | 0.00000 | 2.28786 |
| P5-D3 | | 0.09161 | 0.28118 | 0.00000 | 0.28634 |
| P5-D4 | | 0.07542 | 0.28662 | 0.00000 | 0.00000 |
| P5-D5 | Collinsella aerofaciens | 0.82951 | 0.00000 | 99.39542 | 0.00000 |
| P5-D6 | | 0.00000 | 0.00000 | 0.00000 | 0.00000 |
| P5-D7 | | 0.00000 | 0.30827 | 0.05379 | 0.72024 |
| P5-D8 | | 0.00000 | 0.30035 | 0.06145 | 7.37611 |
| P5-D9 | | 0.00000 | 0.00000 | 5.78695 | 1.77300 |
| P5-E1 | Bacteroides dorei | 0.00000 | 0.00000 | 1.79280 | 0.98924 |
| P5-E10 | | 0.00000 | 0.00000 | 0.00000 | 0.10101 |
| P5-E11 | | 0.00000 | 0.00000 | 96.62218 | 0.83395 |
| P5-E12 | Bifidobacterium pseudocatenulatum | 0.00000 | 0.32475 | 12.02219 | 64.21384 |
| P5-E2 | | 1.61618 | 0.65487 | 0.13583 | 10.44787 |
| P5-E3 | Bifidobacterium longum | 1.45199 | 2.02448 | 2.74125 | 5.82997 |
| P5-E4 | | 0.00000 | 0.52835 | 0.22599 | 18.22831 |
| P5-E5 | | 0.08819 | 2.09533 | 3.56424 | 0.47473 |
| P5-E6 | Bifidobacterium longum | 1.27046 | 2.26067 | 25.94907 | 72.32475 |
| P5-E7 | | 0.09179 | 0.44059 | 0.05398 | 0.48957 |
| P5-E8 | | 0.64140 | 0.60553 | 99.96220 | 0.00000 |
| P5-E9 | | 0.00000 | 0.55511 | 11.92025 | 0.81696 |
| P5-F1 | | 0.10068 | 0.00000 | 0.00000 | 1.54274 |
| P5-F10 | | 0.00000 | 0.00000 | 0.00000 | 3.91072 |
| P5-F11 | | 0.00000 | 0.00000 | 0.02656 | 6.92200 |
| P5-F12 | Bifidobacterium longum | 0.09506 | 2.23312 | 1.80464 | 95.41979 |
| P5-F2 | | 1.22242 | 0.00000 | 100.00000 | 0.00000 |
| P5-F3 | Bifidobacterium longum | 0.20974 | 1.02181 | 0.03011 | 2.32614 |
| P5-F4 | | 0.00000 | 0.25372 | 0.00000 | 0.00000 |
| P5-F5 | | 0.00000 | 0.22998 | 1.14343 | 0.00000 |

Table S2 - PLATE 5 (continued)

#### Sequence Summary Report

| | | | | | |
|---|---|---|---|---|---|
| P5-F6 | Collinsella aerofaciens | 0.00000 | 0.00000 | 6.84434 | 70.72986 |
| P5-F7 | | 0.28076 | 0.56239 | 100.00000 | 0.00000 |
| P5-F8 | | 0.00000 | 0.00000 | 0.09846 | 0.40862 |
| P5-F9 | | 0.00000 | 0.34488 | 0.00000 | 0.00000 |
| P5-G1 | | 1.68478 | 0.00000 | 97.84053 | 0.00000 |
| P5-G10 | | 0.35874 | 0.00000 | 2.64588 | 0.00000 |
| P5-G11 | Bacteroides cellulosilyticus | 0.00000 | 0.00000 | 0.36476 | 95.67585 |
| P5-G12 | | 0.00000 | 0.00000 | 19.20505 | 0.00000 |
| P5-G2 | | 0.23720 | 0.00000 | 99.87347 | 0.00000 |
| P5-G3 | Bifidobacterium longum | 1.80020 | 2.33632 | 0.04824 | 5.57488 |
| P5-G4 | | 0.31259 | 0.00000 | 99.92285 | 0.05581 |
| P5-G5 | | 0.23247 | 0.81635 | 24.24741 | 0.00000 |
| P5-G6 | Bifidobacterium longum | 1.51745 | 0.77337 | 26.56201 | 71.91720 |
| P5-G7 | Bifidobacterium longum | 0.14271 | 2.01638 | 3.64790 | 5.26859 |
| P5-G8 | | 0.00000 | 0.20980 | 0.10540 | 86.94181 |
| P5-G9 | | 0.00000 | 0.00000 | 0.00000 | 0.72537 |
| P5-H1 | | 0.00000 | 0.00000 | 0.00000 | 19.76039 |
| P5-H10 | | 0.00000 | 0.21221 | 0.00000 | 0.00000 |
| P5-H11 | | 2.75009 | 0.00000 | 3.83345 | 0.00000 |
| P5-H12 | | 31.03629 | 0.00000 | 98.30083 | 0.00000 |
| P5-H2 | | 0.00000 | 0.12643 | 0.00000 | 99.39552 |
| P5-H3 | | 0.00000 | 0.64587 | 0.05376 | 0.00000 |
| P5-H4 | | 0.00000 | 0.27373 | 0.00000 | 0.00000 |
| P5-H5 | | 0.00000 | 0.29779 | 0.00000 | 0.10413 |
| P5-H6 | | 4.58706 | 0.67423 | 96.97683 | 0.00000 |
| P5-H7 | | 0.00000 | 0.27028 | 0.00000 | 0.00000 |
| P5-H8 | Bifidobacterium pseudocatenulatum | 0.00000 | 0.18864 | 0.00000 | 70.84122 |
| P5-H9 | Bacteroides fragilis | 0.00000 | 0.00000 | 0.36897 | 14.99881 |

**Table S2 - PLATE 6**

#### Sequence Summary Report

| Plate6 | 16S rDNA sequencing | LCA as substrate | | 3-OxoLCA as substrate | |
|---|---|---|---|---|---|
| | | 3-OxoLCA production (%) | isoLCA production (%) | LCA production (%) | isoLCA production (%) |
| P6-A1 | | 0.04739 | 0.00000 | 0.00000 | 0.00000 |
| P6-A10 | Collinsella intestinalis | 5.03699 | 0.05249 | 80.87789 | 0.20138 |
| P6-A11 | Collinsella intestinalis | 2.18681 | 0.05554 | 80.95377 | 0.23985 |
| P6-A12 | Collinsella intestinalis | 0.78213 | 0.00000 | 88.92832 | 0.16588 |
| P6-A2 | Collinsella intestinalis | 0.79731 | 0.03464 | 82.57390 | 0.15288 |
| P6-A3 | Peptoniphilus harei | 0.00000 | 0.03797 | 4.83845 | 83.78978 |
| P6-A4 | | 0.00000 | 0.03637 | 0.06501 | 0.00000 |
| P6-A5 | | 0.09483 | 0.05829 | 80.81056 | 0.00000 |
| P6-A6 | | 0.00000 | 0.09777 | 0.03097 | 0.00000 |
| P6-A7 | | 0.05779 | 0.09028 | 85.02009 | 0.14596 |
| P6-A8 | Collinsella intestinalis | 1.13690 | 0.00000 | 85.78117 | 0.00000 |
| P6-A9 | Peptoniphilus harei | 0.00000 | 0.00000 | 5.44749 | 77.61245 |
| P6-B1 | | 0.73471 | 0.61133 | 85.10960 | 0.08380 |
| P6-B10 | Collinsella intestinalis | 3.17529 | 0.03265 | 80.95873 | 0.00000 |
| P6-B11 | | 0.83753 | 0.07194 | 87.66914 | 0.22560 |
| P6-B12 | Collinsella intestinalis | 0.81978 | 0.00000 | 86.39735 | 0.31679 |
| P6-B2 | Collinsella intestinalis | 0.74456 | 0.00000 | 86.24369 | 0.28054 |
| P6-B3 | | 0.75311 | 0.11385 | 89.46062 | 0.26828 |
| P6-B4 | | 0.04970 | 0.04470 | 81.48347 | 0.34738 |
| P6-B5 | Collinsella intestinalis | 1.21965 | 0.00000 | 86.81133 | 0.00000 |
| P6-B6 | | 0.00000 | 0.00000 | 18.34299 | 20.28317 |
| P6-B7 | Collinsella aerofaciens | 0.02406 | 0.08259 | 78.76877 | 0.26058 |
| P6-B8 | Collinsella intestinalis | 1.09642 | 0.03280 | 90.20711 | 0.11975 |
| P6-B9 | Collinsella intestinalis | 1.50241 | 0.07474 | 85.28664 | 0.00000 |
| P6-C1 | | 0.06323 | 0.00000 | 90.75315 | 0.15363 |
| P6-C10 | Collinsella intestinalis | 1.26260 | 0.00000 | 84.35965 | 0.36637 |
| P6-C11 | Collinsella intestinalis | 0.08661 | 0.08183 | 84.84703 | 0.53000 |
| P6-C12 | | 0.00000 | 0.00000 | 0.59619 | 0.00000 |
| P6-C2 | | 0.96269 | 0.00000 | 88.12759 | 0.34382 |
| P6-C3 | Collinsella intestinalis | 1.76012 | 0.05587 | 83.43802 | 0.13792 |
| P6-C4 | | 0.07054 | 0.10732 | 90.44832 | 0.08360 |

Table S2 - PLATE 6 (continued)

#### Sequence Summary Report

| | | | | | |
|---|---|---|---|---|---|
| P6-C5 | Collinsella intestinalis | 1.09067 | 0.00000 | 90.76124 | 0.00000 |
| P6-C6 | Eggerthella lenta | 9.72812 | 19.75481 | 22.88320 | 6.79556 |
| P6-C7 | | 0.06349 | 0.08096 | 87.33350 | 0.00000 |
| P6-C8 | | 0.06119 | 0.04519 | 90.22181 | 0.08504 |
| P6-C9 | Collinsella intestinalis | 0.37149 | 0.00000 | 87.54129 | 0.00000 |
| P6-D1 | Bacteroides cellulosilyticus | 0.00000 | 0.10404 | 0.26346 | 29.34336 |
| P6-D10 | Bacteroides vulgatus | 0.00000 | 0.09198 | 5.27356 | 25.00190 |
| P6-D11 | | 0.00000 | 0.08321 | 0.00000 | 0.00000 |
| P6-D12 | | 0.00000 | 0.00000 | 17.08818 | 26.07555 |
| P6-D2 | Collinsella aerofaciens | 0.00000 | 0.07956 | 43.76892 | 9.46219 |
| P6-D3 | Collinsella aerofaciens | 0.42278 | 0.06742 | 88.72734 | 0.29577 |
| P6-D4 | | 0.00000 | 0.00000 | 16.75673 | 23.60516 |
| P6-D5 | | 0.00000 | 0.00000 | 0.10158 | 0.41138 |
| P6-D6 | | 0.00000 | 0.06727 | 0.11505 | 0.00000 |
| P6-D7 | | 0.00000 | 0.00000 | 0.09090 | 0.00000 |
| P6-D8 | | 0.00000 | 0.00000 | 0.10317 | 0.36630 |
| P6-D9 | | 0.00000 | 0.04473 | 43.08762 | 7.73917 |
| P6-E1 | | 0.48776 | 0.04171 | 85.87531 | 0.26055 |
| P6-E10 | | 0.00000 | 0.06279 | 0.08149 | 0.00000 |
| P6-E11 | | 0.00000 | 0.00000 | 17.93394 | 26.84629 |
| P6-E12 | | 0.00000 | 0.05864 | 0.11141 | 0.15901 |
| P6-E2 | | 0.00000 | 0.09794 | 0.12328 | 0.00000 |
| P6-E3 | | 0.02175 | 0.00000 | 87.36951 | 0.29076 |
| P6-E4 | Collinsella aerofaciens | 0.00000 | 0.00000 | 85.35505 | 0.46543 |
| P6-E5 | | 0.00000 | 0.05088 | 0.00000 | 0.13020 |
| P6-E6 | | 0.04599 | 0.00000 | 0.07757 | 0.00000 |
| P6-E7 | | 0.07318 | 0.00000 | 81.12988 | 0.14626 |
| P6-E8 | | 0.04474 | 0.00000 | 0.02863 | 18.39104 |
| P6-E9 | | 0.02917 | 0.00000 | 0.00000 | 0.07482 |
| P6-F1 | | 0.12430 | 0.04989 | 88.38759 | 0.32333 |
| P6-F10 | Bacteroides uniformis/ Bacteroides rodentium | 0.00000 | 0.00000 | 0.00000 | 35.35425 |
| P6-F11 | | 0.00000 | 0.00000 | 0.14505 | 0.19331 |
| P6-F12 | Bacteroides cellulosilyticus | 0.00000 | 0.00000 | 0.02763 | 32.41464 |
| P6-F2 | | 0.00000 | 0.00000 | 5.72530 | 10.11808 |
| P6-F3 | | 0.03621 | 0.00000 | 16.23888 | 25.21364 |
| P6-F4 | | 0.07799 | 0.05097 | 88.65441 | 0.34582 |
| P6-F5 | | 0.00000 | 0.00000 | 0.10286 | 0.20780 |

Table S2 - PLATE 6 (continued)

#### Sequence Summary Report

| | | | | | |
|---|---|---|---|---|---|
| P6-F6 | Collinsella aerofaciens | 0.23472 | 0.12534 | 88.77833 | 0.10917 |
| P6-F7 | | 0.06347 | 0.00000 | 0.12063 | 0.15586 |
| P6-F8 | | 0.00000 | 0.04827 | 45.05085 | 9.34177 |
| P6-F9 | | 0.00000 | 0.06564 | 0.04082 | 0.20986 |
| P6-G1 | Bacteroides fragilis | 0.00000 | 0.00000 | 1.35950 | 45.10501 |
| P6-G10 | | 0.00000 | 0.00000 | 0.04410 | 1.43551 |
| P6-G11 | Collinsella aerofaciens | 0.09276 | 0.00000 | 82.71868 | 0.27251 |
| P6-G12 | | 0.00000 | 0.00000 | 0.00000 | 0.14902 |
| P6-G2 | | 0.10549 | 0.04649 | 83.05247 | 0.24624 |
| P6-G3 | Bacteroides cellulosilyticus | 0.00000 | 0.10601 | 0.04566 | 46.84486 |
| P6-G4 | | 0.00000 | 0.00000 | 45.99707 | 8.15475 |
| P6-G5 | | 0.04633 | 0.00000 | 86.13539 | 0.36727 |
| P6-G6 | | 0.00000 | 0.00000 | 0.00000 | 0.00000 |
| P6-G7 | | 0.00000 | 0.08702 | 0.19198 | 1.95192 |
| P6-G8 | Bacteroides cellulosilyticus | 0.00000 | 0.28700 | 0.10881 | 31.92564 |
| P6-G9 | | 0.00000 | 0.04707 | 0.16848 | 0.14682 |
| P6-H1 | Bacteroides fragilis | 0.00000 | 0.00000 | 2.23577 | 47.41220 |
| P6-H10 | | 0.00000 | 0.07649 | 5.39632 | 9.60726 |
| P6-H11 | | 0.00000 | 0.00000 | 0.07380 | 0.00000 |
| P6-H12 | | 0.03280 | 0.06039 | 0.00000 | 0.00000 |
| P6-H2 | | 0.00000 | 0.00000 | 53.59665 | 9.13661 |
| P6-H3 | | 0.02946 | 0.00000 | 14.06397 | 22.14646 |
| P6-H4 | | 0.00000 | 0.06995 | 0.00000 | 0.00000 |
| P6-H5 | Bacteroides dorei | 0.00000 | 0.10640 | 8.96245 | 19.19089 |
| P6-H6 | | 0.27899 | 0.00000 | 0.15487 | 0.57411 |
| P6-H7 | | 0.05950 | 0.05554 | 11.40488 | 6.62472 |
| P6-H8 | | 0.03547 | 0.00000 | 0.00000 | 0.00000 |
| P6-H9 | | 0.00000 | 0.16372 | 0.54537 | 32.22356 |

Table S2 - PLATE 7

### Sequence Summary Report

| | | LCA as substrate | | 3-OxoLCA as substrate | |
|---|---|---|---|---|---|
| Plate7 | 16S rDNA sequencing | 3-OxoLCA production (%) | isoLCA production (%) | LCA production (%) | isoLCA production (%) |
| P7-A1 | | 0.00000 | 0.00000 | 0.00000 | 0.08149 |
| P7-A10 | Collinsella intestinalis | 2.12578 | 0.00000 | 83.82000 | 0.28239 |
| P7-A11 | Collinsella intestinalis | 2.08956 | 0.03026 | 86.53722 | 0.78321 |
| P7-A12 | Collinsella intestinalis | 2.47589 | 0.00000 | 87.65660 | 0.97189 |
| P7-A2 | Raoultibacter massiliensis | 10.90595 | 0.02497 | 44.33729 | 0.16447 |
| P7-A3 | Collinsella intestinalis | 2.15431 | 0.02110 | 83.68233 | 0.00000 |
| P7-A4 | Collinsella intestinalis | 1.33325 | 0.03000 | 86.22256 | 0.09864 |
| P7-A5 | | 0.01376 | 0.00000 | 0.09730 | 0.00000 |
| P7-A6 | | 0.40948 | 0.05383 | 88.34519 | 0.09547 |
| P7-A7 | | 0.00000 | 0.00000 | 0.05505 | 0.00000 |
| P7-A8 | Collinsella intestinalis | 1.51431 | 0.00000 | 89.24219 | 1.33112 |
| P7-A9 | Collinsella intestinalis | 2.77175 | 0.00000 | 89.11561 | 0.29084 |
| P7-B1 | Collinsella intestinalis | 1.27907 | 0.00000 | 86.15241 | 0.13486 |
| P7-B10 | Peptoniphilus harei | 0.00000 | 0.00000 | 6.05415 | 78.48273 |
| P7-B11 | Collinsella intestinalis | 1.93719 | 0.00000 | 86.48642 | 0.63739 |
| P7-B12 | Collinsella intestinalis | 1.33037 | 0.00000 | 89.83860 | 0.47198 |
| P7-B2 | Collinsella intestinalis | 1.88057 | 0.00000 | 89.96373 | 0.00000 |
| P7-B3 | | 0.73715 | 0.09642 | 85.48110 | 0.00000 |
| P7-B4 | Collinsella intestinalis | 1.34818 | 0.00000 | 90.60103 | 0.31750 |
| P7-B5 | Collinsella intestinalis | 1.13056 | 0.00000 | 90.81693 | 0.38595 |
| P7-B6 | Peptoniphilus harei | 0.00000 | 0.00000 | 4.57459 | 82.98978 |
| P7-B7 | | 0.87547 | 0.00000 | 91.63545 | 0.00000 |
| P7-B8 | Peptoniphilus harei | 0.00000 | 0.01662 | 5.33779 | 78.89307 |
| P7-B9 | Collinsella intestinalis | 3.04259 | 0.04234 | 93.14118 | 0.20256 |
| P7-C1 | Collinsella intestinalis | 1.87849 | 0.00000 | 87.22774 | 0.24323 |
| P7-C10 | Collinsella intestinalis | 2.02710 | 0.04405 | 90.02566 | 0.21949 |
| P7-C11 | Peptoniphilus harei | 0.01055 | 0.01592 | 4.60605 | 85.01845 |
| P7-C12 | Peptoniphilus harei | 0.00000 | 0.00000 | 5.45745 | 82.41179 |
| P7-C2 | | 0.81303 | 0.00000 | 92.60209 | 0.29581 |
| P7-C3 | Peptoniphilus harei | 0.00000 | 0.00000 | 6.02837 | 68.62382 |
| P7-C4 | | 0.69611 | 0.00000 | 93.53722 | 0.24009 |

Table S2 - PLATE 7 (continued)

#### Sequence Summary Report

| | | | | | |
|---|---|---|---|---|---|
| P7-C5 | | 0.91133 | 0.00000 | 94.41519 | 0.30115 |
| P7-C6 | Collinsella intestinalis | 1.22687 | 0.00000 | 93.63386 | 0.18295 |
| P7-C7 | | 0.06364 | 0.00000 | 93.61567 | 0.20943 |
| P7-C8 | | 0.03371 | 0.00000 | 92.56804 | 0.41874 |
| P7-C9 | | 0.26536 | 0.00000 | 93.64264 | 0.22941 |
| P7-D1 | Collinsella intestinalis | 2.41715 | 0.01726 | 90.66354 | 0.28759 |
| P7-D10 | | 0.04593 | 0.03971 | 92.22222 | 0.27757 |
| P7-D11 | Collinsella intestinalis | 3.30840 | 0.00000 | 91.51434 | 0.27907 |
| P7-D12 | Collinsella intestinalis | 1.38002 | 0.00000 | 92.70704 | 0.00000 |
| P7-D2 | Collinsella intestinalis | 3.40013 | 0.00000 | 95.32890 | 0.00000 |
| P7-D3 | Collinsella intestinalis | 2.66549 | 0.00000 | 94.46089 | 0.35258 |
| P7-D4 | Collinsella intestinalis | 1.19199 | 0.04695 | 92.95004 | 0.09506 |
| P7-D5 | Collinsella intestinalis | 1.87314 | 0.00000 | 95.63740 | 0.13336 |
| P7-D6 | | 0.05907 | 0.00000 | 94.85396 | 0.29450 |
| P7-D7 | | 0.21167 | 0.00000 | 93.29147 | 0.35016 |
| P7-D8 | | 0.61819 | 0.00000 | 93.07241 | 0.25222 |
| P7-D9 | Collinsella intestinalis | 1.49679 | 0.00000 | 95.15976 | 0.17585 |
| P7-E1 | Collinsella intestinalis | 4.98890 | 0.00000 | 88.38053 | 0.22665 |
| P7-E10 | | 0.27568 | 0.09186 | 92.58653 | 0.20042 |
| P7-E11 | Collinsella intestinalis | 1.63905 | 0.03831 | 92.41037 | 0.00000 |
| P7-E12 | Collinsella intestinalis | 1.21051 | 0.00000 | 89.24012 | 0.32901 |
| P7-E2 | Collinsella intestinalis | 3.23527 | 0.00000 | 94.40808 | 0.33750 |
| P7-E3 | Gordonibacter pamelaeae | 22.82634 | 3.64573 | 46.80347 | 16.49276 |
| P7-E4 | | 0.05923 | 0.00000 | 94.41703 | 0.36419 |
| P7-E5 | Collinsella intestinalis | 1.59325 | 0.04210 | 94.12225 | 0.27851 |
| P7-E6 | | 0.15709 | 0.00000 | 95.40575 | 0.17234 |
| P7-E7 | | 0.36160 | 0.00000 | 93.02594 | 0.00000 |
| P7-E8 | Collinsella intestinalis | 2.34159 | 0.00000 | 93.28704 | 0.37041 |
| P7-E9 | Collinsella intestinalis | 4.52619 | 0.00000 | 95.69402 | 0.00000 |
| P7-F1 | Collinsella intestinalis | 4.13674 | 0.00000 | 89.59086 | 0.18593 |
| P7-F10 | Eggerthella lenta | 6.84812 | 7.43788 | 17.86148 | 4.72162 |
| P7-F11 | Collinsella intestinalis | 2.83460 | 0.05118 | 90.98127 | 0.29330 |
| P7-F12 | | 0.49667 | 0.00000 | 90.12974 | 0.00000 |
| P7-F2 | | 0.00000 | 0.04336 | 55.87166 | 9.11060 |
| P7-F3 | Collinsella intestinalis | 2.12561 | 0.00000 | 95.80454 | 0.26831 |
| P7-F4 | | 0.93680 | 0.00000 | 94.96916 | 0.14224 |
| P7-F5 | Collinsella intestinalis | 1.06129 | 0.00000 | 93.71275 | 0.29961 |

Table S2 - PLATE 7 (continued)

#### Sequence Summary Report

| | | | | | |
|---|---|---|---|---|---|
| P7-F6 | | 0.48780 | 0.03590 | 97.99953 | 0.00000 |
| P7-F7 | | 0.14085 | 0.05765 | 92.90184 | 0.21848 |
| P7-F8 | Peptoniphilus harei | 0.00000 | 0.03670 | 5.40581 | 77.82719 |
| P7-F9 | | 0.07258 | 0.00000 | 92.60414 | 0.34866 |
| P7-G1 | Collinsella intestinalis | 8.86156 | 0.03543 | 91.46105 | 0.28099 |
| P7-G10 | | 0.19588 | 0.00000 | 88.43309 | 0.26984 |
| P7-G11 | | 0.05257 | 0.00000 | 87.56368 | 0.15353 |
| P7-G12 | | 0.19071 | 0.00000 | 29.96833 | 0.26837 |
| P7-G2 | Collinsella intestinalis | 3.93531 | 0.00000 | 93.39960 | 0.28829 |
| P7-G3 | Collinsella intestinalis | 3.42268 | 0.00000 | 92.52948 | 0.27991 |
| P7-G4 | | 0.00000 | 0.00000 | 0.00000 | 0.00000 |
| P7-G5 | | 0.00000 | 0.00000 | 0.00000 | 0.00000 |
| P7-G6 | | 0.99822 | 0.00000 | 91.11152 | 0.27057 |
| P7-G7 | Eggerthella lenta | 20.90626 | 3.87219 | 11.41579 | 3.47197 |
| P7-G8 | | 0.28227 | 0.00000 | 87.10975 | 0.23175 |
| P7-G9 | Eggerthella lenta | 36.55302 | 47.54130 | 25.20126 | 7.50220 |
| P7-H1 | | 0.00000 | 0.00000 | 0.04876 | 0.00000 |
| P7-H10 | | 0.04172 | 0.00000 | 27.36535 | 0.23802 |
| P7-H11 | | 0.36013 | 0.00000 | 80.45786 | 0.16247 |
| P7-H12 | Peptoniphilus harei | 0.00000 | 0.00000 | 0.02855 | 87.94745 |
| P7-H2 | Collinsella intestinalis | 2.68505 | 0.00000 | 91.02001 | 0.22526 |
| P7-H3 | Collinsella intestinalis | 2.83854 | 0.00000 | 86.13365 | 0.08121 |
| P7-H4 | Peptoniphilus harei | 0.00000 | 0.00000 | 4.88038 | 82.13127 |
| P7-H5 | | 0.18895 | 0.00000 | 0.02948 | 0.00000 |
| P7-H6 | Collinsella intestinalis | 1.12293 | 0.00000 | 82.14284 | 0.09609 |
| P7-H7 | Collinsella intestinalis | 0.12046 | 0.03505 | 85.77946 | 0.00000 |
| P7-H8 | | 0.79596 | 0.00000 | 87.11181 | 0.15417 |
| P7-H9 | | 0.06189 | 0.00000 | 87.97733 | 0.31982 |

Table S2 - PLATE 8

### Sequence Summary Report

| | | LCA as substrate | | 3-OxoLCA as substrate | |
|---|---|---|---|---|---|
| Plate8 | 16S rDNA sequencing | 3-OxoLCA production (%) | isoLCA production (%) | LCA production (%) | isoLCA production (%) |
| P8-A1 | | 0.02605 | 0.00000 | 0.11567 | 0.21029 |
| P8-A10 | | 0.24630 | 0.07986 | 17.49833 | 26.42472 |
| P8-A11 | | 0.11551 | 0.10361 | 8.99470 | 5.57623 |
| P8-A12 | | 0.04131 | 0.10457 | 79.74763 | 0.13274 |
| P8-A2 | | 0.09291 | 0.04534 | 16.59738 | 25.46388 |
| P8-A3 | | 0.03493 | 0.00000 | 49.90561 | 8.22788 |
| P8-A4 | | 0.00000 | 0.00000 | 84.23534 | 0.00000 |
| P8-A5 | | 0.15796 | 0.03469 | 48.69747 | 10.11218 |
| P8-A6 | | 0.05045 | 0.05990 | 83.43207 | 0.00000 |
| P8-A7 | | 0.04655 | 0.05439 | 77.04708 | 0.20146 |
| P8-A8 | | 0.02358 | 0.09240 | 81.20127 | 0.22826 |
| P8-A9 | | 0.09964 | 0.00000 | 0.06667 | 0.10516 |
| P8-B1 | Peptoniphilus harei | 0.15435 | 0.12996 | 5.38119 | 76.32120 |
| P8-B10 | | 0.04593 | 0.04818 | 0.68866 | 0.12481 |
| P8-B11 | Collinsella intestinalis | 5.66346 | 0.15010 | 85.24643 | 0.28681 |
| P8-B12 | Peptoniphilus harei | 0.02900 | 0.07365 | 4.17572 | 82.19406 |
| P8-B2 | Peptoniphilus harei | 0.00000 | 0.15806 | 5.64757 | 77.09931 |
| P8-B3 | | 0.03679 | 0.05010 | 0.00000 | 0.00000 |
| P8-B4 | | 0.07458 | 0.04139 | 0.42340 | 23.49654 |
| P8-B5 | | 0.28013 | 0.07043 | 33.09927 | 0.34213 |
| P8-B6 | Collinsella intestinalis | 13.70085 | 0.05964 | 87.37722 | 0.00000 |
| P8-B7 | Peptoniphilus harei | 0.03342 | 0.04146 | 5.61610 | 74.96311 |
| P8-B8 | | 0.06655 | 0.05562 | 85.90964 | 0.30073 |
| P8-B9 | Monoglobus pectinilyticus | 2.49782 | 0.00000 | 32.55076 | 0.24425 |
| P8-C1 | Monoglobus pectinilyticus | 4.17787 | 0.06604 | 36.30624 | 0.08437 |
| P8-C10 | Collinsella intestinalis | 12.69198 | 0.05114 | 86.84912 | 0.27043 |
| P8-C11 | Monoglobus pectinilyticus | 1.59023 | 0.00000 | 31.83882 | 0.20927 |
| P8-C12 | Collinsella intestinalis | 21.10642 | 0.06973 | 86.36487 | 0.25369 |
| P8-C2 | Collinsella intestinalis | 1.08661 | 0.00000 | 93.97437 | 0.22891 |
| P8-C3 | Collinsella intestinalis | 15.04968 | 0.13271 | 94.27427 | 0.20884 |
| P8-C4 | | 0.34233 | 0.17082 | 35.61409 | 0.32007 |

Table S2 - PLATE 8 (continued)

#### Sequence Summary Report

| | | | | | |
|---|---|---|---|---|---|
| P8-C5 | | 0.25077 | 0.13682 | 32.48034 | 0.00000 |
| P8-C6 | | 0.02147 | 0.17407 | 0.03886 | 0.00000 |
| P8-C7 | Collinsella intestinalis | 8.74024 | 0.02187 | 90.66014 | 0.25646 |
| P8-C8 | Monoglobus pectinilyticus | 2.39225 | 0.16230 | 90.01398 | 0.08637 |
| P8-C9 | Collinsella intestinalis | 1.45061 | 0.00000 | 90.83013 | 0.24617 |
| P8-D1 | Collinsella intestinalis | 23.14536 | 0.00000 | 93.26084 | 0.13663 |
| P8-D10 | Monoglobus pectinilyticus | 1.27344 | 0.03870 | 34.63793 | 0.29143 |
| P8-D11 | Peptoniphilus harei | 0.12227 | 0.12020 | 4.62060 | 78.05975 |
| P8-D12 | Collinsella intestinalis | 3.78555 | 0.00000 | 93.76057 | 0.00000 |
| P8-D2 | Eggerthella lenta | 48.81447 | 8.13633 | 47.23950 | 22.06168 |
| P8-D3 | Collinsella intestinalis | 12.84827 | 0.00000 | 93.06784 | 0.07164 |
| P8-D4 | Peptoniphilus harei | 0.07175 | 0.00000 | 5.20410 | 76.24708 |
| P8-D5 | Collinsella intestinalis | 3.89545 | 0.00000 | 94.34285 | 0.00000 |
| P8-D6 | | 0.78667 | 0.00000 | 92.75317 | 0.00000 |
| P8-D7 | | 0.31686 | 0.00000 | 32.00372 | 0.22009 |
| P8-D8 | Eggerthella lenta | 43.03595 | 9.90261 | 44.69670 | 17.64921 |
| P8-D9 | | 0.14188 | 0.00000 | 0.00000 | 0.00000 |
| P8-E1 | | 0.16003 | 0.09205 | 0.08349 | 0.00000 |
| P8-E10 | Collinsella intestinalis | 6.31235 | 0.20942 | 89.26881 | 0.00000 |
| P8-E11 | Monoglobus pectinilyticus | 1.27524 | 0.07853 | 36.88274 | 0.07824 |
| P8-E12 | Monoglobus pectinilyticus | 1.30826 | 0.31534 | 37.08392 | 0.40482 |
| P8-E2 | Peptoniphilus harei | 0.09433 | 0.00000 | 5.56548 | 73.71652 |
| P8-E3 | | 0.02717 | 0.00000 | 0.03038 | 0.00000 |
| P8-E4 | | 0.39670 | 0.21272 | 33.28813 | 0.23889 |
| P8-E5 | Collinsella intestinalis | 4.68123 | 0.03457 | 94.03777 | 0.21820 |
| P8-E6 | | 0.72109 | 0.10450 | 31.29797 | 0.17059 |
| P8-E7 | | 0.08094 | 0.16421 | 91.25153 | 0.00000 |
| P8-E8 | Peptoniphilus harei | 0.00000 | 0.04227 | 5.51501 | 73.74883 |
| P8-E9 | | 0.13039 | 0.04484 | 0.04554 | 0.00000 |
| P8-F1 | | 0.17217 | 0.07425 | 0.00000 | 0.00000 |
| P8-F10 | Peptoniphilus harei | 0.09541 | 0.11228 | 5.43071 | 76.19893 |
| P8-F11 | Christensenella timonensis/ | 3.30883 | 0.45147 | 0.02676 | 94.68729 |
| P8-F12 | Collinsella intestinalis | 6.89357 | 0.14064 | 0.01545 | 96.39206 |
| P8-F2 | Eggerthella lenta | 20.16405 | 11.92693 | 79.67543 | 17.84863 |
| P8-F3 | Eggerthella lenta | 50.26139 | 10.12897 | 42.85667 | 17.01935 |
| P8-F4 | Monoglobus pectinilyticus | 1.66689 | 0.05879 | 28.27532 | 0.00000 |
| P8-F5 | Peptoniphilus harei | 0.00000 | 0.08212 | 5.00071 | 75.71169 |

Table S2 - PLATE 8 (continued)

#### Sequence Summary Report

| | | | | | |
|---|---|---|---|---|---|
| P8-F6 | Peptoniphilus harei | 0.00000 | 0.04021 | 5.57630 | 73.55224 |
| P8-F7 | Collinsella intestinalis | 7.33921 | 0.03838 | 90.50494 | 0.00000 |
| P8-F8 | Collinsella intestinalis | 7.96172 | 0.03035 | 91.93853 | 0.00000 |
| P8-F9 | Collinsella intestinalis | 5.17015 | 0.05536 | 93.09270 | 0.00000 |
| P8-G1 | Eggerthella lenta | 45.25475 | 9.29648 | 35.89902 | 13.19238 |
| P8-G10 | Peptoniphilus harei | 0.02419 | 0.07393 | 4.70986 | 77.53346 |
| P8-G11 | | 0.08311 | 0.05493 | 85.09003 | 0.24411 |
| P8-G12 | | 0.98549 | 0.06522 | 42.51834 | 0.38487 |
| P8-G2 | | 0.04235 | 0.09199 | 88.44842 | 0.00000 |
| P8-G3 | Peptoniphilus harei | 0.05769 | 0.00000 | 16.81608 | 21.53425 |
| P8-G4 | | 0.04522 | 0.10061 | 86.12116 | 0.30557 |
| P8-G5 | Monoglobus pectinilyticus | 1.21695 | 0.00000 | 34.66803 | 0.21776 |
| P8-G6 | | 0.20583 | 0.27653 | 0.50734 | 26.58322 |
| P8-G7 | Collinsella intestinalis | 1.14283 | 0.06875 | 88.49695 | 0.19565 |
| P8-G8 | | 0.94443 | 0.03972 | 89.67492 | 0.00000 |
| P8-G9 | | 0.06353 | 0.08582 | 0.00000 | 0.00000 |
| P8-H1 | Monoglobus pectinilyticus | 3.10662 | 0.09463 | 42.75429 | 0.39168 |
| P8-H10 | Monoglobus pectinilyticus | 1.16742 | 0.14987 | 30.50877 | 0.23948 |
| P8-H11 | | 0.59432 | 0.00000 | 30.63150 | 0.17592 |
| P8-H12 | Peptoniphilus harei | 0.06031 | 0.05304 | 4.59360 | 80.87113 |
| P8-H2 | Monoglobus pectinilyticus | 2.08661 | 0.08425 | 44.31678 | 0.31906 |
| P8-H3 | Monoglobus pectinilyticus | 1.49068 | 0.04943 | 36.89020 | 0.37450 |
| P8-H4 | Monoglobus pectinilyticus | 1.41018 | 0.03699 | 7.72229 | 2.56529 |
| P8-H5 | Eggerthella lenta | 37.00561 | 14.29137 | 3.33597 | 1.08090 |
| P8-H6 | Monoglobus pectinilyticus | 1.41475 | 0.12192 | 31.71054 | 0.08246 |
| P8-H7 | | 0.07230 | 0.00000 | 0.00000 | 0.00000 |
| P8-H8 | | 0.78278 | 0.26257 | 30.97117 | 0.06489 |
| P8-H9 | | 0.81762 | 0.00000 | 34.11502 | 0.22899 |

**Table S2 - PLATE 9**

### Sequence Summary Report

| | | LCA as substrate | | 3-OxoLCA as substrate | |
|---|---|---|---|---|---|
| Plate9 | 16S rDNA sequencing | 3-OxoLCA production (%) | isoLCA production (%) | LCA production (%) | isoLCA production (%) |
| P9-A1 | Collinsella intestinalis | 7.28082 | 0.00000 | 99.07172 | 0.00000 |
| P9-A10 | Collinsella intestinalis | 4.40681 | 0.00000 | 98.39648 | 0.00000 |
| P9-A11 | Collinsella intestinalis | 4.18293 | 0.00000 | 90.20278 | 0.00000 |
| P9-A12 | Collinsella intestinalis | 5.77091 | 0.00000 | 98.97189 | 0.00000 |
| P9-A2 | Collinsella intestinalis | 3.96405 | 0.00000 | 99.39062 | 0.51599 |
| P9-A3 | | 0.00000 | 0.05769 | 0.00000 | 0.00000 |
| P9-A4 | | 0.07546 | 0.00000 | 0.00000 | 0.00000 |
| P9-A5 | Collinsella intestinalis | 2.56883 | 0.07237 | 99.74925 | 0.00000 |
| P9-A6 | Collinsella intestinalis | 2.70121 | 0.06270 | 99.14807 | 0.00000 |
| P9-A7 | | 0.80434 | 0.00000 | 99.30726 | 0.00000 |
| P9-A8 | Collinsella intestinalis | 2.28450 | 0.00000 | 98.01761 | 0.29373 |
| P9-A9 | Collinsella intestinalis | 3.19246 | 0.09187 | 99.20526 | 0.00000 |
| P9-B1 | Collinsella intestinalis | 4.39110 | 0.09839 | 96.18479 | 0.20904 |
| P9-B10 | | 0.00000 | 0.00000 | 15.12151 | 2.67324 |
| P9-B11 | Bifidobacterium longum | 0.05328 | 0.07403 | 0.00000 | 97.24012 |
| P9-B12 | | 0.00000 | 0.06169 | 0.00000 | 0.12563 |
| P9-B2 | Collinsella intestinalis | 6.02249 | 0.13259 | 98.65584 | 0.17824 |
| P9-B3 | | 0.00000 | 0.00000 | 2.62904 | 40.49974 |
| P9-B4 | | 0.03795 | 0.00000 | 2.40203 | 5.02133 |
| P9-B5 | | 0.00000 | 0.05663 | 0.00000 | 3.47494 |
| P9-B6 | | 0.08775 | 0.00000 | 99.72768 | 0.00000 |
| P9-B7 | | 0.00000 | 0.00000 | 17.07097 | 2.63041 |
| P9-B8 | | 0.03520 | 0.00000 | 0.00000 | 0.00000 |
| P9-B9 | | 0.04006 | 0.00000 | 0.00000 | 0.00000 |
| P9-C1 | | 0.00000 | 0.00000 | 0.00000 | 0.00000 |
| P9-C10 | | 0.07964 | 0.00000 | 1.88794 | 3.55003 |
| P9-C11 | | 0.09238 | 0.00000 | 24.48863 | 43.94762 |
| P9-C12 | | 0.69941 | 0.00000 | 97.11219 | 0.90159 |
| P9-C2 | Parolsenella catena | 0.00000 | 0.07356 | 0.81275 | 95.04366 |
| P9-C3 | | 0.00000 | 0.00000 | 0.00000 | 0.00000 |
| P9-C4 | | 0.00000 | 0.00000 | 0.00000 | 3.06098 |

Table S2 - PLATE 9 (continued)

#### Sequence Summary Report

| | | | | | |
|---|---|---|---|---|---|
| P9-C5 | | 0.04963 | 0.09247 | 0.03656 | 0.00000 |
| P9-C6 | | 0.04087 | 0.05772 | 0.91301 | 1.61369 |
| P9-C7 | | 0.00000 | 0.00000 | 0.00000 | 4.85521 |
| P9-C8 | Parolsenella catena | 0.00000 | 0.00000 | 0.11359 | 96.75234 |
| P9-C9 | | 0.08782 | 0.72376 | 2.67119 | 41.39989 |
| P9-D1 | | 0.00000 | 0.09846 | 0.32721 | 0.12020 |
| P9-D10 | | 0.00000 | 0.00000 | 0.00000 | 0.00000 |
| P9-D11 | | 0.05300 | 0.00000 | 0.00000 | 0.00000 |
| P9-D12 | | 0.63862 | 0.12399 | 98.69657 | 0.13602 |
| P9-D2 | | 0.00000 | 0.00000 | 14.88628 | 2.49990 |
| P9-D3 | | 0.00000 | 0.08075 | 0.00000 | 0.00000 |
| P9-D4 | | 0.05718 | 0.00000 | 98.91456 | 0.00000 |
| P9-D5 | Parolsenella catena | 0.03272 | 0.08326 | 0.43914 | 97.18028 |
| P9-D6 | | 0.00000 | 0.00000 | 0.03597 | 0.00000 |
| P9-D7 | | 0.00000 | 0.09548 | 0.08559 | 11.99631 |
| P9-D8 | | 0.13098 | 0.00000 | 99.11986 | 0.11436 |
| P9-D9 | Parabacteroides chongii | 0.00000 | 0.00000 | 0.08285 | 16.12584 |
| P9-E1 | | 0.00000 | 0.13739 | 0.00000 | 0.00000 |
| P9-E10 | | 0.00000 | 0.05884 | 0.00000 | 0.07089 |
| P9-E11 | | 0.00000 | 0.00000 | 1.14001 | 3.40083 |
| P9-E12 | | 0.03352 | 0.06095 | 0.00000 | 0.00000 |
| P9-E2 | | 0.00000 | 0.06122 | 0.00000 | 0.00000 |
| P9-E3 | | 0.59485 | 0.11064 | 95.54359 | 0.00000 |
| P9-E4 | | 0.00000 | 0.00000 | 0.02774 | 0.00000 |
| P9-E5 | | 0.00000 | 0.00000 | 0.00000 | 0.00000 |
| P9-E6 | | 0.00000 | 0.05504 | 23.01025 | 4.02213 |
| P9-E7 | | 0.00000 | 0.00000 | 0.02937 | 0.00000 |
| P9-E8 | | 0.00000 | 0.00000 | 0.03200 | 0.10620 |
| P9-E9 | | 0.00000 | 0.00000 | 0.00000 | 7.37578 |
| P9-F1 | | 0.04859 | 0.00000 | 2.24862 | 4.15510 |
| P9-F10 | Collinsella intestinalis | 10.64282 | 0.70182 | 94.95824 | 0.00000 |
| P9-F11 | | 0.00000 | 0.00000 | 3.51141 | 40.56271 |
| P9-F12 | Collinsella intestinalis | 4.96933 | 0.00000 | 94.11615 | 0.47160 |
| P9-F2 | Collinsella intestinalis | 3.68657 | 0.06006 | 97.69122 | 0.00000 |
| P9-F3 | Collinsella intestinalis | 4.91153 | 0.11714 | 96.51039 | 0.00000 |
| P9-F4 | Collinsella intestinalis | 1.76157 | 0.00000 | 99.56819 | 0.15655 |
| P9-F5 | Collinsella intestinalis | 2.78870 | 0.04439 | 96.61440 | 0.30722 |

Table S2 - PLATE 9 (continued)

#### Sequence Summary Report

| | | | | | |
|---|---|---|---|---|---|
| P9-F6 | Collinsella intestinalis | 2.67907 | 0.00000 | 95.50631 | 0.27353 |
| P9-F7 | | 0.00000 | 0.00000 | 4.24911 | 48.63308 |
| P9-F8 | Collinsella intestinalis | 2.17470 | 0.00000 | 97.26683 | 0.12911 |
| P9-F9 | Collinsella intestinalis | 3.59277 | 0.00000 | 96.47777 | 0.12295 |
| P9-G1 | Collinsella intestinalis | 6.97920 | 0.00000 | 95.04295 | 0.00000 |
| P9-G10 | | 0.05127 | 0.00000 | 4.18809 | 40.77875 |
| P9-G11 | Collinsella intestinalis | 2.11541 | 0.00000 | 96.49217 | 0.07038 |
| P9-G12 | Collinsella intestinalis | 4.45546 | 0.00000 | 95.76433 | 0.30002 |
| P9-G2 | Collinsella intestinalis | 5.62839 | 0.00000 | 94.49581 | 0.00000 |
| P9-G3 | Collinsella intestinalis | 4.93217 | 0.05593 | 96.59442 | 0.10322 |
| P9-G4 | Collinsella intestinalis | 1.59201 | 0.00000 | 97.11176 | 0.15258 |
| P9-G5 | | 0.60457 | 0.00000 | 97.72769 | 0.13630 |
| P9-G6 | Collinsella intestinalis | 1.08709 | 0.00000 | 97.34953 | 0.16526 |
| P9-G7 | | 0.72354 | 0.00000 | 95.38806 | 0.28939 |
| P9-G8 | | 0.63494 | 0.00000 | 92.89789 | 0.10749 |
| P9-G9 | Collinsella intestinalis | 3.89871 | 0.00000 | 96.45528 | 0.14326 |
| P9-H1 | | 0.00000 | 0.00000 | 3.25783 | 0.00000 |
| P9-H10 | Collinsella intestinalis | 4.23827 | 0.00000 | 99.60749 | 0.12120 |
| P9-H11 | Collinsella intestinalis | 2.62418 | 0.00000 | 96.81246 | 0.08107 |
| P9-H12 | Collinsella intestinalis | 12.69548 | 0.00000 | 95.72582 | 0.09783 |
| P9-H2 | Collinsella intestinalis | 2.85363 | 0.00000 | 93.15008 | 0.29110 |
| P9-H3 | Collinsella intestinalis | 2.86053 | 0.05673 | 96.41722 | 0.32567 |
| P9-H4 | Collinsella intestinalis | 1.24102 | 0.00000 | 96.04583 | 0.12649 |
| P9-H5 | Peptoniphilus harei | 0.00000 | 0.00000 | 3.42344 | 50.89230 |
| P9-H6 | | #DIV/0! | #DIV/0! | 97.23289 | 0.21211 |
| P9-H7 | Collinsella intestinalis | 2.35148 | 0.00000 | 97.70918 | 0.09586 |
| P9-H8 | | 0.80755 | 0.00000 | 94.58732 | 0.28606 |
| P9-H9 | Collinsella intestinalis | 2.80643 | 0.00000 | 97.22985 | 0.12701 |

Table S2 - PLATE 10

### Sequence Summary Report

| | | LCA as substrate | | 3-OxoLCA as substrate | |
|---|---|---|---|---|---|
| Plate10 | 16S rDNA sequencing | 3-OxoLCA production (%) | isoLCA production (%) | LCA production (%) | isoLCA production (%) |
| P10-A1 | Collinsella intestinalis | 34.13153 | 0.04609 | 91.13490 | 0.10915 |
| P10-A10 | | 0.04946 | 0.00000 | 0.00000 | 3.32532 |
| P10-A11 | | 0.20042 | 0.00000 | 3.45043 | 43.48243 |
| P10-A12 | | 0.05607 | 0.00000 | 3.29957 | 48.86068 |
| P10-A2 | Monoglobus pectinilyticus | 2.98334 | 0.00000 | 0.22340 | 0.00000 |
| P10-A3 | Monoglobus pectinilyticus | 2.87063 | 0.00000 | 0.04888 | 0.00000 |
| P10-A4 | Collinsella intestinalis | 27.27120 | 0.09959 | 91.05681 | 0.00000 |
| P10-A5 | | 0.09288 | 0.12955 | 94.76505 | 0.44230 |
| P10-A6 | | 0.97173 | 0.00000 | 0.20248 | 0.00000 |
| P10-A7 | | 0.09553 | 0.13916 | 0.20765 | 0.00000 |
| P10-A8 | | 0.13614 | 0.00000 | 0.19039 | 0.00000 |
| P10-A9 | Peptoniphilus harei | 20.94046 | 0.06083 | 70.95821 | 17.56274 |
| P10-B1 | Monoglobus pectinilyticus | 2.72237 | 0.00000 | 0.10888 | 0.00000 |
| P10-B10 | | 0.63316 | 0.30146 | 0.44183 | 0.00000 |
| P10-B11 | | 0.53360 | 0.24133 | 85.19406 | 0.17345 |
| P10-B12 | | 0.37665 | 0.00000 | 91.23036 | 0.32686 |
| P10-B2 | | 0.14864 | 0.00000 | 0.00000 | 0.00000 |
| P10-B3 | Collinsella intestinalis | 33.52895 | 0.00000 | 87.06503 | 0.00000 |
| P10-B4 | Monoglobus pectinilyticus | 1.48688 | 0.00000 | 0.00000 | 0.00000 |
| P10-B5 | | 0.15448 | 0.00000 | 0.03048 | 0.00000 |
| P10-B6 | | 0.10106 | 0.18386 | 86.38025 | 0.17592 |
| P10-B7 | Collinsella intestinalis | 26.11192 | 0.00000 | 86.18871 | 0.00000 |
| P10-B8 | Collinsella intestinalis | 16.11306 | 0.88903 | 86.27787 | 0.00000 |
| P10-B9 | | 0.13231 | 0.00000 | 0.13349 | 3.05035 |
| P10-C1 | | 0.08269 | 0.09747 | 0.04869 | 0.32269 |
| P10-C2 | Collinsella intestinalis | 8.37484 | 0.16038 | 87.09328 | 0.27866 |
| P10-C3 | Collinsella intestinalis | 6.84692 | 0.00000 | 89.59874 | 0.00000 |
| P10-C4 | Collinsella intestinalis | 29.96992 | 0.17135 | 90.76710 | 0.00000 |
| P10-C5 | | 0.16509 | 0.00000 | 0.04407 | 0.00000 |
| P10-C6 | | 0.07534 | 0.00000 | 0.00000 | 0.00000 |

Table S2 - PLATE 11

### Sequence Summary Report

| Plate11 | 16S rDNA sequencing | LCA as substrate | | 3-OxoLCA as substrate | |
|---|---|---|---|---|---|
| | | 3-OxoLCA production (%) | isoLCA production (%) | LCA production (%) | isoLCA production (%) |
| P11-A1 | | 0.00000 | 0.11066 | 3.66193 | 45.95645 |
| P11-A10 | | 0.86539 | 0.00000 | 42.50473 | 0.13183 |
| P11-A11 | Collinsella intestinalis | 26.97747 | 0.00000 | 83.97298 | 0.23300 |
| P11-A12 | Collinsella intestinalis | 46.30576 | 0.04713 | 78.00327 | 0.44037 |
| P11-A2 | | 0.06262 | 0.10561 | 3.09950 | 37.90957 |
| P11-A3 | Collinsella intestinalis | 17.18537 | 0.06339 | 51.39268 | 0.00000 |
| P11-A4 | | 0.09696 | 0.16847 | 6.02485 | 0.00000 |
| P11-A5 | | 0.00000 | 0.12610 | 4.74418 | 32.36729 |
| P11-A6 | Collinsella intestinalis | 9.60761 | 0.22744 | 89.22872 | 0.32642 |
| P11-A7 | | 0.10321 | 0.29490 | 3.10186 | 39.14505 |
| P11-A8 | Peptoniphilus harei | 0.22200 | 0.41791 | 2.60061 | 50.62676 |
| P11-A9 | | 0.00000 | 0.10367 | 3.28555 | 42.01524 |
| P11-B1 | Bacteroides vulgatus | 0.00000 | 0.11906 | 0.31535 | 2.95486 |
| P11-B10 | Peptoniphilus harei | 28.46166 | 2.75273 | 69.66825 | 21.85608 |
| P11-B11 | | 0.07911 | 0.00000 | 2.92147 | 42.12848 |
| P11-B12 | | 0.00000 | 0.14852 | 3.10709 | 41.23396 |
| P11-B2 | Monoglobus pectinilyticus | 4.09840 | 0.00000 | 0.37248 | 0.00000 |
| P11-B3 | Collinsella intestinalis | 42.75195 | 0.07456 | 78.54120 | 0.10154 |
| P11-B4 | | 0.79552 | 0.00000 | 0.10954 | 0.00000 |
| P11-B5 | | 0.06712 | 0.00000 | 0.00000 | 0.00000 |
| P11-B6 | | 0.30473 | 0.14799 | 0.00000 | 0.00000 |
| P11-B7 | | 0.11328 | 0.20032 | 3.13469 | 36.07062 |
| P11-B8 | Collinsella intestinalis | 27.71460 | 0.08200 | 78.19122 | 0.00000 |
| P11-B9 | | 0.13525 | 0.43195 | 3.38364 | 41.91333 |
| P11-C1 | | 0.05841 | 0.00000 | 91.66464 | 0.41719 |
| P11-C10 | | 0.08099 | 0.00000 | 0.17327 | 2.06836 |
| P11-C11 | Monoglobus pectinilyticus | 1.82339 | 0.12305 | 0.20410 | 0.00000 |
| P11-C12 | | 0.11067 | 0.24922 | 2.11105 | 28.10889 |
| P11-C2 | Collinsella intestinalis | 3.48402 | 0.10618 | 35.66891 | 0.00000 |
| P11-C3 | | 0.12280 | 0.15244 | 2.97525 | 39.33789 |
| P11-C4 | | 0.10822 | 0.24221 | 1.27308 | 0.16743 |

Table S2 - PLATE 11 (continued)

#### Sequence Summary Report

| | | | | | |
|---|---|---|---|---|---|
| P11-C5 | Monoglobus pectinilyticus | 1.19860 | 0.00000 | 0.31154 | 0.08835 |
| P11-C6 | | 0.00000 | 0.00000 | 0.00000 | 0.00000 |
| P11-C7 | | 0.00000 | 0.00000 | 2.68781 | 34.09251 |
| P11-C8 | Adlercreutzia equolifaciens | 39.81859 | 0.19673 | 14.08318 | 0.00000 |
| P11-C9 | Collinsella intestinalis | 13.59109 | 0.00000 | 72.37848 | 0.00000 |
| P11-D1 | | 0.06340 | 0.10766 | 2.83116 | 44.82826 |
| P11-D10 | Collinsella intestinalis | 43.42578 | 0.23388 | 66.25658 | 0.17113 |
| P11-D11 | Collinsella intestinalis | 3.57592 | 0.00000 | 81.78846 | 0.00000 |
| P11-D12 | | 0.07987 | 0.00000 | 0.06574 | 0.00000 |
| P11-D2 | Collinsella intestinalis | 8.15272 | 0.16623 | 79.53094 | 0.00000 |
| P11-D3 | ? - no sequence result | 0.00000 | 0.15700 | 3.81178 | 48.91274 |
| P11-D4 | | 0.27014 | 0.21548 | 3.09270 | 42.12123 |
| P11-D5 | Collinsella intestinalis | 13.96320 | 0.07954 | 86.93987 | 0.00000 |
| P11-D6 | | 0.07203 | 0.13140 | 0.00000 | 0.00000 |
| P11-D7 | | 0.00000 | 0.00000 | 87.14289 | 0.00000 |
| P11-D8 | Monoglobus pectinilyticus | 1.20169 | 0.10486 | 0.06072 | 0.00000 |
| P11-D9 | Collinsella intestinalis | 5.64109 | 0.13079 | 85.44712 | 0.00000 |
| P11-E1 | Monoglobus pectinilyticus | 3.17484 | 0.23541 | 0.08938 | 0.00000 |
| P11-E10 | | 0.11794 | 0.20400 | 94.28662 | 0.21217 |
| P11-E11 | | 0.24904 | 0.24776 | 0.07438 | 0.00000 |
| P11-E12 | Collinsella intestinalis | 38.93470 | 0.07375 | 89.41156 | 0.25841 |
| P11-E2 | | 0.14723 | 0.00000 | 0.00000 | 0.00000 |
| P11-E3 | | 0.00000 | 0.00000 | 0.00000 | 0.00000 |
| P11-E4 | Peptoniphilus harei | 1.22833 | 0.14916 | 3.19996 | 38.01712 |
| P11-E5 | | 0.19518 | 0.13251 | 0.25903 | 0.00000 |
| P11-E6 | Eggerthella lenta | 61.51247 | 4.68565 | 44.52899 | 13.35614 |
| P11-E7 | | 0.08293 | 0.08112 | 84.96732 | 0.00000 |
| P11-E8 | Monoglobus pectinilyticus | 1.92583 | 0.00000 | 0.20726 | 0.00000 |
| P11-E9 | | 0.09064 | 0.00000 | 2.91248 | 32.61786 |
| P11-F1 | Monoglobus pectinilyticus | 3.48797 | 0.00000 | 0.16052 | 0.00000 |
| P11-F10 | | 0.98483 | 0.00000 | 2.42459 | 33.55051 |
| P11-F11 | Collinsella intestinalis | 19.95551 | 0.00000 | 91.59552 | 0.00000 |
| P11-F12 | Monoglobus pectinilyticus | 2.36716 | 0.00000 | 1.61500 | 0.18970 |
| P11-F2 | | 0.05987 | 0.00000 | 4.59756 | 48.34521 |
| P11-F3 | | 0.00000 | 0.23510 | 2.92159 | 40.44338 |
| P11-F4 | | 0.06379 | 0.00000 | 3.68128 | 46.00239 |
| P11-F5 | | 0.07273 | 0.00000 | 0.00000 | 0.00000 |

Table S2 - PLATE 11 (continued)

#### Sequence Summary Report

| | | | | | |
|---|---|---|---|---|---|
| P11-F6 | | 0.14416 | 0.00000 | 2.61786 | 29.49851 |
| P11-F7 | Peptoniphilus harei | 0.00000 | 0.09166 | 4.16206 | 54.03078 |
| P11-F8 | | 0.00000 | 0.11763 | 3.84235 | 43.52608 |
| P11-F9 | | 0.13621 | 0.12565 | 3.17480 | 38.36444 |
| P11-G1 | | 0.21824 | 0.39175 | 0.00000 | 0.00000 |
| P11-G10 | | 0.07388 | 0.15293 | 0.69846 | 0.13031 |
| P11-G11 | Monoglobus pectinilyticus | 1.03655 | 0.00000 | 0.36070 | 0.00000 |
| P11-G12 | | 0.27244 | 0.15139 | 90.90176 | 0.27351 |
| P11-G2 | | 0.06302 | 0.00000 | 3.34763 | 37.33372 |
| P11-G3 | | 0.00000 | 0.13599 | 1.40228 | 37.46754 |
| P11-G4 | Collinsella intestinalis | 19.45308 | 0.00000 | 83.18162 | 0.37567 |
| P11-G5 | | 0.00000 | 0.00000 | 2.70056 | 38.32069 |
| P11-G6 | | 0.06538 | 0.00000 | 0.00000 | 0.00000 |
| P11-G7 | | 0.58927 | 0.00000 | 0.08500 | 0.00000 |
| P11-G8 | | 0.71497 | 0.14668 | 0.29177 | 0.00000 |
| P11-G9 | | 0.06738 | 0.45770 | 0.19571 | 3.77796 |
| P11-H1 | | 0.06181 | 0.12437 | 89.19152 | 0.23390 |
| P11-H10 | Collinsella intestinalis | 12.48378 | 0.00000 | 89.60910 | 0.00000 |
| P11-H11 | | 0.98537 | 0.00000 | 0.88820 | 0.00000 |
| P11-H12 | Collinsella intestinalis | 2.20309 | 0.19412 | 89.71500 | 0.00000 |
| P11-H2 | | 0.00000 | 0.12022 | 4.13325 | 45.90040 |
| P11-H3 | Collinsella intestinalis | 26.38131 | 0.03697 | 89.88760 | 0.09504 |
| P11-H4 | | 0.05660 | 0.23356 | 0.00000 | 0.00000 |
| P11-H5 | | 0.23857 | 0.46837 | 2.85276 | 33.48213 |
| P11-H6 | | 0.00000 | 0.16256 | 1.52549 | 36.50106 |
| P11-H7 | | 0.25774 | 0.00000 | 89.80009 | 0.59266 |
| P11-H8 | | 0.89396 | 0.18908 | 0.11399 | 0.00000 |
| P11-H9 | | 0.00000 | 0.00000 | 3.89692 | 43.50304 |

Table S2 - biological triplicate tubes_3-oxoLCA producers

| With 100uM LCA as substrate for 48hr | | 3-oxoLCA % | | | isoLCA % | | |
|---|---|---|---|---|---|---|---|
| Human isolates | 16S Sequencing hits | Replicate1 | Replicate2 | Replicate3 | Replicate1 | Replicate2 | Replicate3 |
| P4-B10 | Phocea massiliensis | 2.36 | 2.47 | 1.65 | 0.22 | 0.51 | 0.51 |
| P4-F8 | Clostridium aldenense | 0.84 | 1.39 | 1.63 | 5.10 | 3.84 | 3.76 |
| P4-G2 | Ruminococcus gnavus | 1.15 | 0.99 | 1.20 | 22.36 | 26.97 | 32.91 |
| P4-G5 | Clostridium perfringens | 6.05 | 7.17 | 6.61 | 0.21 | 0.35 | 0.18 |
| P7-A2 | Raoultibacter massiliensis | 47.37 | 41.59 | 44.49 | 0.00 | 0.00 | 0.00 |
| P7-E3 | Gordonibacter pamelaeae | 65.42 | 69.60 | 69.47 | 6.44 | 4.73 | 5.25 |
| P7-G1 | Collinsella intestinalis | 5.12 | 4.26 | 4.49 | 0.00 | 0.00 | 0.00 |
| P7-G7 | Eggerthella lenta | 62.89 | 62.41 | 62.20 | 6.13 | 6.41 | 6.04 |
| P8-B9 | Monoglobus pectinilyticus | 4.14 | 4.44 | 3.36 | 0.60 | 0.00 | 0.00 |
| P8-C12 | Collinsella intestinalis | 22.49 | 20.35 | 24.73 | 0.00 | 0.00 | 0.19 |
| P8-F11 | Peptoniphilus harei | 0.25 | 0.19 | 0.00 | 0.00 | 0.23 | 0.00 |
| P2-D12 | Phocea massiliensis | 0.54 | 0.39 | 0.67 | 0.98 | 0.10 | 0.81 |
| P3-E2 | Clostridium perfringens | 3.27 | 4.38 | 4.41 | 0.00 | 0.23 | 0.20 |
| P5-A1 | Collinsella aerofaciens | 1.26 | 1.20 | 0.97 | 0.00 | 0.15 | 0.17 |
| P5-B9 | Bifidobacterium longum | 1.42 | 1.65 | 1.30 | 0.18 | 0.17 | 0.12 |
| P10-A9 | Peptoniphilus harei | 0.70 | 0.98 | 0.87 | 4.92 | 5.41 | 4.87 |
| P11-C8 | Adlercreutzia equolifaciens | 18.21 | 15.34 | 16.36 | 0.00 | 0.26 | 0.12 |
| P11-E6 | Eggerthella lenta | 19.75 | 18.69 | 19.28 | 34.56 | 35.86 | 36.81 |
| P10-A9 | Peptoniphilus harei | 0.77 | 1.22 | 1.66 | 2.80 | 1.52 | 2.89 |
| P11-A12 | Collinsella intestinalis | 2.05 | 3.04 | 2.63 | 0.13 | 0.00 | 0.00 |
| P11-B10 | Peptoniphilus harei | 0.00 | 0.00 | 0.00 | 0.18 | 0.00 | 0.14 |
| P4-G12 | Clostridium perfringens | 2.03 | 1.42 | 1.56 | 1.03 | 0.00 | 0.00 |
| P8-C1 | Monoglobus pectinilyticus | 0.29 | 0.24 | 0.27 | 0.00 | 0.00 | 0.00 |
| P8-C10 | Collinsella intestinalis | 0.82 | 1.36 | 1.52 | 0.00 | 0.00 | 0.00 |
| P10-B8 | Collinsella intestinalis | 3.66 | 3.67 | 3.21 | 0.00 | 0.00 | 0.00 |
| P11-E4 | Peptoniphilus harei | 0.00 | 0.07 | 0.00 | 0.00 | 0.00 | 0.00 |
| P2-B6 | Clostridium citroniae | 3.75 | 3.70 | 3.58 | 9.50 | 7.53 | 8.96 |
| P3-G11 | Clostridium perfringens | 1.89 | 2.23 | 2.45 | 0.00 | 0.00 | 0.00 |
| P5-C2 | Collinsella aerofaciens | 1.40 | 1.35 | 1.05 | 2.00 | 2.84 | 2.17 |
| P5-E12 | Bifidobacterium pseudocatenulatum | 0.00 | 0.11 | 0.00 | 0.00 | 0.00 | 0.00 |
| P5-G3 | Bifidobacterium longum | 1.82 | 1.91 | 1.62 | 0.21 | 0.03 | 0.10 |
| P5-G6 | Bifidobacterium longum | 1.27 | 1.61 | 1.32 | 0.34 | 0.00 | 0.33 |
| P7-G9 | Eggerthella lenta | 26.68 | 26.46 | 25.06 | 2.79 | 2.93 | 2.62 |
| P8-F12 | Collinsella intestinalis | 4.60 | 4.55 | 4.10 | 0.00 | 0.00 | 0.00 |
| P4-B2 | Clostridium aldenense | 2.99 | 1.73 | 2.75 | 0.83 | 0.81 | 1.18 |

**Table S2 - biological triplicate tubes_isoLCA producers**

| With 100uM 3-oxoLCA as substrate for 48hr | | LCA % | | | isoLCA % | | |
|---|---|---|---|---|---|---|---|
| **Human isolates** | **16S Sequencing hits** | **Replicate1** | **Replicate2** | **Replicate3** | **Replicate1** | **Replicate2** | **Replicate3** |
| P4-B10 | Phocea massiliensis | 99.65 | 99.70 | 99.58 | 0.00 | 0.00 | 0.00 |
| P7-E3 | Gordonibacter pamelaeae | 13.48 | 22.46 | 15.64 | 4.36 | 8.35 | 2.71 |
| P7-G7 | Eggerthella lenta | 15.12 | 15.59 | 10.55 | 3.70 | 3.94 | 2.74 |
| P1-B12 | Bacteroides dorei | 2.70 | 2.31 | 2.35 | 6.08 | 6.55 | 5.84 |
| P1-C11 | Bacteroides vulgatus | 0.96 | 1.04 | 0.92 | 2.60 | 2.35 | 2.53 |
| P4-G3 | Clostridium perfringens | 98.93 | 98.62 | 98.75 | 0.00 | 0.00 | 0.00 |
| P5-E1 | Bacteroides dorei | 3.16 | 2.30 | 2.54 | 6.03 | 7.09 | 4.88 |
| P6-C11 | Collinsella intestinalis | 97.42 | 96.80 | 96.80 | 0.00 | 0.00 | 0.00 |
| P6-D1 | Bacteroides cellulosilyticus | 0.00 | 0.00 | 0.00 | 36.31 | 40.87 | 39.92 |
| P6-D2 | Collinsella aerofaciens | 34.30 | 31.83 | 33.76 | 5.32 | 2.88 | 4.65 |
| P6-F10 | Bacteroides uniformis | 0.00 | 0.00 | 0.00 | 47.84 | 48.70 | 44.96 |
| P6-H1 | Bacteroides fragilis | 0.98 | 0.85 | 0.70 | 44.33 | 41.04 | 38.25 |
| P8-G1 | Eggerthella lenta | 57.16 | 51.75 | 51.33 | 21.66 | 21.59 | 23.13 |
| P8-G3 | Peptoniphilus harei | 4.47 | 4.09 | 3.26 | 74.67 | 72.87 | 64.20 |
| P1-A4 | Catenibacterium mitsuokai | 0.00 | 0.00 | 0.00 | 98.01 | 95.71 | 98.98 |
| P1-D2 | Bacteroides uniformis | 0.00 | 0.00 | 0.00 | 0.00 | 0.20 | 0.00 |
| P1-D8 | Parabacteroides merdae | 0.00 | 0.00 | 0.00 | 6.23 | 7.98 | 8.26 |
| P1-F10 | Bacteroides uniformis | 0.00 | 0.00 | 0.00 | 35.50 | 36.66 | 32.20 |
| P1-F2 | Bifidobacterium pseudocatenulatum | 1.22 | 1.30 | 1.03 | 0.89 | 0.71 | 0.62 |
| P1-G2 | Bacteroides rodentium | 0.00 | 0.00 | 0.00 | 38.63 | 38.87 | 38.54 |
| P11-E4 | Peptoniphilus harei | 6.05 | 6.28 | 6.36 | 75.01 | 76.11 | 73.27 |
| P2-A2 | Lachnospira pectinoschiza | 0.00 | 0.00 | 0.00 | 96.94 | 96.27 | 95.15 |
| P2-F2 | Lactobacillus rogosae | 0.80 | 0.00 | 0.00 | 95.71 | 94.08 | 94.31 |
| P3-F8 | Ruminococcus gnavus | 71.23 | 71.01 | 71.48 | 24.89 | 24.58 | 24.11 |
| P3-G11 | Clostridium perfringens | 97.26 | 96.87 | 96.77 | 0.00 | 0.00 | 0.00 |
| P4-A7 | Bacillus coagulans | 0.00 | 0.00 | 0.00 | 81.26 | 68.18 | 51.10 |
| P4-A8 | Bacillus coagulans | 0.00 | 0.00 | 0.00 | 75.99 | 78.36 | 44.35 |
| P5-F6 | Collinsella aerofaciens | 57.82 | 54.15 | 51.63 | 6.89 | 10.96 | 10.63 |
| P5-G11 | Bacteroides cellulosilyticus | 3.82 | 5.03 | 4.02 | 58.69 | 65.18 | 60.39 |
| P6-D10 | Bacteroides vulgatus | 2.63 | 2.38 | 1.80 | 4.18 | 4.26 | 3.15 |
| P6-E4 | Collinsella aerofaciens | 99.17 | 99.25 | 99.68 | 0.00 | 0.00 | 0.00 |
| P6-H1 | Bacteroides fragilis | 2.28 | 2.46 | 3.54 | 38.88 | 39.48 | 40.84 |
| P6-H5 | Bacteroides dorei | 1.05 | 1.37 | 1.32 | 4.47 | 4.78 | 4.65 |
| P7-H12 | Peptoniphilus harei | 6.38 | 5.85 | 5.38 | 75.73 | 75.53 | 74.45 |
| P8-D2 | Eggerthella lenta | 27.52 | 30.34 | 29.29 | 13.03 | 15.32 | 12.06 |
| P8-F11 | Peptoniphilus harei | 6.00 | 5.19 | 4.93 | 82.94 | 77.27 | 78.78 |
| P8-F12 | Collinsella intestinalis | 92.64 | 94.06 | 94.99 | 0.00 | 0.00 | 0.00 |
| P9-B11 | Bifidobacterium longum | 0.00 | | 0.00 | 0.00 | | 0.00 |
| P9-D5 | Parolsenella catena | 0.00 | 0.00 | 0.00 | 0.00 | 0.00 | 0.00 |

**Table S2 - biological triplicate tubes_isoLCA producers (continued)**

| With 100uM 3-oxoLCA as substrate for 48hr | | LCA % | | | isoLCA % | | |
|---|---|---|---|---|---|---|---|
| **Human isolates** | **16S Sequencing hits** | **Replicate1** | **Replicate2** | **Replicate3** | **Replicate1** | **Replicate2** | **Replicate3** |
| P9-D9 | Parabacteroides chongii | 0.00 | 0.00 | 0.00 | 0.00 | 0.00 | 0.00 |
| P4-B2 | Clostridium aldenense | 21.90 | 21.26 | 22.06 | 6.22 | 3.14 | 6.00 |
| P1-A10 | Bacteroides fragilis | 1.03 | 0.88 | 0.88 | 31.45 | 30.82 | 30.50 |
| P4-G2 | Ruminococcus gnavus | 70.76 | 70.49 | 68.18 | 27.76 | 27.87 | 30.32 |
| C.innocuum | Clostridium innocuum DSM1286 | 0.00 | 0.00 | 0.00 | 90.07 | 89.14 | 83.95 |

*C.innocuum was predicted from bioinformatics listed in **Table S9** - Rumgna_genes_mgx

**Supplementary Table 3 | *Bacteroides fragilis* NCTC 9343 3β-HSDH homologs.** BLAST alignment of *B. fragilis* NCTC 9343 genome against known 3β-HSDHs Elen_1325 and Rumgna_00694.

| | ***E.lenta* DSM2243 3βHSDH** | | ***R.gnavus* ATCC29149 3βHSDH** | |
|---|---|---|---|---|
| **Insert name** | Percent Identity (%) | E-value | Percent Identity | E-value |
| BF2144 | 37 | 4.00E-44 | 37 | 6.00E-50 |
| BF3932 | 35 | 2.00E-39 | 31 | 2.00E-35 |
| BF0083 | 33 | 7.00E-35 | 31 | 1.00E-31 |
| BF3320 | 31 | 5.00E-34 | 30 | 7.00E-31 |
| BF1060 | 32 | 2.00E-28 | 30 | 3.00E-16 |
| BF1669 | 31 | 3.00E-24 | 31 | 2.00E-21 |
| BF0143 | 25 | 7.00E-16 | 30 | 4.00E-34 |
| BF2136 | 26 | 8.00E-09 | - | - |
| BF3538 | 28 | 8.00E-08 | - | - |
| BF2931 | 25 | 8.00E-06 | 31 | 2.00E-06 |
| BF2345 | 27 | 1.00E-05 | 28 | 4.00E-07 |

| | |
|---|---|
| Genome BLASTed | *B. fragilis* NCTC 9343 |
| Max e-value | 1.00E-02 |
| BLAST method | protein vs. protein |
| MER-FS | Assembled |
| (green) | expressed and tested |

**Supplementary table 4 | 3α/3β-HSDH homolog search in *E. lenta* isolates.** Presence / absence of Elen_0360, Elen_0690, and Elen_1325 homologs across assayed strains. Locus tags and orthologous group IDs are as reported in ElenMatchR v1.0.9003 [5].

| | | | 3α-hydroxysteroid dehydrogenase | 3β-hydroxysteroid dehydrogenase |
|---|---|---|---|---|
| **Strain Name** | **Strain Collection IDs** | **OG_ID30_COV50_2004 (Elen_0360)** | **OG_ID30_COV50_2038 (Elen_0690)** | **OG_ID30_COV50_1765 (Elen_1325)** |
| Eggerthella lenta 11C | DSM110905 | Eggerthella_lenta_11C_00554 | Eggerthella_lenta_11C_02552 | Eggerthella_lenta_11C_00083 |
| Eggerthella lenta 14A | DSM110907 | Eggerthella_lenta_14A_00561 | Eggerthella_lenta_14A_02867 | Eggerthella_lenta_14A_00168 |
| Eggerthella lenta 22C | DSM110908 | Eggerthella_lenta_22C_01352 | Eggerthella_lenta_22C_00285 | Eggerthella_lenta_22C_00914 |
| Eggerthella lenta 28B | DSM110909 | Eggerthella_lenta_28B_02480 | Not found | Eggerthella_lenta_28B_00228 |
| Eggerthella lenta A2 | DSM110911 | Eggerthella_lenta_A2_00417 | Eggerthella_lenta_A2_00758 | Homolog is split across contigs |
| Eggerthella lenta DSM 11767 | DSM11767 | Eggerthella_lenta_DSM11767_02725 | Eggerthella_lenta_DSM11767_02379 | Eggerthella_lenta_DSM11767_00025 |
| Eggerthella lenta DSM 11863 | DSM11863 | Eggerthella_lenta_DSM11863_01062 | Eggerthella_lenta_DSM11863_00011 | Eggerthella_lenta_DSM11863_00792 |
| Eggerthella lenta DSM 15644 | DSM15644,ATCC4305,CCUG34779,CIP104211 | Eggerthella_lenta_DSM15644_01907 | Not found | Eggerthella_lenta_DSM15644_00171 |
| Eggerthella lenta DSM 2243 | DSM2243,ATC25559,JCM9979,NCTC11813,VPI0255 | Eggerthella_lenta_DSM2243REF_00373 | Eggerthella_lenta_DSM2243REF_00704 | Eggerthella_lenta_DSM2243REF_01345 |
| Eggerthella lenta FAA 1-1-60AUCSF | DSM110904 | Eggerthella_lenta_1160AFAAUCSF_01952 | Eggerthella_lenta_1160AFAAUCSF_01137 | Eggerthella_lenta_1160AFAAUCSF_00507 |
| Eggerthella lenta FAA 1-3-56 | DSM110906 | Eggerthella_lenta_1356FAA_01826 | Eggerthella_lenta_1356FAA_02297 | Eggerthella_lenta_1356FAA_00478 |
| Eggerthella lenta Valencia | | Eggerthella_lenta_Valencia_02042 | Not found | Eggerthella_lenta_Valencia_00149 |
| Eggerthella sinensis DSM 16107 | DSM16107,LMG22123 | Eggerthella_sinensis_DSM16107_00168 | Eggerthella_sinensis_DSM16107_03005 | Eggerthella_sinensis_DSM16107_00947 |
| Gordonibacter pamelaeae 3C | DSM110924 | Not found | Gordonibacter_pamelaeae_3C_01808 | Gordonibacter_pamelaeae_3C_02959 |
| Gordonibacter species 28C | DSM110925 | Not found | Gordonibacter_species_28C_02314 | Gordonibacter_species_28C_01887 |

#### Supplementary Table 5a | qPCR results for 16s rRNA gene to confirm successful colonization for the second set of experiment in Fig. 4b.

| Sample # | *E.lenta* | Diet | Target | Ct values | Feces (mg) |
|---|---|---|---|---|---|
| 1 | DSM 2243 | Control | E.lenta 16s | 25.82 | 36.6 |
| 2 | DSM 2243 | Control | E.lenta 16s | 24.86 | 81.6 |
| 3 | DSM 2243 | LCA | E.lenta 16s | 24.96 | 80.0 |
| 4 | DSM 2243 | LCA | E.lenta 16s | 23.62 | 120.3 |
| 5 | DSM 2243 | LCA | E.lenta 16s | 22.99 | 60.5 |
| 6 | DSM 15644 | Control | E.lenta 16s | 25.50 | 50.8 |
| 7 | DSM 15644 | Control | E.lenta 16s | 26.21 | 65.9 |
| 8 | DSM 15644 | LCA | E.lenta 16s | 27.06 | 51.7 |
| 9 | DSM 15644 | LCA | E.lenta 16s | 25.35 | 89.7 |
| 10 | DSM 15644 | LCA | E.lenta 16s | 26.17 | 92.8 |

#### Supplementary Table 5b | qPCR results for 16s rRNA gene to confirm successful colonization for the second sets of experiment in Fig. 4c and Extended Data Fig.5.

| Sample # | *E.lenta* | 2nd bacteria | Target | Ct value | Feces (mg) |
|---|---|---|---|---|---|
| 1 | - | - | E.lenta 16s | - | 25.9 |
| 2 | - | - | E.lenta 16s | 33.63 | 23.2 |
| 3 | - | - | E.lenta 16s | 32.87 | 39.0 |
| 4 | - | - | E.lenta 16s | - | 4.0 |
| 17 | DSM 2243 | *B.fragilis* | E.lenta 16s | 22.52 | 101.5 |
| 18 | DSM 2243 | *B.fragilis* | E.lenta 16s | 24.08 | 33.3 |
| 19 | DSM 2243 | *B.fragilis* | E.lenta 16s | 23.86 | 71.5 |
| 20 | DSM 2243 | *B.fragilis* | E.lenta 16s | 24.28 | 33.6 |
| 9 | DSM 2243 | *C.innocuum* | E.lenta 16s | 23.98 | 14.6 |
| 10 | DSM 2243 | *C.innocuum* | E.lenta 16s | 24.26 | 11.3 |
| 11 | DSM 2243 | *C.innocuum* | E.lenta 16s | 22.60 | 23.3 |
| 12 | DSM 2243 | *C.innocuum* | E.lenta 16s | 23.67 | 23.6 |
| 13 | DSM 2243 | *R.gnavus* | E.lenta 16s | 22.90 | 51.5 |
| 14 | DSM 2243 | *R.gnavus* | E.lenta 16s | 22.42 | 53.3 |
| 15 | DSM 2243 | *R.gnavus* | E.lenta 16s | 22.75 | 61.7 |
| 16 | DSM 2243 | *R.gnavus* | E.lenta 16s | 23.65 | 43.5 |

| Sample # | *E.lenta* | 2nd bugs | | Ct value | Feces (mg) |
|---|---|---|---|---|---|
| 21 | - | - | E.lenta 16s | - | 26.80 |
| 22 | - | - | E.lenta 16s | - | 35.20 |
| 23 | - | - | E.lenta 16s | - | 48.50 |
| 24 | - | - | E.lenta 16s | 34.01 | 33.20 |
| 25 | DSM15644 | *B.fragilis* | E.lenta 16s | 25.91 | 48.70 |
| 26 | DSM15644 | *B.fragilis* | E.lenta 16s | 25.12 | 57.20 |
| 27 | DSM15644 | *B.fragilis* | E.lenta 16s | 25.83 | 49.90 |
| 28 | DSM15644 | *B.fragilis* | E.lenta 16s | 25.09 | 44.50 |
| 29 | DSM15644 | *C.innocuum* | E.lenta 16s | 26.12 | 28.80 |
| 30 | DSM15644 | *C.innocuum* | E.lenta 16s | 24.05 | 81.70 |
| 31 | DSM15644 | *C.innocuum* | E.lenta 16s | 24.01 | 32.70 |
| 32 | DSM15644 | *C.innocuum* | E.lenta 16s | 24.72 | 35.90 |
| 37 | DSM15644 | *R.gnavus* | E.lenta 16s | 23.29 | 48.50 |
| 38 | DSM15644 | *R.gnavus* | E.lenta 16s | 25.25 | 17.00 |
| 39 | DSM15644 | *R.gnavus* | E.lenta 16s | 24.28 | 28.60 |
| 40 | DSM15644 | *R.gnavus* | E.lenta 16s | 22.96 | 64.70 |

| Sample # | *E.lenta* | 2nd bugs | Target | Ct value | Feces (mg) |
|---|---|---|---|---|---|
| 1 | - | - | B.fragilis 16s | - | 25.9 |
| 2 | - | - | B.fragilis 16s | - | 23.2 |
| 3 | - | - | B.fragilis 16s | - | 39.0 |
| 4 | - | - | B.fragilis 16s | - | 4.0 |
| 17 | DSM 2243 | *B.fragilis* | B.fragilis 16s | 15.13 | 101.5 |
| 18 | DSM 2243 | *B.fragilis* | B.fragilis 16s | 15.04 | 33.3 |
| 19 | DSM 2243 | *B.fragilis* | B.fragilis 16s | 15.65 | 71.5 |
| 20 | DSM 2243 | *B.fragilis* | B.fragilis 16s | 16.77 | 33.6 |

| Sample # | *E.lenta* | 2nd bugs | | Ct value | Feces (mg) |
|---|---|---|---|---|---|
| 21 | - | - | B.fragilis 16s | - | 26.80 |
| 22 | - | - | B.fragilis 16s | - | 35.20 |
| 23 | - | - | B.fragilis 16s | - | 48.50 |
| 24 | - | - | B.fragilis 16s | - | 33.20 |
| 25 | DSM15644 | *B.fragilis* | B.fragilis 16s | 17.97 | 48.70 |
| 26 | DSM15644 | *B.fragilis* | B.fragilis 16s | 18.17 | 57.20 |
| 27 | DSM15644 | *B.fragilis* | B.fragilis 16s | 17.15 | 49.90 |
| 28 | DSM15644 | *B.fragilis* | B.fragilis 16s | 16.86 | 44.50 |

| Sample # | *E.lenta* | 2nd bugs | Target | Ct value | Feces (mg) |
|---|---|---|---|---|---|
| 1 | - | - | C.innocuum 16s | - | 25.9 |
| 2 | - | - | C.innocuum 16s | - | 23.2 |
| 3 | - | - | C.innocuum 16s | - | 39.0 |
| 4 | - | - | C.innocuum 16s | - | 4.0 |
| 9 | DSM 2243 | *C.innocuum* | C.innocuum 16s | 23.63 | 14.6 |
| 10 | DSM 2243 | *C.innocuum* | C.innocuum 16s | 23.62 | 11.3 |
| 11 | DSM 2243 | *C.innocuum* | C.innocuum 16s | 22.96 | 23.3 |
| 12 | DSM 2243 | *C.innocuum* | C.innocuum 16s | 26.53 | 23.6 |

| Sample # | *E.lenta* | 2nd bugs | | Ct value | Feces (mg) |
|---|---|---|---|---|---|
| 21 | - | - | C.innocuum 16s | - | 26.80 |
| 22 | - | - | C.innocuum 16s | - | 35.20 |
| 23 | - | - | C.innocuum 16s | - | 48.50 |
| 24 | - | - | C.innocuum 16s | - | 33.20 |
| 29 | DSM15644 | *C.innocuum* | C.innocuum 16s | 27.85 | 28.80 |
| 30 | DSM15644 | *C.innocuum* | C.innocuum 16s | 26.47 | 81.70 |
| 31 | DSM15644 | *C.innocuum* | C.innocuum 16s | 28.41 | 32.70 |
| 32 | DSM15644 | *C.innocuum* | C.innocuum 16s | 26.71 | 35.90 |

| Sample # | *E.lenta* | 2nd bugs | Target | Ct value | Feces (mg) |
|---|---|---|---|---|---|
| 1 | - | - | R.gnavus 16s | 35.87 | 25.9 |
| 2 | - | - | R.gnavus 16s | | 23.2 |
| 3 | - | - | R.gnavus 16s | | 39.0 |
| 4 | - | - | R.gnavus 16s | | 4.0 |
| 13 | DSM 2243 | *R.gnavus* | R.gnavus 16s | 23.57 | 51.5 |
| 14 | DSM 2243 | *R.gnavus* | R.gnavus 16s | 23.42 | 53.3 |
| 15 | DSM 2243 | *R.gnavus* | R.gnavus 16s | 23.12 | 61.7 |
| 16 | DSM 2243 | *R.gnavus* | R.gnavus 16s | 23.84 | 43.5 |

| Sample # | *E.lenta* | 2nd bugs | | Ct value | Feces (mg) |
|---|---|---|---|---|---|
| 21 | - | - | R.gnavus 16s | 36.81 | 26.80 |
| 22 | - | - | R.gnavus 16s | | 35.20 |
| 23 | - | - | R.gnavus 16s | | 48.50 |
| 24 | - | - | R.gnavus 16s | | 33.20 |
| 37 | DSM15644 | *R.gnavus* | R.gnavus 16s | 23.20 | 48.50 |
| 38 | DSM15644 | *R.gnavus* | R.gnavus 16s | 24.29 | 17.00 |
| 39 | DSM15644 | *R.gnavus* | R.gnavus 16s | 25.07 | 28.60 |
| 40 | DSM15644 | *R.gnavus* | R.gnavus 16s | 23.08 | 64.70 |

**Supplementary Table 6 | Statistical analysis of metabolite differential abundance in the PRISM and HMP2 cohorts based on linear mixed effects models.** Nominal p-values were corrected for multiple hypothesis testing using the Benjamini-Hochberg FDR method (yielding adjusted q-values).

**- PRISM**

| Compound | Compound ID | HMDB.ID | Metabolite | prevalence | RT | m.z | coefCD | coefUC | tvalCD | tvalUC | pvalCD | pvalUC | qvalCD | qvalUC |
|---|---|---|---|---|---|---|---|---|---|---|---|---|---|---|
| 12.63_373.2744m/z_C18-neg | C18-neg_Cluster_0722 | | 3-oxolithocholic acid | 0.831168831 | 12.63 | 373.2744 | -3.051117574 | -1.817156482 | -2.722486426 | -1.524136827 | 0.007297556 | 0.129714323 | 0.039196199 | 0.295497187 |
| 11.73_375.2901m/z_C18-neg | C18-neg_Cluster_0733 | | isolithocholic acid | 0.785714286 | 11.73 | 375.2901 | -2.666696574 | -2.072739042 | -2.407205363 | -1.758769868 | 0.017369009 | 0.080785981 | 0.070958679 | 0.226126972 |
| 12.42_375.2898m/z_C18-neg | C18-neg_Cluster_0731 | HMDB00761 | lithocholate | 1 | 12.42 | 375.2898 | -2.018168128 | -1.178904823 | -2.638771939 | -1.448932346 | 0.009256365 | 0.149576805 | 0.045924789 | 0.321180272 |
| 10.89_432.3114m/z_C18-neg | C18-neg_Cluster_1081 | HMDB00698 | glycolithocholate | 0.967532468 | 10.89 | 432.3114 | -2.438842055 | -1.60607758 | -2.811996622 | -1.740694225 | 0.005625708 | 0.083918663 | 0.032839297 | 0.231100616 |
| 9.51_448.3066m/z_C18-neg | C18-neg_Cluster_1196 | HMDB00631 | glycodeoxycholate | 0.974025974 | 9.51 | 448.3066 | -1.518289268 | -0.267304493 | -1.67078337 | -0.276501015 | 0.096982683 | 0.782568173 | 0.226814835 | 0.870923553 |
| 10.76_391.2852m/z_C18-neg | C18-neg_Cluster_0833 | HMDB00626 | deoxycholate | 1 | 10.76 | 391.2852 | -1.373244369 | -0.714923024 | -2.026957613 | -0.991929881 | 0.044551549 | 0.322930926 | 0.134608138 | 0.508429502 |
| 8.54_498.2893m/z_C18-neg | C18-neg_Cluster_1575 | HMDB00896 | taurodeoxycholate | 0.993506494 | 8.54 | 498.2893 | -0.321925722 | -0.080687537 | -0.339763864 | -0.08004843 | 0.734539945 | 0.936312241 | 0.841569358 | 0.964156984 |
| 9.23_391.2850m/z_C18-neg | C18-neg_Cluster_0830 | HMDB00733 | hyodeoxycholate/ursodeoxycholate | 1 | 9.23 | 391.285 | -0.198749585 | -0.582990758 | -0.305481759 | -0.842297618 | 0.760449778 | 0.401047815 | 0.859151544 | 0.582609837 |
| 9.70_482.2939m/z_C18-neg | C18-neg_Cluster_1465 | HMDB00722 | taurolithocholate | 0.941558442 | 9.7 | 482.2939 | 0.234693469 | 0.917648263 | 0.215130141 | 0.790681092 | 0.829976711 | 0.430458006 | 0.902423 | 0.608449872 |
| 8.08_448.3067m/z_C18-neg | C18-neg_Cluster_1197 | HMDB00708 | glycoursodeoxycholate | 0.948051948 | 8.08 | 448.3067 | 0.173135415 | 0.282975358 | 0.144345529 | 0.22176392 | 0.885433739 | 0.824818531 | 0.93583355 | 0.89804924 |
| 8.29_498.2890m/z_C18-neg | C18-neg_Cluster_1574 | HMDB00951 | taurochenodesoxycholate | 0.974025974 | 8.29 | 498.289 | 1.105655356 | 0.630752329 | 1.191988784 | 0.639199029 | 0.235267983 | 0.523730529 | 0.410622957 | 0.684716258 |
| 9.26_448.3065m/z_C18-neg | | HMDB00637 | glycochenodeoxycholate | 1 | 9.26 | 448.3065 | 1.057801547 | 0.425824501 | 1.439857395 | 0.544842252 | 0.152124317 | 0.586722736 | 0.307469784 | 0.733768617 |
| 8.12_464.3014m/z_C18-neg | | HMDB00138 | glycocholate | 1 | 8.12 | 464.3014 | 1.157067104 | 0.465029948 | 1.373601015 | 0.518928864 | 0.171745595 | 0.604623864 | 0.333592809 | 0.746416383 |
| 10.55_391.2848m/z_C18-neg | C18-neg_Cluster_0832 | HMDB00518 | chenodeoxycholate | 1 | 10.55 | 391.2848 | 1.631078758 | 0.555210458 | 3.455744913 | 1.105730356 | 0.000725405 | 0.270726338 | 0.008162944 | 0.455712149 |
| 7.38_514.2839m/z_C18-neg | | HMDB00036 | taurocholate | 1 | 7.38 | 514.2839 | 1.941784942 | 0.947324535 | 2.025610707 | 0.928920378 | 0.044691327 | 0.354517876 | 0.134849206 | 0.540051832 |
| 8.10_407.2800m/z_C18-neg | C18-neg_Cluster_0927 | HMDB00506 | alpha-muricholate | 0.993506494 | 8.1 | 407.28 | 2.187165151 | 0.808464858 | 2.402343564 | 0.834717533 | 0.01759214 | 0.405288812 | 0.071594497 | 0.585766293 |
| 8.33_405.2643m/z_C18-neg | C18-neg_Cluster_0900 | HMDB00391 | ketodeoxycholate | 0.993506494 | 8.33 | 405.2643 | 2.371379351 | 0.893146022 | 2.900632585 | 1.026925659 | 0.004321786 | 0.306213958 | 0.027346976 | 0.49187273 |
| 9.11_407.2799m/z_C18-neg | C18-neg_Cluster_0930 | HMDB00619 | cholate | 1 | 9.11 | 407.2799 | 2.30877639 | 0.867590157 | 3.963581926 | 1.400056081 | 0.000116869 | 0.163693041 | 0.002283773 | 0.337880822 |
| 7.26_498.2898m/z_C18-neg | C18-neg_Cluster_1576 | HMDB00874 | taurohyodeoxycholic acid/tauroursodeoxycholic acid | 0.792207792 | 7.26 | 498.2898 | 4.603187682 | 2.124010486 | 2.924937788 | 1.268643086 | 0.004016146 | 0.206658841 | 0.026045119 | 0.387456826 |
| 9.72_391.2852m/z_C18-neg | C18-neg_Cluster_0831 | | isodeoxycholic acid | 1 | 9.72 | 391.2852 | -1.022070144 | -0.891097641 | -1.49746267 | -1.227228539 | 0.136508192 | 0.221781788 | 0.285791388 | 0.404565656 |
| 10.72_389.2697m/z_C18-neg | C18-neg_Cluster_0818 | | 3-oxodeoxycholic acid | 0.993506494 | 10.72 | 389.2697 | -0.80866379 | -0.868133869 | -1.29059009 | -1.302362488 | 0.198958117 | 0.194915806 | 0.368284884 | 0.374710369 |

#### Table S6 - HMP2

| Compound | HMDB ID | Metabolite | RT | m.z | prevalence | coefActiveNonIBD | coefActiveCD | coefActiveUC | tvalActiveNonIBD | tvalActiveCD | tvalActiveUC | pvalActiveNonIBD | pvalActiveCD | pvalActiveUC | qvalActiveNonIBD | qvalActiveCD | qvalActiveUC |
|---|---|---|---|---|---|---|---|---|---|---|---|---|---|---|---|---|---|
| C18n_QI6169 | | 3-oxolithocholic acid | 12.2 | 373.2749 | 0.97985348 | -0.60744546 | -2.086262023 | -2.482534657 | -0.774345658 | -4.035855028 | -2.912574823 | 0.439252173 | 6.69E-05 | 0.003816046 | 0.816430442 | 0.00180551 | 0.316071164 |
| C18n_QI13593 | | isoallo-lithocholic acid | 11.15 | 751.5889 | 0.509157509 | -3.136617214 | -2.793226081 | -2.807499558 | -2.214795882 | -2.962169115 | -1.824036278 | 0.02742134 | 0.00326481 | 0.069004054 | 0.331568822 | 0.028381117 | 0.668512129 |
| C18n_QI6230 | | isolithocholic acid | 11.31 | 375.2906 | 0.998168498 | -0.621043597 | -1.944397702 | -2.175999246 | -0.99048648 | -4.629538746 | -3.1925764 | 0.322624575 | 5.18E-06 | 0.001538654 | 0.749582339 | 0.000278217 | 0.251447275 |
| C18n_QI48 | HMDB00761 | lithocholate | 11.98 | 375.2905 | 1 | -0.572602791 | -1.750705103 | -1.789450074 | -1.066461615 | -4.856683353 | -3.065712017 | 0.286954329 | 1.81E-06 | 0.002341326 | 0.722670778 | 0.000127584 | 0.271011442 |
| C18n_QI55 | HMDB00698 | glycolithocholate | 10.45 | 432.3119 | 1 | -0.558529916 | -1.102496943 | -1.624733916 | -1.096732885 | -3.258664169 | -2.935665726 | 0.273516514 | 0.001229738 | 0.003549607 | 0.712486372 | 0.014567985 | 0.312254274 |
| C18n_QI57 | HMDB00631 | glycodeoxycholate | 9.08 | 448.3074 | 1 | -0.548016241 | -0.987318352 | -0.984076359 | -0.964145592 | -2.699955306 | -1.593606222 | 0.335642431 | 0.007273654 | 0.111932075 | 0.756770115 | 0.049434961 | 0.735124874 |
| C18n_QI50 | HMDB00626 | deoxycholate | 10.32 | 391.2857 | 1 | -0.22194394 | -1.045715993 | -1.54579115 | -0.499189364 | -3.544349631 | -3.199309747 | 0.617961408 | 0.000447335 | 0.001504175 | 0.894519868 | 0.007198192 | 0.250067094 |
| C18n_QI62 | HMDB00896 | taurodeoxycholate | 8.39 | 498.29 | 1 | -0.876820603 | -0.624932091 | -1.259852008 | -1.22632091 | -1.341151152 | -1.621924115 | 0.220907061 | 0.180746031 | 0.105725193 | 0.6659701 | 0.39271489 | 0.727955971 |
| C18n_QI51 | HMDB00733 | hyodeoxycholate/ursodeoxycholate | 8.85 | 391.2855 | 0.998168498 | 0.611562243 | 0.157177606 | 0.531396148 | 1.484723235 | 0.579827832 | 1.187375523 | 0.138522375 | 0.562405922 | 0.235889367 | 0.579186679 | 0.754495436 | 0.824660967 |
| C18n_QI60 | HMDB00722 | taurolithocholate | 9.65 | 482.295 | 1 | -0.825346749 | -0.626933991 | -1.316810359 | -1.396949278 | -1.601950911 | -2.051026051 | 0.163318971 | 0.110073749 | 0.041012761 | 0.609225428 | 0.292026026 | 0.592096945 |
| C18n_QI58 | HMDB00708 | glycoursodeoxycholate | 7.69 | 448.3071 | 0.994505495 | 0.590763639 | 1.387065804 | 1.587552352 | 0.872495392 | 3.163432312 | 2.158245667 | 0.38353977 | 0.001696487 | 0.031592219 | 0.785633186 | 0.018226509 | 0.553826924 |
| C18n_QI61 | HMDB00951 | taurochenodeoxycholate | 8.12 | 498.2899 | 1 | 0.534155705 | 1.946317678 | 1.241317222 | 0.774035779 | 4.390797929 | 1.655654372 | 0.439435154 | 1.50E-05 | 0.098693263 | 0.816511526 | 0.000602642 | 0.716717188 |
| C18n_QI56 | HMDB00637 | glycochenodeoxycholate | 8.84 | 448.307 | 1 | 0.77033984 | 1.325585229 | 0.591029322 | 1.46272214 | 3.912140276 | 1.032979755 | 0.144445953 | 0.000110023 | 0.302330701 | 0.586076488 | 0.002596692 | 0.855063823 |
| C18n_QI59 | HMDB00138 | glycocholate | 7.74 | 464.3021 | 1 | 0.226444234 | 1.536269431 | 1.014019547 | 0.382731019 | 4.008241791 | 1.577620284 | 0.702152869 | 7.49E-05 | 0.115561416 | 0.924570504 | 0.001957851 | 0.737744449 |
| C18n_QI49 | HMDB00518 | chenodeoxycholate | 10.12 | 391.2855 | 1 | 0.337056263 | 1.477807615 | 1.287355322 | 0.775761907 | 5.159392781 | 2.726936454 | 0.438416448 | 4.17E-07 | 0.006716349 | 0.815984292 | 4.21E-05 | 0.381383737 |
| C18n_QI65 | HMDB00036 | taurocholate | 7.18 | 514.2845 | 1 | 0.245060491 | 2.017327474 | 1.487859607 | 0.366253375 | 4.624465983 | 2.046853405 | 0.714398577 | 5.30E-06 | 0.041423224 | 0.928726976 | 0.000282505 | 0.594363113 |
| C18n_QI53 | HMDB00506 | alpha-muricholate | 7.74 | 407.2804 | 0.996336996 | 1.494679417 | 1.207564188 | 1.372751672 | 2.482738741 | 3.031087096 | 2.0984038 | 0.013508416 | 0.002619327 | 0.036589923 | 0.266327585 | 0.024364032 | 0.575925564 |
| C18n_QI52 | HMDB00391 | ketodeoxycholate | 7.97 | 405.2648 | 1 | 0.971201619 | 1.763261605 | 1.623888453 | 1.796157506 | 4.940754434 | 2.763956034 | 0.073336715 | 1.21E-06 | 0.006014417 | 0.469223823 | 9.51E-05 | 0.366183288 |
| C18n_QI54 | HMDB00619 | cholate | 8.73 | 407.2804 | 1 | 0.444236712 | 1.676168032 | 1.488917799 | 0.94303391 | 5.387931327 | 2.908828363 | 0.346317773 | 1.32E-07 | 0.003860957 | 0.763641529 | 1.80E-05 | 0.317225696 |
| C18n_QI63 | HMDB00874 | taurohyodeoxycholate/tauroursodeoxycholate | 7.06 | 498.29 | 1 | -0.101610082 | 1.551583707 | 1.265646526 | -0.172858466 | 4.007115884 | 1.981634945 | 0.862863171 | 7.52E-05 | 0.048306649 | 0.97018839 | 0.00196508 | 0.61401926 |
| C18n_QI64 | HMDB00932 | tauro-alpha-muricholate/tauro-beta-muricholate | 6.15 | 514.2844 | 0.95970696 | 0.546420194 | 2.563750376 | 2.321012033 | 0.495781415 | 3.49985466 | 1.937783584 | 0.620361586 | 0.000526015 | 0.053458322 | 0.895106098 | 0.00805811 | 0.629342019 |
| C18n_QI6769 | | isodeoxycholic acid | 9.31 | 391.2858 | 1 | 0.025287813 | -0.879409103 | -1.130266744 | 0.060391028 | -3.20380616 | -2.484472406 | 0.95187887 | 0.00148155 | 0.013443986 | 0.991905926 | 0.016592322 | 0.451248618 |
| C18n_QI6700 | | 3-oxodeoxycholic acid | 10.28 | 389.2703 | 1 | 0.129492557 | 0.314102878 | -0.642858366 | 0.37122224 | 1.363903475 | -1.69606246 | 0.710697904 | 0.173479459 | 0.090768368 | 0.927365458 | 0.383159277 | 0.705949302 |

**Supplementary Table 7 | TH17/IL-17a-related genes used in this study.** We collected TH17/IL-17α-related genes including a group of genes enriched in the IL-17 signaling pathway[6] and another group of TH17 signature genes found by Revu et al[7].

| gene | host | reference |
|---|---|---|
| CCL2 | Human | [6] |
| CCL11 | Human | [6] |
| CCL20 | Human | [6] |
| CXCL1 | Human | [6] |
| CXCL2 | Human | [6] |
| CXCL3 | Human | [6] |
| CXCL5 | Human | [6] |
| CXCL6 | Human | [6] |
| CXCL10 | Human | [6] |
| DEFB4A | Human | [6] |
| IFNG | Human | [6] |
| IL1B | Human | [6] |
| IL6 | Human | [6] |
| IL17A | Human | [6] |
| LCN2 | Human | [6] |
| MMP1 | Human | [6] |
| MMP3 | Human | [6] |
| MMP9 | Human | [6] |
| MMP13 | Human | [6] |
| MUC5AC | Human | [6] |
| TNF | Human | [6] |
| RORC | Human | [7] |
| IL17A | Human | [7] |
| IL17F | Human | [7] |
| IL26 | Human | [7] |
| GZMA | Human | [7] |
| GZMB | Human | [7] |
| CXCL13 | Human | [7] |
| CCR6 | Human | [7] |
| IL23R | Human | [7] |
| IL1R1 | Human | [7] |
| IL6R | Human | [7] |
| IL6ST | Human | [7] |
| TCF7 | Human | [7] |
| LEF1 | Human | [7] |
| BATF | Human | [7] |
| MAF | Human | [7] |
| JUN | Human | [7] |
| JUNB | Human | [7] |
| JUND | Human | [7] |
| FOS | Human | [7] |

**Table S7 (continued)**

| | | |
|---|---|---|
| FOSL2 | Human | [7] |
| BCL-XL | Human | [7] |
| BCL6 | Human | [7] |
| IL2 | Human | [7] |
| IL2RA | Human | [7] |
| OSM | Human | [7] |
| SOCS3 | Human | [7] |
| GFI1 | Human | [7] |
| FOXP3 | Human | [7] |
| CTLA4 | Human | [7] |
| LAG3 | Human | [7] |
| PD1 | Human | [7] |
| IL21 | Human | [7] |
| CXCR5 | Human | [7] |
| ICOS | Human | [7] |
| IL22 | Human | [7] |

**Supplementary Table 8 | Differential expression modeling of TH17/IL-17α-related genesin HMP2.** Table fields indicate 1) gene name; 2) condition (i.e. CD:Non-IBD and UC:Non-IBD); 3) effect size estimated from the linear model's diagnosis coefficient; 4) nominal p-values from the linear models (**Methods**); 5) adjusted p-values based on the Benjamini-Hochberg procedure with target false discovery rate (FDR) of 0.25.

| gene | condition | *effect size | *p*-value | *q*-value |
|---|---|---|---|---|
| BATF | CD_vs_nonIBD | 0.66565954 | 0.0832764 | 0.3825344 |
| BATF | UC_vs_nonIBD | 0.47592211 | 0.25372625 | 0.75758555 |
| BCL6 | CD_vs_nonIBD | 0.2382921 | 0.42731984 | 0.75283582 |
| BCL6 | UC_vs_nonIBD | -0.0618586 | 0.84998722 | 0.96238886 |
| CCL11 | CD_vs_nonIBD | 1.61334959 | 0.0049228 | 0.1225743 |
| CCL11 | UC_vs_nonIBD | 2.16071217 | 0.00067176 | 0.544394 |
| CCL2 | CD_vs_nonIBD | 1.14045415 | 0.00146368 | 0.07581674 |
| CCL2 | UC_vs_nonIBD | 0.69496116 | 0.06839691 | 0.67073598 |
| CCL20 | CD_vs_nonIBD | 1.25406623 | 0.02053657 | 0.21877253 |
| CCL20 | UC_vs_nonIBD | 0.22469294 | 0.69826699 | 0.91934478 |
| CCR6 | CD_vs_nonIBD | 0.58705187 | 0.22486062 | 0.58186804 |
| CCR6 | UC_vs_nonIBD | 0.56940436 | 0.28042483 | 0.76823466 |
| CTLA4 | CD_vs_nonIBD | 0.92062258 | 0.05297442 | 0.31656802 |
| CTLA4 | UC_vs_nonIBD | 0.72813153 | 0.15824584 | 0.71702747 |
| CXCL1 | CD_vs_nonIBD | 2.59536684 | 6.60E-05 | 0.0249417 |
| CXCL1 | UC_vs_nonIBD | 1.31474778 | 0.05240926 | 0.66118259 |
| CXCL10 | CD_vs_nonIBD | 2.11897054 | 6.72E-05 | 0.0249417 |
| CXCL10 | UC_vs_nonIBD | 0.44831128 | 0.41297844 | 0.82083204 |
| CXCL13 | CD_vs_nonIBD | -0.7948318 | 0.42144955 | 0.74813925 |
| CXCL13 | UC_vs_nonIBD | -0.727229 | 0.50024863 | 0.85092407 |
| CXCL2 | CD_vs_nonIBD | 1.97153759 | 0.00164748 | 0.07966999 |
| CXCL2 | UC_vs_nonIBD | 0.46483351 | 0.48121401 | 0.84327297 |
| CXCL3 | CD_vs_nonIBD | 2.01392878 | 0.00251582 | 0.09752794 |
| CXCL3 | UC_vs_nonIBD | 0.78341499 | 0.26733981 | 0.76421472 |
| CXCL5 | CD_vs_nonIBD | 3.50329909 | 7.12E-05 | 0.0249417 |
| CXCL5 | UC_vs_nonIBD | 1.8847253 | 0.04072925 | 0.64174526 |
| CXCL6 | CD_vs_nonIBD | 2.44953922 | 2.03E-05 | 0.01542235 |
| CXCL6 | UC_vs_nonIBD | 1.39610444 | 0.0194653 | 0.61892197 |
| CXCR5 | CD_vs_nonIBD | -0.5195156 | 0.48335088 | 0.78780195 |
| CXCR5 | UC_vs_nonIBD | -0.3038025 | 0.70701093 | 0.92221105 |
| DEFB4A | CD_vs_nonIBD | 2.49965596 | 0.0101783 | 0.16390778 |
| DEFB4A | UC_vs_nonIBD | 1.63524631 | 0.11780651 | 0.69451674 |
| FOS | CD_vs_nonIBD | 0.30656161 | 0.47545646 | 0.7834018 |
| FOS | UC_vs_nonIBD | 0.28562745 | 0.54236613 | 0.86507241 |
| FOSL2 | CD_vs_nonIBD | 0.03764232 | 0.74903629 | 0.91611769 |
| FOSL2 | UC_vs_nonIBD | -0.1049566 | 0.41497136 | 0.82126252 |
| FOXP3 | CD_vs_nonIBD | 0.88351876 | 0.04632224 | 0.30033775 |
| FOXP3 | UC_vs_nonIBD | 0.7314862 | 0.12845262 | 0.70114245 |
| GFI1 | CD_vs_nonIBD | -0.0469246 | 0.64657473 | 0.8736897 |
| GFI1 | UC_vs_nonIBD | 0.04423487 | 0.69215384 | 0.91742631 |
| GZMA | CD_vs_nonIBD | 0.41631414 | 0.14528505 | 0.48601648 |

**Table S8 (continued)**

| | | | | |
|---|---|---|---|---|
| GZMA | UC_vs_nonIBD | 0.24585154 | 0.42830192 | 0.82734642 |
| GZMB | CD_vs_nonIBD | 2.0712882 | 1.45E-05 | 0.0138762 |
| GZMB | UC_vs_nonIBD | 0.96043774 | 0.05101453 | 0.65633306 |
| ICOS | CD_vs_nonIBD | 0.67927459 | 0.07568473 | 0.36633229 |
| ICOS | UC_vs_nonIBD | 0.68415145 | 0.10073633 | 0.6926757 |
| IFNG | CD_vs_nonIBD | 2.14467674 | 0.00021539 | 0.03418664 |
| IFNG | UC_vs_nonIBD | 1.35779491 | 0.02649308 | 0.62423699 |
| IL17A | CD_vs_nonIBD | 2.5119816 | 0.00024102 | 0.03617232 |
| IL17A | UC_vs_nonIBD | 1.31400284 | 0.06745852 | 0.66897715 |
| IL17F | CD_vs_nonIBD | 1.5004707 | 0.00773224 | 0.14643648 |
| IL17F | UC_vs_nonIBD | 0.27859696 | 0.64202275 | 0.90085366 |
| IL1B | CD_vs_nonIBD | 2.43670028 | 6.88E-05 | 0.0249417 |
| IL1B | UC_vs_nonIBD | 0.73837514 | 0.2429393 | 0.75120037 |
| IL1R1 | CD_vs_nonIBD | 0.64263058 | 0.00982924 | 0.16230729 |
| IL1R1 | UC_vs_nonIBD | 0.48776957 | 0.06910566 | 0.67133823 |
| IL2 | CD_vs_nonIBD | 0.38557979 | 0.33467547 | 0.68342041 |
| IL2 | UC_vs_nonIBD | 0.39172329 | 0.36917548 | 0.80287802 |
| IL21 | CD_vs_nonIBD | 0.41709149 | 0.50180211 | 0.79823766 |
| IL21 | UC_vs_nonIBD | 0.55030783 | 0.4173966 | 0.82217495 |
| IL22 | CD_vs_nonIBD | 2.49564249 | 0.00015701 | 0.03033808 |
| IL22 | UC_vs_nonIBD | 0.91028228 | 0.18528717 | 0.73205688 |
| IL23R | CD_vs_nonIBD | -0.0800676 | 0.69881493 | 0.89620542 |
| IL23R | UC_vs_nonIBD | -0.2096058 | 0.35487654 | 0.80015574 |
| IL26 | CD_vs_nonIBD | 2.32519973 | 2.82E-05 | 0.01684496 |
| IL26 | UC_vs_nonIBD | 1.52221211 | 0.0088761 | 0.59963935 |
| IL2RA | CD_vs_nonIBD | 1.13797108 | 0.00686762 | 0.13905217 |
| IL2RA | UC_vs_nonIBD | 0.77523553 | 0.0863925 | 0.68533727 |
| IL6 | CD_vs_nonIBD | 1.92724443 | 0.00254056 | 0.09752794 |
| IL6 | UC_vs_nonIBD | 0.35941129 | 0.59396285 | 0.88580386 |
| IL6R | CD_vs_nonIBD | -0.1221817 | 0.33898405 | 0.68657973 |
| IL6R | UC_vs_nonIBD | -0.2360782 | 0.0929874 | 0.68618118 |
| IL6ST | CD_vs_nonIBD | 0.25539572 | 0.08590989 | 0.38841833 |
| IL6ST | UC_vs_nonIBD | 0.25076081 | 0.12174327 | 0.69451674 |
| JUN | CD_vs_nonIBD | -0.3127255 | 0.12800662 | 0.45920034 |
| JUN | UC_vs_nonIBD | -0.4123252 | 0.06715476 | 0.66897715 |
| JUNB | CD_vs_nonIBD | 0.142444 | 0.624793 | 0.86285372 |
| JUNB | UC_vs_nonIBD | -0.1910048 | 0.54829593 | 0.86708588 |
| JUND | CD_vs_nonIBD | -0.1804596 | 0.26474693 | 0.62220295 |
| JUND | UC_vs_nonIBD | -0.0224416 | 0.89846453 | 0.97534441 |
| LAG3 | CD_vs_nonIBD | 0.75072333 | 0.00353271 | 0.11068579 |
| LAG3 | UC_vs_nonIBD | 0.84006144 | 0.00283684 | 0.58005734 |
| LCN2 | CD_vs_nonIBD | 1.99882921 | 0.00621587 | 0.13483987 |
| LCN2 | UC_vs_nonIBD | 0.80038912 | 0.30386257 | 0.77779511 |

**Table S8 (continued)**

| | | | | |
|---|---|---|---|---|
| LEF1 | CD_vs_nonIBD | 0.21464072 | 0.56904538 | 0.83350678 |
| LEF1 | UC_vs_nonIBD | 0.47128625 | 0.25391462 | 0.7576485 |
| MAF | CD_vs_nonIBD | 0.090307 | 0.84436383 | 0.95422368 |
| MAF | UC_vs_nonIBD | 0.60468103 | 0.23110106 | 0.74722093 |
| MMP1 | CD_vs_nonIBD | 2.75410459 | 0.00074791 | 0.0589297 |
| MMP1 | UC_vs_nonIBD | 1.43742136 | 0.09586379 | 0.69047639 |
| MMP13 | CD_vs_nonIBD | 2.59811573 | 0.00049254 | 0.04894808 |
| MMP13 | UC_vs_nonIBD | 0.84126055 | 0.28121054 | 0.76831517 |
| MMP3 | CD_vs_nonIBD | 3.72374924 | 1.00E-04 | 0.02560496 |
| MMP3 | UC_vs_nonIBD | 2.19330868 | 0.02895379 | 0.62423699 |
| MMP9 | CD_vs_nonIBD | 0.51876709 | 0.22250504 | 0.57976923 |
| MMP9 | UC_vs_nonIBD | 0.9222929 | 0.04902445 | 0.65510444 |
| MUC5AC | CD_vs_nonIBD | 0.83868511 | 0.24737376 | 0.60565531 |
| MUC5AC | UC_vs_nonIBD | 0.40275613 | 0.6094913 | 0.89067144 |
| OSM | CD_vs_nonIBD | 2.88265035 | 0.00013296 | 0.02793591 |
| OSM | UC_vs_nonIBD | 1.10084449 | 0.16073828 | 0.71829205 |
| RORC | CD_vs_nonIBD | -0.5521908 | 0.00327323 | 0.1081523 |
| RORC | UC_vs_nonIBD | -0.4316474 | 0.03249 | 0.62423699 |
| SOCS3 | CD_vs_nonIBD | 2.04467339 | 2.34E-05 | 0.01616636 |
| SOCS3 | UC_vs_nonIBD | 0.93418108 | 0.0614125 | 0.66644493 |
| TCF7 | CD_vs_nonIBD | 0.14508972 | 0.6346556 | 0.86771627 |
| TCF7 | UC_vs_nonIBD | 0.47570566 | 0.15640088 | 0.71454055 |
| TNF | CD_vs_nonIBD | 0.88193266 | 0.01994997 | 0.21628315 |
| TNF | UC_vs_nonIBD | 0.54404506 | 0.18251773 | 0.72969256 |

*The effect size is represented by the estimated coefficient from the linear model. Positive value means that genes are up-regulated in IBD (CD or UC) compared to Non-IBD condition. Negetive value means down-regulated cases.

**Supplementary Table 9 | Differential abundance modeling of 3α- and 3β-HSDH homologs in HMP2.** Table fields in the first sheet indicate 1) homolog ID; 2) class (i.e. 3α-HSDH-based homolog and 3β-HSDH based homolog); 3-7) the prevalence of each homolog in each disease phenotype; 8-9) effect size estimated from the linear model's dysbiosis coefficient; 10-11) nominal p-values from the linear mixed-effects models (**Methods**); and 12-13) adjusted p-values based on the Benjamini-Hochberg procedure with target false discovery rate (FDR) of 0.05. Table fields in the 2-4 sheets follow the same format as the first sheet except the first three fields: 1) the UniRef90 annotations (which had been pre-computed using HUMAnN) (i.e. protein sequences with >90% amino acid identity and >80% coverage) of the genes identified as 3α- and/ or 3β-HSDHs; 2) the genes identified as 3α- and/ or 3β-HSDHs (Elen_1325 and Elen_0690 from E. lenta; Rumgna_02133 and Rumgna_00694 from R. gnavus; BF3538 from B. fragilis NCTC 9343); 3) the homolog of a given UniRef90 - the UniRef50 family (i.e. a set of proteins expected to have >50% identity and high coverage of the query) including all UniRef90 families belonging to it.

**- sum_mgx**

| Homolog ID | class | pre.CD-dys | pre.CD-nondys | pre.UC-dys | pre.UC-nondys | prev.nonIBD-nondys | coef.CD-dys | coef.UC-dys | pval.CD-dys | pval.UC-dys | qval.CD-dys | qval.UC-dys |
|---|---|---|---|---|---|---|---|---|---|---|---|---|
| 3aHSDH | 3aHSDH | 0.81920904 | 0.990990991 | 0.921568627 | 0.989637306 | 0.997389034 | -0.655616711 | -0.3446491 | 9.32E-25 | 7.72E-05 | 6.52E-24 | 0.000199341 |
| Rumgna_02133 homolog | 3aHSDH | 0.807909605 | 0.987387387 | 0.921568627 | 0.989637306 | 0.997389034 | -0.666843011 | -0.357198708 | 9.61E-24 | 8.54E-05 | 3.36E-23 | 0.000199341 |
| Elen_0690 homolog | 3aHSDH | 0.237288136 | 0.598198198 | 0.509803922 | 0.621761658 | 0.72845953 | -0.346298054 | -0.187409185 | 5.29E-05 | 0.107944832 | 0.000123515 | 0.151122764 |
| 3bHSDH | 3bHSDH | 0.700564972 | 0.637837838 | 0.607843137 | 0.60880829 | 0.514360313 | -0.101283607 | -0.433158446 | 0.278365728 | 0.000459439 | 0.31886309 | 0.000804018 |
| BF3538 homolog | 3bHSDH | 0.355932203 | 0.466666667 | 0.392156863 | 0.409326425 | 0.318537859 | -0.15905823 | -0.439106202 | 0.031893425 | 6.94E-06 | 0.055813494 | 4.86E-05 |
| Elen_1325 homolog | 3bHSDH | 0.333333333 | 0.174774775 | 0.215686275 | 0.186528497 | 0.161879896 | 0.108491423 | -0.040712517 | 0.056796773 | 0.600118774 | 0.079515483 | 0.600118774 |
| Rumgna_00694 homolog | 3bHSDH | 0.242937853 | 0.216216216 | 0.156862745 | 0.233160622 | 0.187989556 | 0.055924554 | -0.057733265 | 0.31886309 | 0.457082572 | 0.31886309 | 0.533263001 |

#### Table S9 - Elen_genes_mgx

| HUMAnN2 Homolog + Stratification | Query Gene ID | Query UniRef50 | pre.CD-dys | pre.CD-nondys | pre.UC-dys | pre.UC-nondys | prev.nonIBD-nondys | coef.CD-dys | coef.UC-dys | pval.CD-dys | pval.UC-dys | qval.CD-dys | qval.UC-dys |
|---|---|---|---|---|---|---|---|---|---|---|---|---|---|
| UniRef90_C4Z6X3 | ELEN_0690 | UniRef50_R5H8Q8 | 0.1469 | 0.5117 | 0.3529 | 0.5699 | 0.6710 | -0.2372 | -0.2852 | 0.0005 | 0.0021 | 0.0067 | 0.0103 |
| UniRef90_C4Z6X3\|unclassified | ELEN_0690 | UniRef50_R5H8Q8 | 0.1469 | 0.5117 | 0.3529 | 0.5699 | 0.6710 | -0.2372 | -0.2852 | 0.0005 | 0.0021 | 0.0067 | 0.0103 |
| UniRef90_R7R433 | ELEN_0690 | UniRef50_R5H8Q8 | 0.0339 | 0.0523 | 0.0588 | 0.0751 | 0.1697 | -0.0289 | -0.0040 | 0.3448 | 0.9282 | 0.7869 | 0.9953 |
| UniRef90_R7R433\|unclassified | ELEN_0690 | UniRef50_R5H8Q8 | 0.0339 | 0.0523 | 0.0588 | 0.0751 | 0.1697 | -0.0289 | -0.0040 | 0.3448 | 0.9282 | 0.7869 | 0.9953 |
| UniRef90_R5IF04 | ELEN_0690 | UniRef50_R5H8Q8 | 0.0282 | 0.0865 | 0.0784 | 0.0829 | 0.0992 | -0.0229 | 0.0442 | 0.4161 | 0.2490 | 0.7869 | 0.7782 |
| UniRef90_R5IF04\|unclassified | ELEN_0690 | UniRef50_R5H8Q8 | 0.0282 | 0.0865 | 0.0784 | 0.0829 | 0.0992 | -0.0229 | 0.0442 | 0.4161 | 0.2490 | 0.7869 | 0.7782 |
| UniRef90_C8WMP0\|unclassified | ELEN_0690 | UniRef50_R5H8Q8 | 0.0000 | 0.0018 | 0.0000 | 0.0000 | 0.0052 | -0.0012 | -0.0001 | 0.5292 | 0.9762 | 0.8268 | 0.9953 |
| UniRef90_C8WMP0\|g__Gordonibacter\|s__Gordonibacter_p | ELEN_0690 | UniRef50_R5H8Q8 | 0.0000 | 0.0000 | 0.0000 | 0.0078 | 0.0052 | -0.0011 | -0.0045 | 0.7532 | 0.4616 | 0.9953 | 0.9331 |
| UniRef90_R7BJD8 | ELEN_0690 | UniRef50_R5H8Q8 | 0.0000 | 0.0000 | 0.0588 | 0.0052 | 0.0444 | -0.0006 | 0.0791 | 0.9555 | 0.0000 | 0.9953 | 0.0000 |
| UniRef90_R7BJD8\|unclassified | ELEN_0690 | UniRef50_R5H8Q8 | 0.0000 | 0.0000 | 0.0588 | 0.0052 | 0.0444 | -0.0006 | 0.0791 | 0.9555 | 0.0000 | 0.9953 | 0.0000 |
| UniRef90_R5U0E4\|unclassified | ELEN_1325 | UniRef50_C8WGQ3 | 0.0000 | 0.0018 | 0.0000 | 0.0000 | 0.0000 | -0.0005 | 0.0000 | 0.4721 | 0.9878 | 0.7869 | 0.9953 |
| UniRef90_C8WGQ3\|unclassified | ELEN_1325 | UniRef50_C8WGQ3 | 0.0000 | 0.0018 | 0.0000 | 0.0000 | 0.0000 | -0.0005 | 0.0000 | 0.4713 | 0.9953 | 0.7869 | 0.9953 |
| UniRef90_C8WMP0\|g__Peptostreptococcaceae_noname\|s_ | ELEN_0690 | UniRef50_R5H8Q8 | 0.0000 | 0.0000 | 0.0196 | 0.0000 | 0.0000 | 0.0000 | 0.0060 | 0.9977 | 0.0000 | 0.9977 | 0.0000 |
| UniRef90_C8WGQ3\|g__Gordonibacter\|s__Gordonibacter_p | ELEN_1325 | UniRef50_C8WGQ3 | 0.0056 | 0.0000 | 0.0000 | 0.0104 | 0.0052 | 0.0005 | -0.0083 | 0.9272 | 0.3201 | 0.9953 | 0.8893 |
| UniRef90_R5H8Q8 | ELEN_0690 | UniRef50_R5H8Q8 | 0.0056 | 0.0108 | 0.0196 | 0.0181 | 0.0183 | 0.0013 | -0.0107 | 0.9005 | 0.4742 | 0.9953 | 0.9331 |
| UniRef90_R5H8Q8\|unclassified | ELEN_0690 | UniRef50_R5H8Q8 | 0.0056 | 0.0108 | 0.0196 | 0.0181 | 0.0183 | 0.0013 | -0.0107 | 0.9005 | 0.4742 | 0.9953 | 0.9331 |
| UniRef90_C8WGQ3\|g__Peptostreptococcaceae_noname\|s_ | ELEN_1325 | UniRef50_C8WGQ3 | 0.0056 | 0.0000 | 0.0196 | 0.0026 | 0.0000 | 0.0020 | 0.0072 | 0.4170 | 0.0674 | 0.7869 | 0.2807 |
| UniRef90_A6NSX2 | ELEN_0690 | UniRef50_R5H8Q8 | 0.0282 | 0.0360 | 0.0784 | 0.0492 | 0.0548 | 0.0030 | -0.0059 | 0.8865 | 0.8482 | 0.9953 | 0.9953 |
| UniRef90_A6NSX2\|g__Pseudoflavonifractor\|s__Pseudoflavo | ELEN_0690 | UniRef50_R5H8Q8 | 0.0282 | 0.0360 | 0.0784 | 0.0492 | 0.0548 | 0.0030 | -0.0059 | 0.8865 | 0.8482 | 0.9953 | 0.9953 |
| UniRef90_C8WMP0 | ELEN_0690 | UniRef50_R5H8Q8 | 0.0395 | 0.0108 | 0.0196 | 0.0078 | 0.0078 | 0.0087 | 0.0108 | 0.3970 | 0.4852 | 0.7869 | 0.9331 |
| UniRef90_C8WMP0\|g__Eggerthella\|s__Eggerthella_lenta | ELEN_0690 | UniRef50_R5H8Q8 | 0.0395 | 0.0090 | 0.0000 | 0.0000 | 0.0026 | 0.0094 | 0.0004 | 0.2318 | 0.9707 | 0.7869 | 0.9953 |
| UniRef90_C8WGQ3\|g__Eggerthella\|s__Eggerthella_lenta | ELEN_1325 | UniRef50_C8WGQ3 | 0.0508 | 0.0072 | 0.0000 | 0.0052 | 0.0026 | 0.0116 | -0.0048 | 0.0928 | 0.6292 | 0.3868 | 0.9953 |
| UniRef90_C8WGQ3 | ELEN_1325 | UniRef50_C8WGQ3 | 0.0621 | 0.0090 | 0.0196 | 0.0181 | 0.0078 | 0.0244 | -0.0088 | 0.0354 | 0.6145 | 0.2948 | 0.9953 |
| UniRef90_R5U0E4 | ELEN_1325 | UniRef50_C8WGQ3 | 0.3164 | 0.1694 | 0.2157 | 0.1813 | 0.1567 | 0.1042 | -0.0265 | 0.0632 | 0.7286 | 0.3161 | 0.9953 |
| UniRef90_R5U0E4\|g__Clostridium\|s__Clostridium_hathewa | ELEN_1325 | UniRef50_C8WGQ3 | 0.3164 | 0.1676 | 0.2157 | 0.1813 | 0.1567 | 0.1047 | -0.0268 | 0.0614 | 0.7253 | 0.3161 | 0.9953 |

### Table S9 - Rumgna_genes_mgx

| HUMAnN2 Homolog + Stratification | Query Gene ID | Query UniRef50 | pre.CD-dys | pre.CD-nondys | pre.UC-dys | pre.UC-nondys | prev.nonIBD-nondys | coef.CD-dys | coef.UC-dys | pval.CD-dys | pval.UC-dys | qval.CD-dys | qval.UC-dys |
|---|---|---|---|---|---|---|---|---|---|---|---|---|---|
| UniRef90_D4LGZ3 | Rumgna_02133 | UniRef50_R5B9W8 | 0.5989 | 0.9766 | 0.9020 | 0.9819 | 0.9922 | -0.8349 | -0.4206 | 0.0000 | 0.0000 | 0.0000 | 0.0038 |
| UniRef90_D4LGZ3\|g__Roseburia\|s__Roseburia_inulinivorans | Rumgna_02133 | UniRef50_R5B9W8 | 0.2599 | 0.5730 | 0.4706 | 0.7073 | 0.7963 | -0.7154 | -0.0522 | 0.0000 | 0.6734 | 0.0000 | 0.9993 |
| UniRef90_D4LGZ3\|g__Eubacterium\|s__Eubacterium_rectale | Rumgna_02133 | UniRef50_R5B9W8 | 0.2599 | 0.6306 | 0.3333 | 0.6218 | 0.7415 | -0.6060 | -0.4550 | 0.0000 | 0.0014 | 0.0000 | 0.0700 |
| UniRef90_D4LGZ3\|g__Blautia\|s__Ruminococcus_obeum | Rumgna_02133 | UniRef50_R5B9W8 | 0.2881 | 0.8108 | 0.7059 | 0.7953 | 0.8773 | -0.4845 | -0.2470 | 0.0000 | 0.0261 | 0.0000 | 0.2179 |
| UniRef90_D4LGZ3\|g__Dorea\|s__Dorea_formicigenerans | Rumgna_02133 | UniRef50_R5B9W8 | 0.0565 | 0.4919 | 0.4706 | 0.6528 | 0.6945 | -0.3700 | -0.2329 | 0.0000 | 0.0299 | 0.0000 | 0.2223 |
| UniRef90_D7GRT2 | Rumgna_02133 | UniRef50_R5B9W8 | 0.2542 | 0.5009 | 0.3725 | 0.5233 | 0.6136 | -0.3348 | -0.2855 | 0.0000 | 0.0096 | 0.0004 | 0.1360 |
| UniRef90_D7GRT2\|unclassified | Rumgna_02133 | UniRef50_R5B9W8 | 0.2542 | 0.5009 | 0.3725 | 0.5233 | 0.6136 | -0.3348 | -0.2855 | 0.0000 | 0.0096 | 0.0004 | 0.1360 |
| UniRef90_D4LGZ3\|g__Eubacterium\|s__Eubacterium_hallii | Rumgna_02133 | UniRef50_R5B9W8 | 0.1356 | 0.3910 | 0.2353 | 0.5440 | 0.5614 | -0.2458 | -0.2837 | 0.0006 | 0.0044 | 0.0036 | 0.1137 |
| UniRef90_D4LGZ3\|unclassified | Rumgna_02133 | UniRef50_R5B9W8 | 0.2090 | 0.4180 | 0.2549 | 0.3938 | 0.4778 | -0.2412 | -0.1718 | 0.0002 | 0.0619 | 0.0014 | 0.3576 |
| UniRef90_D4LGZ3\|g__Ruminococcus\|s__Ruminococcus_lactaris | Rumgna_02133 | UniRef50_R5B9W8 | 0.0791 | 0.1892 | 0.0392 | 0.3523 | 0.3995 | -0.1780 | -0.1159 | 0.0019 | 0.1237 | 0.0100 | 0.5362 |
| UniRef90_U2C7T4\|g__Clostridium\|s__Clostridium_sp_KLE_1755 | Rumgna_02133 | UniRef50_R5B9W8 | 0.0056 | 0.0721 | 0.0784 | 0.0466 | 0.0052 | -0.0947 | -0.0063 | 0.0064 | 0.8934 | 0.0263 | 0.9993 |
| UniRef90_A4E7X7 | Rumgna_02133 | UniRef50_R5B9W8 | 0.0565 | 0.2739 | 0.1765 | 0.3705 | 0.2924 | -0.0940 | -0.1121 | 0.0771 | 0.1165 | 0.2430 | 0.5267 |
| UniRef90_A4E7X7\|g__Collinsella\|s__Collinsella_aerofaciens | Rumgna_02133 | UniRef50_R5B9W8 | 0.0565 | 0.2703 | 0.1765 | 0.3705 | 0.2924 | -0.0865 | -0.0996 | 0.0934 | 0.1502 | 0.2775 | 0.6247 |
| UniRef90_R6BIQ6 | Rumgna_02133 | UniRef50_R5B9W8 | 0.0169 | 0.0685 | 0.0000 | 0.0104 | 0.0914 | -0.0762 | -0.0037 | 0.0051 | 0.9228 | 0.0219 | 0.9993 |
| UniRef90_R6BIQ6\|unclassified | Rumgna_02133 | UniRef50_R5B9W8 | 0.0169 | 0.0685 | 0.0000 | 0.0104 | 0.0914 | -0.0762 | -0.0037 | 0.0051 | 0.9228 | 0.0219 | 0.9993 |
| UniRef90_B9Y5C1\|g__Holdemania\|s__Holdemania_filiformis | Rumgna_00694 | UniRef50_R5TQC2 | 0.0904 | 0.1892 | 0.0588 | 0.2124 | 0.1671 | -0.0662 | -0.1525 | 0.1740 | 0.0259 | 0.4413 | 0.2179 |
| UniRef90_B9Y5C1 | Rumgna_00694 | UniRef50_R5TQC2 | 0.1017 | 0.1892 | 0.0588 | 0.2124 | 0.1697 | -0.0460 | -0.1523 | 0.3491 | 0.0272 | 0.7892 | 0.2179 |
| UniRef90_R5TKT5\|g__Lachnospiraceae_noname\|s__Lachnospiraceae_bacterium_2_1_58FAA | Rumgna_02133 | UniRef50_R5B9W8 | 0.1186 | 0.0793 | 0.1373 | 0.0777 | 0.0627 | -0.0451 | -0.0076 | 0.1976 | 0.8772 | 0.4892 | 0.9993 |
| UniRef90_D4LGZ3\|g__Ruminococcus\|s__Ruminococcus_sp_5_1_39BFAA | Rumgna_02133 | UniRef50_R5B9W8 | 0.0169 | 0.0793 | 0.0196 | 0.0777 | 0.1070 | -0.0386 | -0.1468 | 0.3862 | 0.0246 | 0.8313 | 0.2179 |
| UniRef90_C4FBF0 | Rumgna_02133 | UniRef50_R5B9W8 | 0.0226 | 0.0450 | 0.1176 | 0.0829 | 0.0183 | -0.0356 | -0.0715 | 0.1666 | 0.0396 | 0.4332 | 0.2577 |
| UniRef90_C4FBF0\|g__Collinsella\|s__Collinsella_intestinalis | Rumgna_02133 | UniRef50_R5B9W8 | 0.0169 | 0.0234 | 0.1176 | 0.0622 | 0.0104 | -0.0325 | -0.0674 | 0.0977 | 0.0105 | 0.2821 | 0.1360 |
| UniRef90_R5BAL7 | Rumgna_02133 | UniRef50_R5B9W8 | 0.0056 | 0.0252 | 0.0000 | 0.0026 | 0.0313 | -0.0295 | 0.0000 | 0.0285 | 0.9993 | 0.0927 | 0.9993 |
| UniRef90_R5BAL7\|unclassified | Rumgna_02133 | UniRef50_R5B9W8 | 0.0056 | 0.0252 | 0.0000 | 0.0026 | 0.0313 | -0.0295 | 0.0000 | 0.0285 | 0.9993 | 0.0927 | 0.9993 |
| UniRef90_U5F5L7\|g__Eubacterium\|s__Eubacterium_sp_3_1_31 | Rumgna_02133 | UniRef50_R5B9W8 | 0.0791 | 0.0613 | 0.0980 | 0.0518 | 0.0653 | -0.0237 | 0.0128 | 0.4858 | 0.7793 | 0.9294 | 0.9993 |
| UniRef90_R6WN40 | Rumgna_02133 | UniRef50_R5B9W8 | 0.0226 | 0.0306 | 0.0196 | 0.0207 | 0.0444 | -0.0229 | 0.0031 | 0.1529 | 0.8881 | 0.4078 | 0.9993 |
| UniRef90_R6WN40\|unclassified | Rumgna_02133 | UniRef50_R5B9W8 | 0.0226 | 0.0306 | 0.0196 | 0.0207 | 0.0444 | -0.0229 | 0.0031 | 0.1529 | 0.8881 | 0.4078 | 0.9993 |
| UniRef90_A4E7X7\|unclassified | Rumgna_02133 | UniRef50_R5B9W8 | 0.0000 | 0.0324 | 0.0000 | 0.0544 | 0.0209 | -0.0209 | -0.0723 | 0.2783 | 0.0075 | 0.6731 | 0.1360 |
| UniRef90_R6NVR4 | Rumgna_02133 | UniRef50_R5B9W8 | 0.0113 | 0.0378 | 0.0588 | 0.0622 | 0.0392 | -0.0114 | 0.0062 | 0.5487 | 0.8182 | 0.9746 | 0.9993 |
| UniRef90_R6NVR4\|unclassified | Rumgna_02133 | UniRef50_R5B9W8 | 0.0113 | 0.0378 | 0.0588 | 0.0622 | 0.0392 | -0.0114 | 0.0062 | 0.5487 | 0.8182 | 0.9746 | 0.9993 |
| UniRef90_S6CFR4 | Rumgna_02133 | UniRef50_R5B9W8 | 0.0113 | 0.0144 | 0.0000 | 0.0207 | 0.0809 | -0.0103 | -0.0154 | 0.5904 | 0.5870 | 0.9746 | 0.9993 |
| UniRef90_S6CFR4\|g__Adlercreutzia\|s__Adlercreutzia_equolifaciens | Rumgna_02133 | UniRef50_R5B9W8 | 0.0113 | 0.0144 | 0.0000 | 0.0207 | 0.0809 | -0.0103 | -0.0154 | 0.5904 | 0.5870 | 0.9746 | 0.9993 |
| UniRef90_E7GCZ0 | Rumgna_02133 | UniRef50_R5B9W8 | 0.0113 | 0.0198 | 0.0000 | 0.0078 | 0.0183 | -0.0074 | -0.0080 | 0.5334 | 0.6779 | 0.9746 | 0.9993 |
| UniRef90_R5QVC4 | Rumgna_02133 | UniRef50_R5B9W8 | 0.0226 | 0.0306 | 0.0392 | 0.0130 | 0.0235 | -0.0066 | 0.0031 | 0.6753 | 0.8982 | 0.9851 | 0.9993 |
| UniRef90_R5QVC4\|unclassified | Rumgna_02133 | UniRef50_R5B9W8 | 0.0226 | 0.0306 | 0.0392 | 0.0130 | 0.0235 | -0.0066 | 0.0031 | 0.6753 | 0.8982 | 0.9851 | 0.9993 |
| UniRef90_R7NJY1 | Rumgna_02133 | UniRef50_R5B9W8 | 0.0169 | 0.0198 | 0.0000 | 0.0130 | 0.0104 | -0.0048 | -0.0291 | 0.7418 | 0.2075 | 0.9851 | 0.7992 |
| UniRef90_R7NJY1\|unclassified | Rumgna_02133 | UniRef50_R5B9W8 | 0.0169 | 0.0198 | 0.0000 | 0.0130 | 0.0104 | -0.0048 | -0.0291 | 0.7418 | 0.2075 | 0.9851 | 0.7992 |
| UniRef90_E7GCZ0\|unclassified | Rumgna_02133 | UniRef50_R5B9W8 | 0.0056 | 0.0090 | 0.0000 | 0.0052 | 0.0078 | -0.0039 | -0.0048 | 0.5771 | 0.6584 | 0.9746 | 0.9993 |
| UniRef90_R9IUF6 | Rumgna_02133 | UniRef50_R5B9W8 | 0.0169 | 0.0162 | 0.0000 | 0.0104 | 0.0131 | -0.0036 | -0.0101 | 0.7316 | 0.5422 | 0.9851 | 0.9993 |
| UniRef90_R9IUF6\|unclassified | Rumgna_02133 | UniRef50_R5B9W8 | 0.0169 | 0.0162 | 0.0000 | 0.0104 | 0.0131 | -0.0036 | -0.0101 | 0.7316 | 0.5422 | 0.9851 | 0.9993 |
| UniRef90_R7AYJ2 | Rumgna_02133 | UniRef50_R5B9W8 | 0.0000 | 0.0000 | 0.0000 | 0.0233 | 0.0078 | -0.0031 | -0.0189 | 0.6814 | 0.0816 | 0.9851 | 0.4241 |
| UniRef90_R7AYJ2\|g__Erysipelotrichaceae_noname\|s__Eubacterium_cylindroides | Rumgna_02133 | UniRef50_R5B9W8 | 0.0000 | 0.0000 | 0.0000 | 0.0233 | 0.0078 | -0.0031 | -0.0189 | 0.6814 | 0.0816 | 0.9851 | 0.4241 |
| UniRef90_C4FBF0\|g__Collinsella\|s__Collinsella_tanakaei | Rumgna_02133 | UniRef50_R5B9W8 | 0.0056 | 0.0216 | 0.0000 | 0.0155 | 0.0078 | -0.0027 | -0.0005 | 0.8410 | 0.9768 | 0.9851 | 0.9993 |
| UniRef90_R5B9W8 | Rumgna_02133 | UniRef50_R5B9W8 | 0.0000 | 0.0144 | 0.0000 | 0.0000 | 0.0026 | -0.0021 | 0.0013 | 0.6974 | 0.8663 | 0.9851 | 0.9993 |
| UniRef90_R5B9W8\|unclassified | Rumgna_02133 | UniRef50_R5B9W8 | 0.0000 | 0.0144 | 0.0000 | 0.0000 | 0.0026 | -0.0021 | 0.0013 | 0.6974 | 0.8663 | 0.9851 | 0.9993 |
| UniRef90_E7GCZ0\|g__Coprobacillus\|s__Coprobacillus_sp_D6 | Rumgna_02133 | UniRef50_R5B9W8 | 0.0000 | 0.0036 | 0.0000 | 0.0000 | 0.0052 | -0.0017 | -0.0001 | 0.5999 | 0.9896 | 0.9749 | 0.9993 |
| UniRef90_E7GCZ0\|g__Coprobacillus\|s__Coprobacillus_sp_29_1 | Rumgna_02133 | UniRef50_R5B9W8 | 0.0056 | 0.0090 | 0.0000 | 0.0026 | 0.0078 | -0.0014 | -0.0020 | 0.8474 | 0.8679 | 0.9851 | 0.9993 |
| UniRef90_R7GA34 | Rumgna_02133 | UniRef50_R5B9W8 | 0.0000 | 0.0018 | 0.0000 | 0.0052 | 0.0235 | -0.0011 | -0.0016 | 0.8834 | 0.8895 | 0.9851 | 0.9993 |
| UniRef90_R7GA34\|g__Erysipelotrichaceae_noname\|s__Eubacterium_dolichum | Rumgna_02133 | UniRef50_R5B9W8 | 0.0000 | 0.0018 | 0.0000 | 0.0052 | 0.0235 | -0.0011 | -0.0016 | 0.8834 | 0.8895 | 0.9851 | 0.9993 |
| UniRef90_R7B6R4 | Rumgna_02133 | UniRef50_R5B9W8 | 0.0000 | 0.0000 | 0.0000 | 0.0000 | 0.0104 | -0.0009 | 0.0004 | 0.8845 | 0.9668 | 0.9851 | 0.9993 |
| UniRef90_R7B6R4\|unclassified | Rumgna_02133 | UniRef50_R5B9W8 | 0.0000 | 0.0000 | 0.0000 | 0.0000 | 0.0104 | -0.0009 | 0.0004 | 0.8845 | 0.9668 | 0.9851 | 0.9993 |
| UniRef90_D4LGZ3\|g__Clostridium\|s__Clostridium_sp_L2_50 | Rumgna_02133 | UniRef50_R5B9W8 | 0.0000 | 0.0054 | 0.0000 | 0.0052 | 0.0078 | -0.0007 | -0.0070 | 0.8701 | 0.2590 | 0.9851 | 0.8980 |
| UniRef90_R5CJS4 | Rumgna_02133 | UniRef50_R5B9W8 | 0.0000 | 0.0000 | 0.0000 | 0.0026 | 0.0026 | -0.0006 | -0.0005 | 0.7910 | 0.9008 | 0.9851 | 0.9993 |
| UniRef90_R5CJS4\|unclassified | Rumgna_02133 | UniRef50_R5B9W8 | 0.0000 | 0.0000 | 0.0000 | 0.0026 | 0.0026 | -0.0006 | -0.0005 | 0.7910 | 0.9008 | 0.9851 | 0.9993 |
| UniRef90_F3BB74 | Rumgna_00694 | UniRef50_R5TQC2 | 0.0000 | 0.0000 | 0.0000 | 0.0000 | 0.0104 | -0.0005 | -0.0005 | 0.9321 | 0.9549 | 0.9851 | 0.9993 |
| UniRef90_R6HKH7 | Rumgna_02133 | UniRef50_R5B9W8 | 0.0000 | 0.0000 | 0.0000 | 0.0104 | 0.0026 | -0.0005 | -0.0036 | 0.9073 | 0.5577 | 0.9851 | 0.9993 |
| UniRef90_R6HKH7\|unclassified | Rumgna_02133 | UniRef50_R5B9W8 | 0.0000 | 0.0000 | 0.0000 | 0.0104 | 0.0026 | -0.0005 | -0.0036 | 0.9073 | 0.5577 | 0.9851 | 0.9993 |
| UniRef90_F3BB74\|g__Lachnospiraceae_noname\|s__Lachnospiraceae_bacterium_2_1_46FAA | Rumgna_00694 | UniRef50_R5TQC2 | 0.0000 | 0.0000 | 0.0000 | 0.0000 | 0.0104 | -0.0003 | -0.0003 | 0.9334 | 0.9551 | 0.9851 | 0.9993 |
| UniRef90_F3BB74\|unclassified | Rumgna_00694 | UniRef50_R5TQC2 | 0.0000 | 0.0000 | 0.0000 | 0.0000 | 0.0078 | -0.0003 | -0.0002 | 0.9079 | 0.9442 | 0.9851 | 0.9993 |
| UniRef90_R5FS88 | Rumgna_02133 | UniRef50_R5B9W8 | 0.0000 | 0.0000 | 0.0000 | 0.0000 | 0.0026 | -0.0001 | 0.0001 | 0.8254 | 0.9491 | 0.9851 | 0.9993 |
| UniRef90_R5FS88\|unclassified | Rumgna_02133 | UniRef50_R5B9W8 | 0.0000 | 0.0000 | 0.0000 | 0.0000 | 0.0026 | -0.0001 | 0.0001 | 0.8254 | 0.9491 | 0.9851 | 0.9993 |
| UniRef90_R7NI08 | Rumgna_02133 | UniRef50_R5B9W8 | 0.0000 | 0.0000 | 0.0000 | 0.0026 | 0.0000 | -0.0001 | -0.0007 | 0.9404 | 0.5326 | 0.9851 | 0.9993 |
| UniRef90_R7NI08\|unclassified | Rumgna_02133 | UniRef50_R5B9W8 | 0.0000 | 0.0000 | 0.0000 | 0.0026 | 0.0000 | -0.0001 | -0.0007 | 0.9404 | 0.5326 | 0.9851 | 0.9993 |
| UniRef90_R7KB31 | Rumgna_02133 | UniRef50_R5B9W8 | 0.0000 | 0.0000 | 0.0000 | 0.0026 | 0.0000 | 0.0000 | -0.0005 | 0.9784 | 0.6575 | 0.9851 | 0.9993 |
| UniRef90_R7KB31\|unclassified | Rumgna_02133 | UniRef50_R5B9W8 | 0.0000 | 0.0000 | 0.0000 | 0.0026 | 0.0000 | 0.0000 | -0.0005 | 0.9784 | 0.6575 | 0.9851 | 0.9993 |
| UniRef90_R5TQC2 | Rumgna_00694 | UniRef50_R5TQC2 | 0.0000 | 0.0000 | 0.0000 | 0.0000 | 0.0052 | 0.0000 | 0.0000 | 0.9851 | 0.9883 | 0.9851 | 0.9993 |
| UniRef90_R5TQC2\|unclassified | Rumgna_00694 | UniRef50_R5TQC2 | 0.0000 | 0.0000 | 0.0000 | 0.0000 | 0.0052 | 0.0000 | 0.0000 | 0.9851 | 0.9883 | 0.9851 | 0.9993 |
| UniRef90_R6D564 | Rumgna_02133 | UniRef50_R5B9W8 | 0.0000 | 0.0000 | 0.0000 | 0.0000 | 0.0052 | 0.0001 | 0.0000 | 0.9498 | 0.9958 | 0.9851 | 0.9993 |
| UniRef90_R6D564\|unclassified | Rumgna_02133 | UniRef50_R5B9W8 | 0.0000 | 0.0000 | 0.0000 | 0.0000 | 0.0052 | 0.0001 | 0.0000 | 0.9498 | 0.9958 | 0.9851 | 0.9993 |
| UniRef90_C4FBF0\|unclassified | Rumgna_02133 | UniRef50_R5B9W8 | 0.0000 | 0.0000 | 0.0000 | 0.0052 | 0.0000 | 0.0002 | -0.0019 | 0.8917 | 0.4528 | 0.9851 | 0.9993 |
| UniRef90_R5TKT5\|unclassified | Rumgna_02133 | UniRef50_R5B9W8 | 0.0056 | 0.0018 | 0.0000 | 0.0000 | 0.0000 | 0.0018 | 0.0000 | 0.0915 | 0.9910 | 0.2775 | 0.9993 |
| UniRef90_R6UAF4\|g__Clostridium\|s__Clostridium_sp_HGF2 | Rumgna_00694 | UniRef50_R5TQC2 | 0.0056 | 0.0000 | 0.0000 | 0.0000 | 0.0000 | 0.0018 | 0.0000 | 0.0068 | 0.9883 | 0.0263 | 0.9993 |

#### Table S9 - Rumgna_genes_mgx (continued)

| | | | | | | | | | | | | | |
|---|---|---|---|---|---|---|---|---|---|---|---|---|---|
| UniRef90_R6UAF4\|g__Peptostreptococcaceae_noname\|s__Clostridium_difficile | Rumgna_00694 | UniRef50_R5TQC2 | 0.0056 | 0.0000 | 0.0000 | 0.0000 | 0.0000 | 0.0018 | 0.0000 | 0.0068 | 0.9883 | 0.0263 | 0.9993 |
| UniRef90_R9N7L8 | Rumgna_02133 | UniRef50_R5B9W8 | 0.0056 | 0.0072 | 0.0000 | 0.0000 | 0.0026 | 0.0024 | 0.0002 | 0.5733 | 0.9724 | 0.9746 | 0.9993 |
| UniRef90_R9N7L8\|unclassified | Rumgna_02133 | UniRef50_R5B9W8 | 0.0056 | 0.0072 | 0.0000 | 0.0000 | 0.0026 | 0.0024 | 0.0002 | 0.5733 | 0.9724 | 0.9746 | 0.9993 |
| UniRef90_R6ZBB2 | Rumgna_02133 | UniRef50_R5B9W8 | 0.0056 | 0.0018 | 0.0000 | 0.0026 | 0.0000 | 0.0028 | -0.0052 | 0.4915 | 0.4090 | 0.9294 | 0.9993 |
| UniRef90_R6ZBB2\|g__Coriobacteriaceae_noname\|s__Coriobacteriaceae_bacterium_phI | Rumgna_02133 | UniRef50_R5B9W8 | 0.0056 | 0.0018 | 0.0000 | 0.0026 | 0.0000 | 0.0028 | -0.0052 | 0.4915 | 0.4090 | 0.9294 | 0.9993 |
| UniRef90_R5TKT5\|g__Blautia\|s__Ruminococcus_gnavus | Rumgna_02133 | UniRef50_R5B9W8 | 0.4068 | 0.3009 | 0.3922 | 0.2254 | 0.1671 | 0.0032 | 0.0703 | 0.9634 | 0.4545 | 0.9851 | 0.9993 |
| UniRef90_R9K3B3 | Rumgna_02133 | UniRef50_R5B9W8 | 0.0282 | 0.0180 | 0.0000 | 0.0181 | 0.0235 | 0.0048 | -0.0164 | 0.6957 | 0.3935 | 0.9851 | 0.9993 |
| UniRef90_R9K3B3\|unclassified | Rumgna_02133 | UniRef50_R5B9W8 | 0.0282 | 0.0180 | 0.0000 | 0.0181 | 0.0235 | 0.0048 | -0.0164 | 0.6957 | 0.3935 | 0.9851 | 0.9993 |
| UniRef90_B1CB64 | Rumgna_02133 | UniRef50_R5B9W8 | 0.0056 | 0.0000 | 0.0196 | 0.0000 | 0.0000 | 0.0068 | 0.0053 | 0.0105 | 0.2275 | 0.0377 | 0.8159 |
| UniRef90_B1CB64\|g__Anaerofustis\|s__Anaerofustis_stercorihominis | Rumgna_02133 | UniRef50_R5B9W8 | 0.0056 | 0.0000 | 0.0196 | 0.0000 | 0.0000 | 0.0068 | 0.0053 | 0.0105 | 0.2275 | 0.0377 | 0.8159 |
| UniRef90_R5TKT5 | Rumgna_02133 | UniRef50_R5B9W8 | 0.4181 | 0.3045 | 0.3922 | 0.2383 | 0.1749 | 0.0083 | 0.0248 | 0.9094 | 0.8034 | 0.9851 | 0.9993 |
| UniRef90_R6UAF4\|g__Erysipelotrichaceae_noname\|s__Erysipelotrichaceae_bacterium_3_1_53 | Rumgna_00694 | UniRef50_R5TQC2 | 0.0169 | 0.0072 | 0.0000 | 0.0000 | 0.0026 | 0.0092 | -0.0003 | 0.1364 | 0.9737 | 0.3834 | 0.9993 |
| UniRef90_U2C7T4 | Rumgna_02133 | UniRef50_R5B9W8 | 0.0621 | 0.1171 | 0.0980 | 0.0829 | 0.0313 | 0.0094 | -0.0558 | 0.8397 | 0.3773 | 0.9851 | 0.9993 |
| UniRef90_R6UAF4\|unclassified | Rumgna_00694 | UniRef50_R5TQC2 | 0.0169 | 0.0000 | 0.0000 | 0.0000 | 0.0000 | 0.0098 | 0.0001 | 0.0050 | 0.9838 | 0.0219 | 0.9993 |
| UniRef90_R9MDG8 | Rumgna_02133 | UniRef50_R5B9W8 | 0.0452 | 0.0198 | 0.0000 | 0.0259 | 0.0261 | 0.0122 | -0.0173 | 0.3997 | 0.4423 | 0.8313 | 0.9993 |
| UniRef90_R9MDG8\|unclassified | Rumgna_02133 | UniRef50_R5B9W8 | 0.0452 | 0.0198 | 0.0000 | 0.0259 | 0.0261 | 0.0122 | -0.0173 | 0.3997 | 0.4423 | 0.8313 | 0.9993 |
| UniRef90_B9Y5C1\|unclassified | Rumgna_00694 | UniRef50_R5TQC2 | 0.0226 | 0.0000 | 0.0000 | 0.0000 | 0.0026 | 0.0124 | -0.0001 | 0.0000 | 0.9828 | 0.0000 | 0.9993 |
| UniRef90_R6UAF4\|g__Erysipelotrichaceae_noname\|s__Erysipelotrichaceae_bacterium_6_1_45 | Rumgna_00694 | UniRef50_R5TQC2 | 0.0678 | 0.0162 | 0.0980 | 0.0104 | 0.0183 | 0.0163 | 0.0734 | 0.3374 | 0.0020 | 0.7797 | 0.0700 |
| UniRef90_R6UAF4\|g__Erysipelotrichaceae_noname\|s__Erysipelotrichaceae_bacterium_21_3 | Rumgna_00694 | UniRef50_R5TQC2 | 0.0621 | 0.0072 | 0.0784 | 0.0104 | 0.0052 | 0.0172 | 0.0441 | 0.2879 | 0.0550 | 0.6804 | 0.3367 |
| UniRef90_R6R2D2 | Rumgna_02133 | UniRef50_R5B9W8 | 0.0621 | 0.0324 | 0.0980 | 0.0466 | 0.0627 | 0.0194 | 0.0113 | 0.4309 | 0.7628 | 0.8618 | 0.9993 |
| UniRef90_R6R2D2\|unclassified | Rumgna_02133 | UniRef50_R5B9W8 | 0.0621 | 0.0324 | 0.0980 | 0.0466 | 0.0627 | 0.0194 | 0.0113 | 0.4309 | 0.7628 | 0.8618 | 0.9993 |
| UniRef90_H7CU74 | Rumgna_02133 | UniRef50_R5B9W8 | 0.0282 | 0.0054 | 0.0196 | 0.0026 | 0.0000 | 0.0305 | 0.0209 | 0.0001 | 0.0903 | 0.0009 | 0.4267 |
| UniRef90_H7CU74\|g__Clostridium\|s__Clostridium_perfringens | Rumgna_02133 | UniRef50_R5B9W8 | 0.0282 | 0.0054 | 0.0196 | 0.0026 | 0.0000 | 0.0305 | 0.0209 | 0.0001 | 0.0903 | 0.0009 | 0.4267 |
| UniRef90_U5F5L7 | Rumgna_02133 | UniRef50_R5B9W8 | 0.0904 | 0.0631 | 0.2157 | 0.0648 | 0.0653 | 0.0322 | -0.0006 | 0.3763 | 0.9907 | 0.8313 | 0.9993 |
| UniRef90_R6UAF4\|g__Erysipelotrichaceae_noname\|s__Clostridium_innocuum | Rumgna_00694 | UniRef50_R5TQC2 | 0.0847 | 0.0072 | 0.0196 | 0.0130 | 0.0026 | 0.0336 | 0.0111 | 0.0250 | 0.5978 | 0.0866 | 0.9993 |
| UniRef90_S2XY50\|unclassified | Rumgna_02133 | UniRef50_R5B9W8 | 0.0678 | 0.0126 | 0.0196 | 0.0026 | 0.0026 | 0.0427 | 0.0070 | 0.0000 | 0.6139 | 0.0001 | 0.9993 |
| UniRef90_S2XY50\|g__Lachnospiraceae_noname\|s__Lachnospiraceae_bacterium_4_1_37FAA | Rumgna_02133 | UniRef50_R5B9W8 | 0.0452 | 0.0018 | 0.0000 | 0.0026 | 0.0026 | 0.0516 | -0.0009 | 0.0001 | 0.9651 | 0.0008 | 0.9993 |
| UniRef90_R6UAF4\|g__Erysipelotrichaceae_noname\|s__Erysipelotrichaceae_bacterium_2_2_44A | Rumgna_00694 | UniRef50_R5TQC2 | 0.0791 | 0.0108 | 0.0980 | 0.0207 | 0.0131 | 0.0562 | 0.0623 | 0.0031 | 0.0258 | 0.0152 | 0.2179 |
| UniRef90_S2XY50\|g__Lachnospiraceae_noname\|s__Lachnospiraceae_bacterium_9_1_43BFAA | Rumgna_02133 | UniRef50_R5B9W8 | 0.1130 | 0.0090 | 0.0000 | 0.0052 | 0.0026 | 0.0601 | 0.0003 | 0.0007 | 0.9911 | 0.0040 | 0.9993 |
| UniRef90_U5F5L7\|g__Erysipelotrichaceae_noname\|s__Erysipelotrichaceae_bacterium_5_2_54FAA | Rumgna_02133 | UniRef50_R5B9W8 | 0.0226 | 0.0090 | 0.1765 | 0.0233 | 0.0000 | 0.0604 | 0.0163 | 0.0001 | 0.4467 | 0.0009 | 0.9993 |
| UniRef90_U2C7T4\|unclassified | Rumgna_02133 | UniRef50_R5B9W8 | 0.0565 | 0.0450 | 0.0196 | 0.0363 | 0.0261 | 0.0832 | -0.0335 | 0.0006 | 0.3283 | 0.0036 | 0.9993 |
| UniRef90_R6UAF4 | Rumgna_00694 | UniRef50_R5TQC2 | 0.1864 | 0.0324 | 0.0980 | 0.0311 | 0.0287 | 0.1394 | 0.0821 | 0.0000 | 0.0382 | 0.0000 | 0.2577 |
| UniRef90_S2XY50 | Rumgna_02133 | UniRef50_R5B9W8 | 0.1977 | 0.0216 | 0.0196 | 0.0104 | 0.0052 | 0.1612 | 0.0108 | 0.0000 | 0.7652 | 0.0000 | 0.9993 |

#### Table S9 - BF3538_mgx

| HUMAnN2 Homolog + Stratification | Query Gene ID | UniRef50 | pre.CD-dys | pre.CD-nondys | pre.UC-dys | pre.UC-nondys | prev.nonIBD-nondys | coef.CD-dys | coef.UC-dys | pval.CD-dys | pval.UC-dys | qval.CD-dys | qval.UC-dys |
|---|---|---|---|---|---|---|---|---|---|---|---|---|---|
| UniRef90_E1WV34 | BF3538 | UniRef50_E1WV34 | 0.3107 | 0.4324 | 0.3137 | 0.3497 | 0.3185 | -0.1870 | -0.3871 | 0.0087 | 0.0000 | 0.0261 | 0.0001 |
| UniRef90_E1WV34\|g__Bacteroides\|s__Bacteroides_fragilis | BF3538 | UniRef50_E1WV34 | 0.3107 | 0.4324 | 0.3137 | 0.3446 | 0.3159 | -0.1871 | -0.3866 | 0.0084 | 0.0000 | 0.0261 | 0.0001 |
| UniRef90_E1WV34\|g__Bacteroides\|s__Bacteroides_sp_3_2_5 | BF3538 | UniRef50_E1WV34 | 0.0056 | 0.0018 | 0.0000 | 0.0000 | 0.0235 | 0.0006 | 0.0000 | 0.9485 | 0.9993 | 0.9485 | 0.9993 |
| UniRef90_E1WV34\|unclassified | BF3538 | UniRef50_E1WV34 | 0.0282 | 0.0090 | 0.0196 | 0.0155 | 0.0026 | 0.0140 | 0.0095 | 0.1781 | 0.5764 | 0.3562 | 0.6917 |
| UniRef90_R5RWD9 | BF3538 | UniRef50_E1WV34 | 0.0452 | 0.0360 | 0.0784 | 0.0596 | 0.0000 | 0.0124 | -0.0336 | 0.4799 | 0.1410 | 0.5758 | 0.2115 |
| UniRef90_R5RWD9\|g__Bacteroides\|s__Bacteroides_fragilis | BF3538 | UniRef50_E1WV34 | 0.0452 | 0.0360 | 0.0784 | 0.0596 | 0.0000 | 0.0124 | -0.0336 | 0.4799 | 0.1410 | 0.5758 | 0.2115 |

**Supplementary Table 10 | Differential abundance modeling of species with 3α-/3β-HSDH homologs in HMP2.** Table fields follow the same format as Supplementary Information Table 9 (detailed above) without the ‘class’ field.

| species | pre.CD-dys | pre.CD-nondys | pre.UC-dys | pre.UC-nondys | pre.nonIBD-nondys | coef.CD-dys | coef.UC-dys | pval.CD-dys | pval.UC-dys | qval.CD-dys | qval.UC-dys |
|---|---|---|---|---|---|---|---|---|---|---|---|
| Adlercreutzia_equolifaciens | 0.04519774 | 0.066666667 | 0.039215686 | 0.10880829 | 0.161879896 | -0.095280005 | -0.018421386 | 0.115874981 | 0.826534144 | 0.202781217 | 0.826534144 |
| Bacteroides_cellulosilyticus | 0.06779661 | 0.358558559 | 0.176470588 | 0.341968912 | 0.441253264 | -0.244964165 | -0.28521006 | 0.003337869 | 0.008327685 | 0.008761907 | 0.031264721 |
| Bacteroides_dorei | 0.412429379 | 0.666666667 | 0.62745098 | 0.652849741 | 0.796344648 | -0.741864941 | -0.181390829 | 6.43E-11 | 0.215956759 | 4.50E-10 | 0.302339463 |
| Bacteroides_fragilis | 0.457627119 | 0.645045045 | 0.549019608 | 0.533678756 | 0.527415144 | -0.61688588 | -0.761459857 | 1.50E-05 | 4.29E-05 | 7.90E-05 | 0.000450683 |
| Bacteroides_intestinalis | 0.04519774 | 0.172972973 | 0 | 0.106217617 | 0.125326371 | -0.04394252 | -0.059681073 | 0.3909779 | 0.369828381 | 0.58646685 | 0.456846823 |
| Bacteroides_uniformis | 0.598870056 | 0.936936937 | 0.607843137 | 0.914507772 | 0.934725849 | -1.199319275 | -0.673457777 | 3.65E-26 | 4.10E-06 | 7.67E-25 | 8.61E-05 |
| Bacteroides_vulgatus | 0.740112994 | 0.924324324 | 0.843137255 | 0.948186528 | 0.966057441 | -0.933533454 | -0.249291185 | 2.46E-21 | 0.048381849 | 2.58E-20 | 0.107090565 |
| Bifidobacterium_pseudocatenulatum | 0.033898305 | 0.189189189 | 0.098039216 | 0.111398964 | 0.164490862 | -0.125890139 | -0.06113882 | 0.096339207 | 0.546183713 | 0.183920305 | 0.603676736 |
| Catenibacterium_mitsuokai | 0.011299435 | 0.068468468 | 0 | 0.010362694 | 0.057441253 | -0.019473213 | -0.012038734 | 0.287458109 | 0.609420567 | 0.464355408 | 0.639891596 |
| Clostridium_bolteae | 0.774011299 | 0.747747748 | 0.784313725 | 0.79015544 | 0.537859008 | -0.238484829 | -0.338418712 | 0.074784155 | 0.057753542 | 0.157046726 | 0.110256761 |
| Clostridium_citroniae | 0.192090395 | 0.340540541 | 0.137254902 | 0.313471503 | 0.25848564 | -0.200713759 | -0.323093724 | 0.039348582 | 0.015652418 | 0.091813358 | 0.046957253 |
| Clostridium_innocuum | 0.129943503 | 0.028828829 | 0.058823529 | 0.080310881 | 0.026109661 | -0.004560498 | -0.049939383 | 0.919363197 | 0.421096189 | 0.965331357 | 0.491278888 |
| Clostridium_scindens | 0.09039548 | 0.068468468 | 0.039215686 | 0.062176166 | 0.096605744 | 0.007855126 | -0.10848664 | 0.863852421 | 0.083603735 | 0.954784255 | 0.146306536 |
| Collinsella_aerofaciens | 0.192090395 | 0.565765766 | 0.254901961 | 0.554404145 | 0.64229765 | -0.386254564 | -0.274874344 | 4.26E-05 | 0.026130731 | 0.000178956 | 0.068593168 |
| Collinsella_intestinalis | 0.04519774 | 0.043243243 | 0.117647059 | 0.090673575 | 0.028720627 | -0.108742319 | -0.125633981 | 0.003090799 | 0.008932778 | 0.008761907 | 0.031264721 |
| Eggerthella_lenta | 0.118644068 | 0.041441441 | 0 | 0.025906736 | 0.018276762 | -0.011704783 | -0.066596964 | 0.746871705 | 0.181417694 | 0.915616498 | 0.272126541 |
| Gordonibacter_pamelaeae | 0.005649718 | 0.009009009 | 0 | 0.023316062 | 0.020887728 | -0.000713535 | -0.029115134 | 0.968528731 | 0.288585251 | 0.968528731 | 0.378768142 |
| Parabacteroides_merdae | 0.214689266 | 0.630630631 | 0.215686275 | 0.668393782 | 0.618798956 | -0.392522235 | -0.350067731 | 8.23E-05 | 0.006654119 | 0.000287893 | 0.031264721 |
| Peptoniphilus_harei | 0 | 0.005405405 | 0 | 0.025906736 | 0.007832898 | -0.004679827 | -0.036509825 | 0.784814141 | 0.136508428 | 0.915616498 | 0.220513615 |
| Pseudoflavonifractor_capillosus | 0.129943503 | 0.147747748 | 0.117647059 | 0.18134715 | 0.279373368 | 0.029500081 | -0.155222993 | 0.621468627 | 0.050995507 | 0.815677573 | 0.107090565 |
| Ruminococcus_gnavus | 0.768361582 | 0.607207207 | 0.882352941 | 0.603626943 | 0.467362924 | 0.098122606 | 0.458889809 | 0.433419916 | 0.005888036 | 0.606787882 | 0.031264721 |
